# Exact expressions and numerical evaluation of average evolvability measures for characterizing and comparing G matrices

**DOI:** 10.1101/2022.11.02.514929

**Authors:** Junya Watanabe

**Affiliations:** Department of Earth Sciences, University of Cambridge, Downing Street, Cambridge, CB2 3EQ, United Kingdom

**Keywords:** evolutionary constraint, phenotypic integration, quadratic form, quantitative genetics, random skewers, zonal polynomial

## Abstract

Theory predicts that the additive genetic covariance (**G**) matrix determines a population’s short-term (in)ability to respond to directional selection—evolvability in the Hansen–Houle sense—which is typically quantified and compared via certain scalar indices called evolvability measures. Often, interest is in obtaining the averages of these measures across all possible selection gradients, but explicit formulae for most of these average measures have not been known. Previous authors relied either on approximations by the delta method, whose accuracy is generally unknown, or Monte Carlo evaluations (including the random skewers analysis), which necessarily involve random fluctuations. This study presents new, exact expressions for the average conditional evolvability, average autonomy, average respondability, average flexibility, average response difference, and average response correlation, utilizing their mathematical structures as ratios of quadratic forms. The new expressions are infinite series involving top-order zonal and invariant polynomials of matrix arguments, and can be numerically evaluated as their partial sums with, for some measures, known error bounds. Whenever these partial sums numerically converge within reasonable computational time and memory, they will replace the previous approximate methods. In addition, new expressions are derived for the average measures under a general normal distribution for the selection gradient, extending the applicability of these measures into a substantially broader class of selection regimes.

## 1 Introduction

The quantitative genetic theory of multivariate trait evolution provides a powerful framework to analyze and predict phenotypic evolution (Steppan *et al*., 2002; Blows, 2007; Blows & Walsh, 2009; Walsh & Blows, 2009; Teplitsky *et al*., 2014). At the core of the theory is the Lande equation, which describes a population’s response to directional selection under certain simplifying conditions (Lande, 1979; Lande & Arnold, 1983):

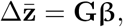

where **β** is a *p*-dimensional selection gradient vector (partial regression coefficients of trait values to relative fitness), **G** is a *p* × *p* (additive) genetic covariance matrix, and 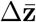 is a *p*-dimensional vector of per-generation change of the population’s mean trait values. (Throughout the paper, *p* denotes the (nominal) number of the traits/variables analyzed, and all vectors are column vectors without transposition.) In light of this theory, the **G** matrix is supposed to represent the population’s (in)ability to respond to directional selection—what has been termed evolvability by Hansen, Houle and colleagues (Houle, 1992; Hansen, 2003; Hansen *et al*., 2003a,b, 2011; Hansen & Houle, 2008; Hansen & Pélabon, 2021).^1^ The theory has motivated extensive investigations into aspects of the **G** matrix or its substitutes, ranging from theoretical and simulation-based analyses (e.g., Pavlicev & Hansen, 2011; Pavlicev *et al*., 2011; Jones *et al*., 2012; Chevin, 2013; Melo & Marroig, 2015; Hansen *et al*., 2019) to empirical inter-population/specific comparisons of trait covariation and its expansion to inter-population divergences (e.g., Cheverud, 1982, 1989; Schluter, 1996; Porto *et al*., 2009; Marroig *et al*., 2009; Rolian, 2009; Bolstad *et al*., 2014; Haber, 2016; Puentes *et al*., 2016; Costa e Silva *et al*., 2020; Machado, 2020; McGlothlin *et al*., 2022; Opedal *et al*., 2022, 2023; Hubbe *et al*., 2023).

Based on this framework, and following earlier theoretical developments (Hansen, 2003; Hansen *et al*., 2003a,b), Hansen & Houle (2008) proposed several scalar indices to capture certain aspects of the multivariate evolvability represented in **G** (see also Kirkpatrick, 2009; Marroig *et al*., 2009). Of these, (unconditional) evolvability *e*, conditional evolvability *c*, autonomy *a*, integration *i*, respondability *r*, and flexibility *f* (due to Marroig *et al*., 2009) concern a single **G** matrix (Fig. 1A), whereas response difference *d* concerns comparison between two **G** matrices. Related to the latter is the use of response correlation *ρ* for matrix comparison (Cheverud, 1996; Cheverud & Marroig, 2007) (Fig. 1B). Notably, all these indices are simple or multiple ratios of quadratic forms in **β** (see below). In this paper, they are collectively called evolvability measures. Many of them can be related to the rate of adaptation or evolutionary constraint/bias (e.g., Hansen & Houle, 2008; Chevin, 2013; Bolstad *et al*., 2014; Hansen *et al*., 2019), providing means to characterize and compare **G** matrices in biologically meaningful ways.

**Figure 1.**
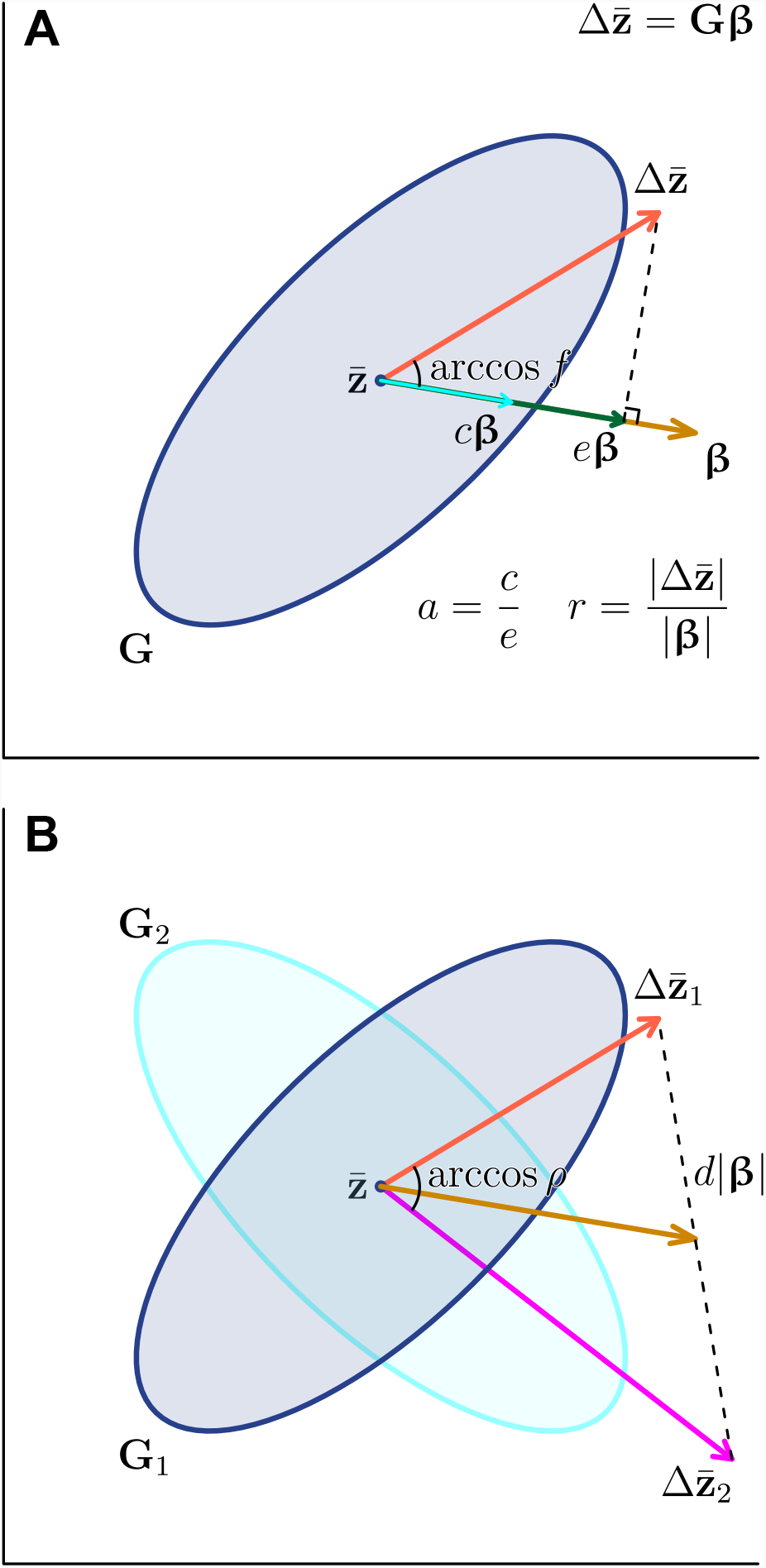
Geometric interpretations of evolvability measures. In a hypothetical bivariate trait space, genetic covariance matrices **G** are schematically represented by equiprobability ellipses around the mean phenotype 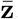, and selection gradients **β** by acute-headed brown arrows. Response vectors 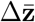 are matrix products of **G** and **β**, represented by broad-headed red/magenta arrows. **A**) One-matrix measures: evolvability *e* is the norm of 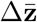 projected on **β**, standardized by the norm of the latter, |**β**|; conditional evolvability *c* is the same but when 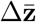 is constrained along **β**; autonomy *a* is the ratio of *c* to *e*; respondability *r* is 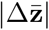 standardized by |**β**|; and flexibility *f* is the similarity in direction (cosine of the angle) between 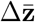 and **β. B**) Two-matrix comparison measures: response difference *d* is the distance between the end points of the two 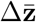’s standardized by |**β**|, and response correlation *ρ* is their similarity in direction.

The evolvability measures are primarily defined with respect to a given selection gradient **β**. In this sense, they represent aspects of **G** matrices under a fixed, deterministic selection. In many theoretical and comparative analyses of evolvability (see references above), however, interest is often in characterizing **G** matrices without referring to **β**. For this purpose, it is sensible to take the expectation of an evolvability measure assuming a certain probability distribution of **β**. Most typically, although not necessarily, the uniform distribution on a unit hypersphere is assumed for this purpose, representing a completely randomly directed and uncorrelated selection regime. The resultant quantities are called average measures hereafter.^2^ Biologically speaking, the average measures may be regarded as representations of general evolvability (in the sense of Riederer *et al*., 2022), as compared to specific evolvability which is represented by the evolvability measures as functions of a fixed **β**.

However, there is a major obstacle to using average measures for practical or even theoretical investigations. That is, most of the average measures lack known explicit, exact expressions beyond as expectations, except for the average evolvability *ē* and, under certain special conditions, the average conditional evolvability 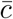 and the average respondability 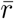 (Hansen & Houle, 2008; Kirkpatrick, 2009). Consequently, previous studies had to rely on approximations of average measures. Presumably the most widely used approximation method is Monte Carlo evaluation (e.g., Marroig *et al*., 2009; Bolstad *et al*., 2014; Haber, 2016; Grabowski & Porto, 2017), in which a large number of **β** is randomly generated on a computer and the resultant values calculated with the primary definition are literally averaged to yield an estimate of average measure. This has long been done in the so-called random skewers analysis for matrix comparison using response correlation *ρ* (e.g., Cheverud, 1996; Cheverud & Marroig, 2007; Revell, 2007; see also Rohlf, 2017). It is implemented in the `R` packages `evolvability` (Bolstad *et al*., 2014) and `evolqg` (Melo *et al*., 2016) for calculating various average measures. This method necessarily involves random fluctuations in the estimate and can take a large computational time, although the latter is becoming less of a concern with modern-day computers. On the other hand, Hansen & Houle (2008, 2009) themselves provided approximate expressions for average measures based on the delta method (which they called “analytical” or “standard” approximations). Apart from ill-defined notations used therein, a practical problem there is that the accuracy of this sort of approximation is generally unknown, so a separate Monte Carlo evaluation is usually required to ascertain whether the delta method approximation has an acceptable accuracy. This approximation has been used in some subsequent studies (e.g., Hansen & Voje, 2011; Brommer, 2014; Delahaie *et al*., 2017; Saltzberg *et al*., 2022) and is available in the `evolvability` package. Apart from these methods, Kirkpatrick (2009) used numerical integration to evaluate his average selection response, a measure equivalent to respondability (below), but this method has not been widely used in evaluating average measures.

This technical paper provides exact expressions for the following average measures: average evolvability *ē*, average conditional evolvability 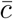, average autonomy *ā*, average respondability 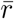, average flexibility 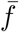, average response difference 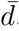, and average response correlation 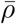. These expressions are derived from existing and new results on the moments of simple or multiple ratios of quadratic forms in normal variables. They are expressed as infinite series involving zonal or invariant polynomials of matrix arguments, and in most cases can be numerically evaluated as their partial sums to yield quasi-exact values. For some of the expressions, an upper bound for the truncation error is available. In addition to the special condition where **β** is spherically distributed as assumed in most previous studies (but see also Chevin, 2013), this study also concerns the general condition that **β** is normally distributed with potentially nonzero mean and nonspherical covariance. The latter condition can be of substantial biological interest, as it can model a fairly wide range of random directional and/or correlated selection regimes.

The paper is structured as follows. Section 2 first reviews the definition of the evolvability measures; some of them are redefined to accommodate potentially singular **G** matrices. After reviewing some known results on the moment of a ratio of quadratic forms, it then provides new expressions for average evolvability measures under the spherical distribution of **β**. Section 3 presents a new `R` implementation of the analytic results and evaluate its performance by numerical experiments. Section 4 concludes the main body of the paper by adding some theoretical and practical considerations on the use of average measures. As the zonal and invariant polynomials appear to have rarely been used in the biological literature, Appendix A gives a brief overview on their theories, providing a basis for the present mathematical results. Appendix B states a new result on the moment of a multiple ratio of quadratic forms in normal variables, and Appendix C presents new expressions for average measures under the general normal distribution of **β**. Appendix D clarifies connections to previous results derived under special conditions (Hansen & Houle, 2008; Kirkpatrick, 2009).

## 2 Theory

### 2.1 Notations

In the following discussion, it is convenient to define the linear combination of the variables (traits) along the direction of **β**. For this purpose, consider the decomposition **β** = |**β**|**u**, where |·| is the vector norm or length, 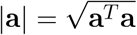 for an arbitrary vector **a** (the superscript ^*T*^ denotes matrix transpose), and **u** is a unit vector: **u**^*T*^ **u** = 1. Define the *p* × (*p* − 1) matrix **U**_(−1)_ so that the matrix **U** = (**u, U**_(−1)_) is orthogonal. With these, the orthogonal linear transform of the variables **z**\* = **U**^*T*^ **z** will be considered; the first entry of this new vector **u**^*T*^ **z** represents the score along the direction of **β**. Note that the covariance matrix of the new variables **z**\* can be written as

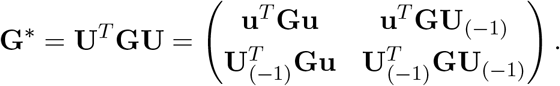

Previous authors (Hansen & Houle, 2008; Marroig *et al*., 2009; Grabowski & Porto, 2017) defined some evolvability measures under the (sometimes implicit) constraint |**β**| = 1. To avoid potential confusion, this standardized vector is always denoted by **u** herein. Throughout this paper, **G** is assumed to be given rather than random and validly constructed as a covariance matrix, that is, symmetric and nonnegative definite (i.e., either positive definite or positive semidefinite, in the terminology of Schott, 2016).

Although Hansen & Houle (2008) assumed **G** to be positive definite, this assumption is not imposed here to accommodate potentially singular (and positive semidefinite) **G** where possible. This is done by using the generalized inverse; for a matrix **A**, a generalized inverse **A**^−^ is such a matrix that satisfies **AA**^−^**A** = **A**. If **A** is nonsingular, **A**^−^ = **A**^−1^.

For notational simplicity, the range or column space and the null space of a nonzero *p* × *q* matrix **A** are defined: *R*(**A**) = {**y** : **y** = **Ax, x** ∈ ℝ^*q*^}, and *N*(**A**) = {**x** : **0**_*p*_ = **Ax, x** ∈ ℝ^*q*^}, where **0**_*p*_ is the *p*-dimensional vector of zeros. When **A** is nonnegative definite, these are the spaces spanned by the eigenvectors corresponding to its nonzero and zero eigenvalues, respectively. The *p*-dimensional identity matrix is denoted by **I**_*p*_.

### 2.2 Evolvability measures: fixed selection

Evolvability *e*(**β**) is the variance in **G** along the direction of selection **u**, or equivalently the norm of the response vector 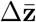 projected onto **β**, standardized by |**β**| (Hansen & Houle, 2008) (Fig. 1):

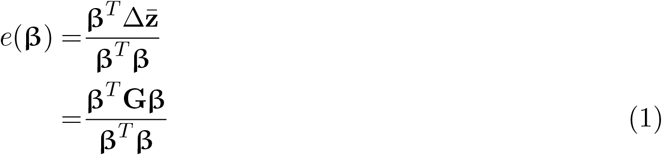

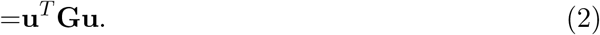

The numerator in (1) can be related to the rate of adaptation (change in mean fitness) under simple directional selection (Agrawal & Stinchcombe, 2009; Chevin, 2013).

Conditional evolvability *c*(**β**) is conceptually defined as the variance along **u** when evolution in other traits/directions is not allowed due to stabilizing selection (Hansen, 2003; Hansen *et al*., 2003a, 2019). With above notations, it is the conditional variance of the first element of **z**\* given the other elements (redefined here after Hansen & Houle, 2008):

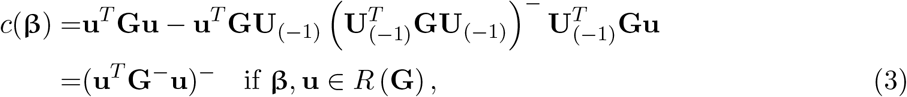

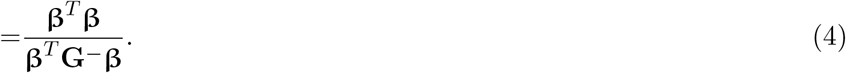

Here, the equation of (3) is from well-known results on the (generalized) inverse of a partitioned matrix (e.g., Schott, 2016, theorems 7.1 and 7.12), for which Hansen & Houle (2008) restated a proof in nonsingular **G**. A lack of conditional variance along **β** results in *c*(**β**) = 0, which happens when **G** is singular and **β** ∉ *R*(**G**); the expressions (3) and (4) do not hold in this case.

Autonomy *a*(**β**) is defined as the complement of the squared multiple correlation of the first element of **z**\* with respect to the other elements:

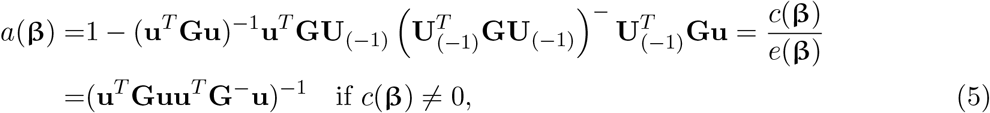

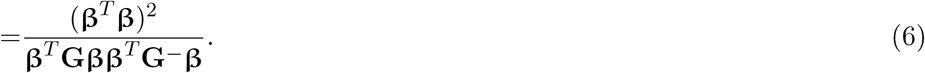

This quantity is undefined when *e*(**β**) = **u**^*T*^**Gu** = 0. Hansen & Houle (2008) also defined the integration *i*(**β**), which is the squared correlation just mentioned: *i*(**β**) = 1 − *a*(**β**).

Respondability *r*(**β**) is the magnitude of response standardized by that of selection, or the ratio of the norms of 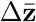 and **β**:

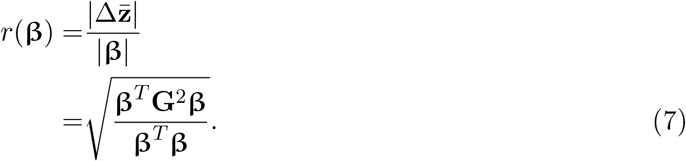

The definition and terminology of *r* here follows Hansen & Houle (2008), but it should be noted that Kirkpatrick (2009) independently devised an equivalent measure as the (relative) selection response (*R* in the latter author’s notation) with a slightly different standardization.

Flexibility *f*(**β**) quantifies similarity in direction between 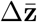 and **β** in terms of vector correlation (cosine of the angle formed by two vectors), and is also the ratio of *e*(**β**) to *r*(**β**) (Hansen & Houle, 2008; Marroig *et al*., 2009):

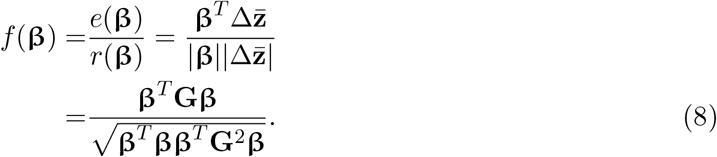

This quantity is undefined if 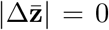, i.e., **β** ∈ *N*(**G**). Otherwise, 0 < *f*(**β**) ≤ 1 from the nonnegative definiteness of **G** and the definition of vector correlation. The term flexibility was coined by Marroig *et al*. (2009) due to Hansen’s suggestion, although the use of the vector correlation or angle was suggested in a few different works (Blows & Walsh, 2009; Rolian, 2009).

All the above indices are one-matrix measures that primarily concern characterization of a single **G** matrix. On the other hand, the following two indices concern pairwise comparisons between two **G** matrices in terms of (dis)similarity of the response vectors given the same **β**. The two **G** matrices and 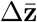 are denoted here with subscripts: **G**_*i*_, 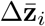. Response difference *d*(**β**) is a measure of dissimilarity, defined as a standardized difference between the two response vectors (Hansen & Houle, 2008):

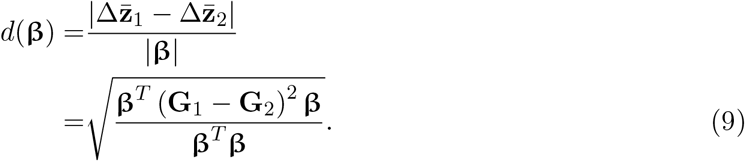

Response correlation *ρ*(**β**) is a measure of similarity in direction, defined as the vector correlation between the two response vectors (e.g., Cheverud, 1996; Cheverud & Marroig, 2007):

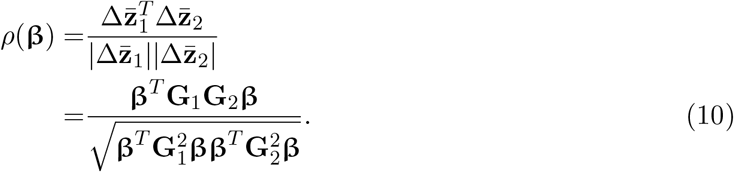

This quantity is undefined when at least one of 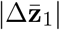 and 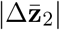 is 0. Otherwise, −1 ≤ *ρ*(**β**) ≤ 1.^3^

### 2.3 Ratio of quadratic forms

A key to derive expressions for average measures is that the measures defined above are simple or multiple ratios of quadratic forms in **β** or **u**. Ratios of quadratic forms, especially those in normal random variables, play a pivotal role in statistics, as many practically important test statistics can be written as a ratio of quadratic forms. Naturally, there is a vast body of literature regarding the distribution and moments of quadratic forms and ratios thereof (e.g., Magnus, 1986; Jones, 1986, 1987; Smith, 1989, 1993; Mathai & Provost, 1992; Hillier, 2001; Forchini, 2002; Hillier *et al*., 2009, 2014; Bao & Kan, 2013).

Regarding the ratio of quadratic forms in normal variables, there are several equivalent ways to derive its moments. One starts from the joint moment-generation function (e.g., Cressie *et al*., 1981; Meng, 2005) of quadratic forms, and typically results in integrations which does not have simple closed-form expressions (e.g., Magnus, 1986; Jones, 1986, 1987; Gupta & Kabe, 1998). Another way is to expand a ratio into infinite series, which in turn is integrated using the zonal and invariant polynomials (e.g., Smith, 1989, 1993; Hillier, 2001; Hillier *et al*., 2009, 2014). (See Bao & Kan (2013) for connections between these two approaches.) The latter way typically yields an infinite series including zonal or invariant polynomials, which can be evaluated with reasonable speed and accuracy with the aid of recent algorithmic developments (Hillier *et al*., 2009, 2014; Bao & Kan, 2013). This is the way followed in the present paper. The zonal polynomials are a special form of homogeneous polynomials in eigenvalues of a matrix which generalize powers of scalars into symmetric matrices (e.g., James, 1960, 1964; Muirhead, 1982, 2006; Gross & Richards, 1987; Mathai *et al*., 1995). The invariant polynomials are a generalization of the zonal polynomials into multiple matrix arguments (Davis, 1979, 1980; Chikuse, 1980, 2003; Chikuse & Davis, 1986b). A brief overview on these mathematical tools is provided in Appendix A.

A general result for the moments of a simple ratio of quadratic forms in normal variables is restated in the following proposition.

#### Proposition 1

(Smith (1989), Bao & Kan (2013)). *Let* **x** ∼ *N*_*p*_ (**μ, I**_*p*_), **A** *be a p* × *p symmetric matrix*, **B** *be a p* × *p nonnegative definite matrix, and m, n be positive real numbers. When the expectation of* 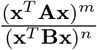*exists, it can be written as*

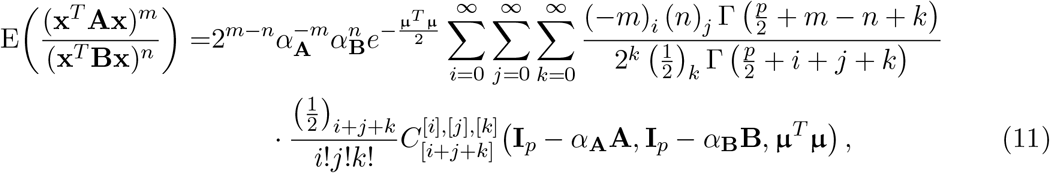

*where α*_**A**_ *and α*_**B**_ *are any positive constants that satisfy* 0 < *α*_**A**_ < 2/*λ*_max_(**A**) *and* 0 < *α*_**B**_ < 2/*λ*_max_(**B**), *with λ*_max_(·) *denoting the largest eigenvalue of the argument matrix*, (*a*)_*k*_ = *a*(*a* + 1) … (*a* + *k* − 1), *and* 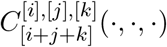 *are the*(*i, j, k*)*-th top-order invariant polynomials (whose explicit form is presented in Appendix A)*.

*Remark*. Conditions for the existence of the moment are stated in Bao & Kan (2013, proposition 1 therein): when **B** is positive definite, the moment exists if and only if 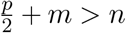; and when **B** is positive semidefinite, slightly different conditions apply, depending on the rank of **B** and structures of **A** and **B**. When *m* is an integer, the expressions can be simplified so that the summation for *i* disappears. Also, when **μ** = **0**_*p*_, they substantially simplify as all terms with *k* > 0 are zero. If **A** or **B** is asymmetric, it can be symmetrized by using **x**^*T*^**Ax** = **x**^*T*^**A**^*T*^**x** = **x**^*T*^ (**A** + **A**^*T*^)**x**/2.

In practice, the above series can be approximated by its partial sum, with recursive algorithms to calculate 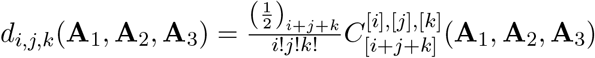 (Chikuse, 1987; Hillier *et al*., 2009, 2014; Bao & Kan, 2013). General covariance structures for **x** can be accommodated under certain conditions by transforming the variables and quadratic forms, as described in Appendix C. Appendix C also provides a similar result for multiple ratios of the form 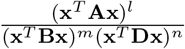.

### 2.4 Average measures for uniformly distributed selection

With the above theories, average measures can be readily derived as expectations with respect to random **β** under the assumption of normality. Previous treatments of evolvability measures (Hansen & Houle, 2008; Kirkpatrick, 2009) and random skewers analysis (Revell, 2007) typically assumed that **u** is uniformly distributed on the unit hypersphere in the *ps*dimensional space. This condition is entailed in spherical multivariate normal distributions of 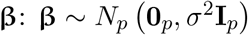 for arbitrary positive *σ*^2^. This section provides expressions of average measures under this condition. (Note that the normality of **β** is not necessary for **u** to be uniformly distributed; i.e., all results in this section hold as long as the distribution of **β** is spherically symmetric.) Expressions for a general normal case **β** ∼ *N*_*p*_ (**η, Σ**) are given in Appendix C. Most of the results below can be derived either by applying Proposition 1 or Proposition 4 in Appendix B, or directly as done in Appendix A.

The average evolvability *ē* is straightforward to obtain:s

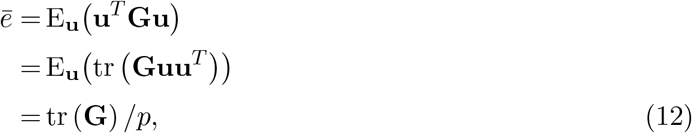

as derived by Hansen & Houle (2008). This is because E_**u**_(**uu**^*T*^) = **I**_*p*_/*p* under this condition.

Conditional evolvability *c*(**β**) and autonomy *a*(**β**) can be nonzero only when **β** ∈ *R*(**G**) (see above). When **G** is singular, this happens with probability 0 when **β** is continuously distributed across the entire space. Hence, the average conditional evolvability 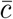 and the average autonomy *ā* are 0 when **G** is singular. Although the following expressions can operationally be applied to singular **G** by using **G**^−^ in place of **G**^−1^, such a result is invalid because the ratio expressions do not hold when **β** ∉ *R*(**G**).

When **G** is nonsingular, the average conditional evolvability 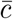 is:

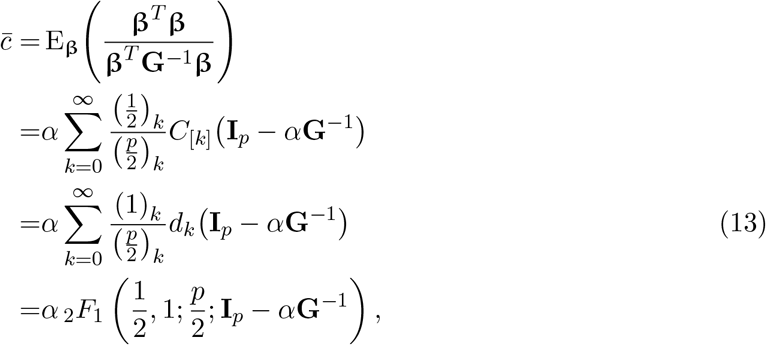

where *α* is any positive constant that satisfies 0 < *α* < 2/*λ*_max_ (**G**^−1^) (see Proposition 1), *C*_[*k*]_(·) are the *k*th top-order zonal polynomials, and _2_*F*_1_ (*a, b*; *c*; ·) is an alternative expression using the hypergeometric function of matrix argument with the three parameters *a, b*, and *c* (see Appendix A). When *p* = 2, this expression simplifies to the geometric mean of the two eigenvalues of **G** as mentioned by Hansen & Houle (2008) (Appendix D).

The average autonomy *ā* is, when **G** is nonsingular,

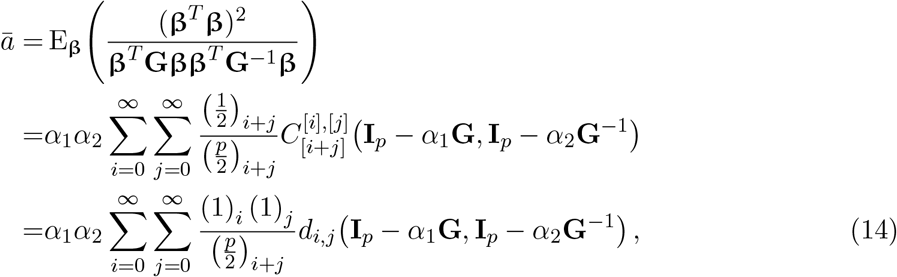

where *α*_1_ and *α*_2_ are to satisfy 0 < *α*_1_ < 2/*λ*_max_(**G**) and 0 < *α*_2_ < 2/*λ*_max_ (**G**^−1^). The average integration *ī* is simply 1 − *ā* (and equals 1 when **G** is singular).

The average respondability 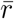 is

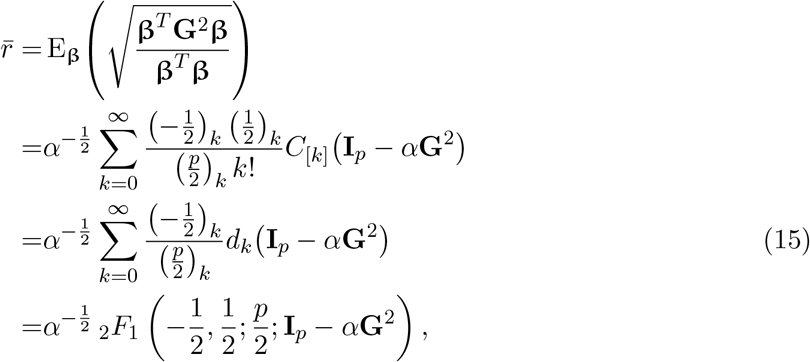

where *α* is to satisfy 0 < *α* < 2/*λ*_max_ (**G**^2^) (note that *λ*_max_ (**G**^2^) = *λ*_max_(**G**)^2^ since **G** is nonnegative definite). Kirkpatrick (2009) discussed on an equivalent quantity, the average (relative) selection response (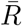 there), but did not provide a closed-form expression for arbitrary **G**. It is possible to show his expression for the special case with complete integration is entailed in the present result (Appendix D).

The average flexibility 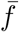 is

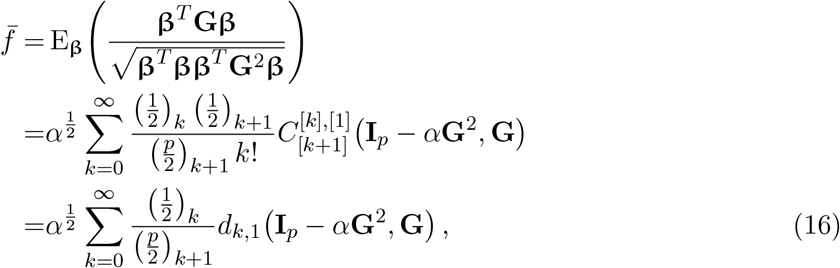

where *α* is to satisfy 0 < *α* < 2/*λ*_max_ (**G**^2^). In general, 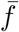 is well defined even if **G** is singular, unless **β** ∈ *N*(**G**) with nonzero probability (which of course does not happen here).

The average response difference 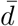 has a form essentially identical to 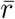 except for the argument matrix

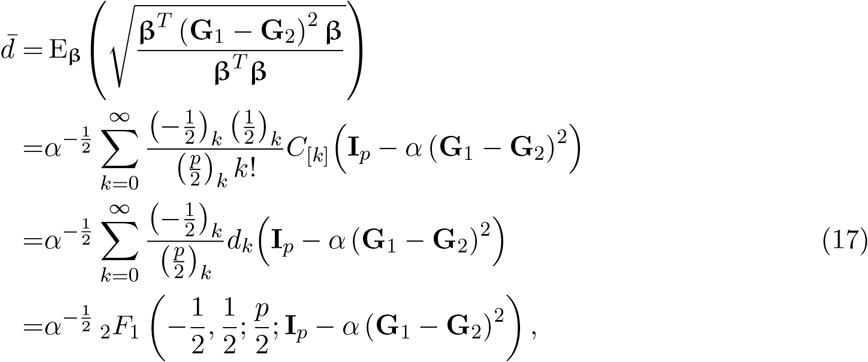

where *α* is to satisfy 0 < *α* < 2/*λ*_max_ ((**G**_1_ − **G**_2_)^2^).

The average response correlation 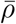 is

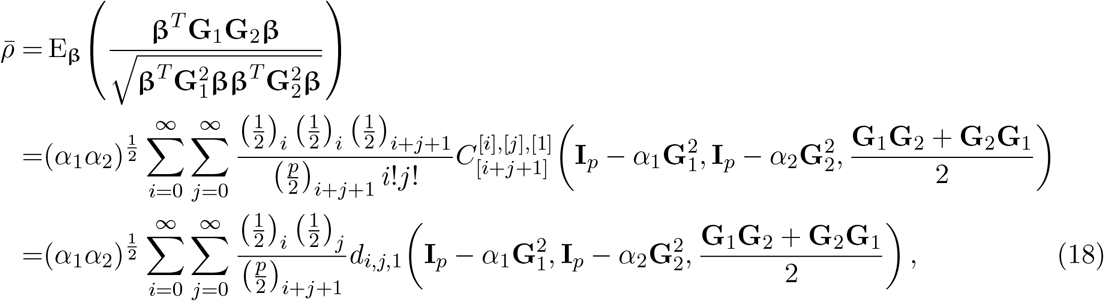

where *α*_1_ and *α*_2_ are to satisfy 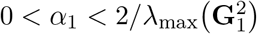 and 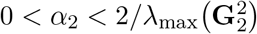.

### 2.5 Truncation error

It is seen that the successive terms in the above series decrease in magnitude, thus a partial sum up to certain higher-order terms can in principle be used as an approximation of the exact value of the infinite series. Although each term can be easily evaluated with a recursive algorithm (Bao & Kan, 2013; Hillier *et al*., 2014), accurate numerical evaluation involving higher-order terms is practically a computer-intensive problem. The speed of numerical convergence and computational time will be examined in the next section.

The choice of *α* is arbitrary within the constraint 0 < *α* < 2/*λ*_max_ with respect to the relevant matrix (see above), but influences the signs of the terms as well as the speed of convergence. When 0 < *α* ≤ 1/*λ*_max_, all the zonal and invariant polynomials in the above expressions are positive because the argument matrices have only nonnegative eigenvalues. Therefore, all terms in the summations in 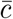 (13), *ā* (14), 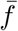 (16), and 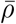 (18) are positive, so truncation results in (slight) underestimation. On the other hand, all terms in 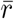 (15) and 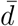 (17), except those for *k* = 0, are negative in the same condition, so truncation results in overestimation. If 1/*λ*_max_ < *α* < 2/*λ*_max_, the signs of the polynomials possibly, though not always, fluctuate, rendering the behavior of the series less predictable.

It is possible to derive bounds for the truncation errors in 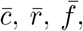, and 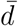 by applying the method of Hillier *et al*. (2009, theorem 6) with slight modifications, if **G** is nonsingular and the constants *α* are taken within 0 < *α* ≤ 1/*λ*_max_. Let 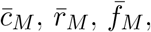, and 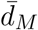 be the relevant partial sums up to *k* = *M*. The truncation error bounds are:

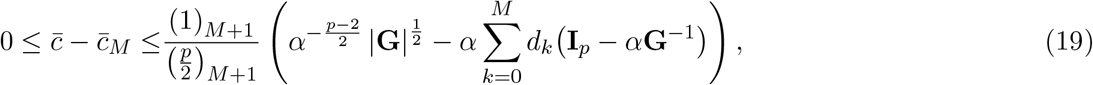

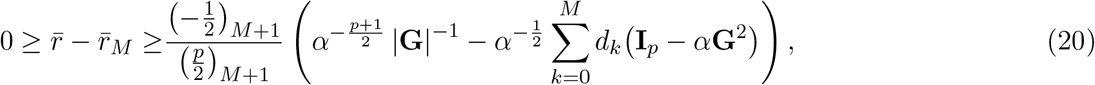

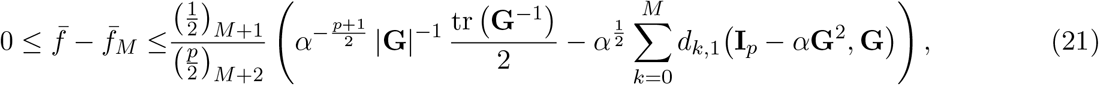

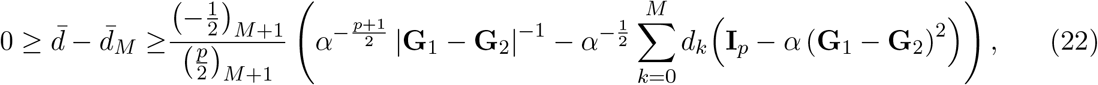

where the constants *α* are as in the corresponding definitions (but to satisfy 0 < *α* ≤ 1/*λ*_max_). Unfortunately, the same method are not applicable to *ā* or 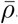. Also, if **G** (or **G**_1_ − **G**_2_ in case of 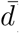) is singular, these error bounds do not hold.

Empirically, a larger value of *α* often improves the speed of convergence, as more eigenvalues of the argument matrix approach 0, but this should be used with caution since the above error bounds are not strictly applicable if *α* > 1/*λ*_max_. To be precise, the error bounds should hold even under this condition provided that all the zonal/invariant polynomials above *k* = *M* are nonnegative, but it is not obvious in which case that condition can be shown to be true.

## 3 Numerical evaluation

### 3.1 Software implementation

There exist two `Matlab` programs distributed by Raymond M. Kan (https://www-2.rotman.utoronto.ca/~kan/) for evaluating the moments of simple ratios of quadratic forms in normal variables (Hillier *et al*., 2009, 2014; Bao & Kan, 2013), which can be used to evaluate 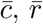, and 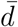 if a suitable environment is available. Alternatively, these average measures could be evaluated as hypergeometric functions of matrix argument, for which an efficient, general algorithm has been developed by Koev & Edelman (2006) and implemented in the `R` package `HypergeoMat` (Laurent, 2022). However, that seems relatively inefficient in the present applications where the terms corresponding to all non-top-order partitions can be omitted. None of these can be used to evaluate moments of multiple ratios like *ā*, 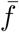, and 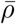.

In order to make all above expressions available in the popular statistical software environment `R`, the present author has developed an independent implementation of the algorithms of Hillier *et al*. (2014) and Bao & Kan (2013) and published it as the package `qfratio` on the CRAN repository (Watanabe, 2023). This package can be used to evaluate general moments of ratios of quadratic forms in normal variables with arbitrary mean and covariance (under certain constraints for singular cases as discussed in Appendix C.1). It uses `RcppEigen` (Bates & Eddelbuettel, 2013) to effectively execute the computationally intensive recursive algorithms.

### 3.2 Numerical experiment: methods

Using the new `R` package, numerical experiments were conducted in a small set of artificial covariance matrices to evaluate behaviors and efficiency of the present series expressions. For each of *p* = 5, 20, 50, and 200, three different eigenvalue conformations were used; two were set to have single large eigenvalues with different magnitudes of integration, *V*_rel_ = 0.1, 0.5— corresponding to relatively weak and rather strong integration, respectively—and one was set to have a quadratically decreasing series of eigenvalues (for the algorithms see Watanabe (2022) and the `R` package `eigvaldisp` (https://github.com/watanabe-j/eigvaldisp)). The matrices with these eigenvalue conformations are referred to as **G**_0.1_, **G**_0.5_, and **G**_Q_. For each of these, two eigenvector conformations were used so that the resultant covariance matrices were either diagonal matrices of the eigenvalues or correlation matrices with the same eigenvalues constructed with Givens rotations (Davies & Higham, 2000). For the one-matrix measures (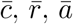, and 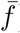), the choice of eigenvectors is inconsequential. For the two-matrix measures (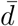 and 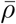), comparisons were made between pairs of matrices with identical and different eigenvectors. The results were also compared to those from the traditional Monte Carlo evaluation using the `qfratio` package and the delta method approximation of Hansen & Houle (2008, 2009) using new codes.

When possible, an average computational time from 10 runs is recorded for the present series evaluation and Monte Carlo evaluation with 10,000 iterations in each condition. This is except for 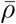 in certain conditions with *p* ≥ 50 where only a single run was timed for the series evaluation as it did not reach numerical convergence (below). Where a truncation error bound is available, an order of series evaluation *M* was chosen so that the error is below 1.0 × 10^−8^. Where an error bound is unavailable, it was aimed to evaluate the series until the result rounded to the digit of 10^−9^ appeared stable. (These values were arbitrarily chosen for benchmarking purposes, and not to be used as a guide for applications.) No formal timing was done for the delta method approximation as it is not computer-intensive. All computational times reported are in elapsed time rather than CPU time. The calculations were executed on a regular desktop PC with Intel^®^ Core i7-8700 CPU and 16 GB RAM, running `R` version 4.2.3 (R Core Team, 2023) with its built-in BLAS. For those problems that required larger computational memory for convergence (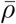 with **G**_0.5_ and *p* ≥ 20; see below), another machine with 256 GB RAM was used to calculate accurate values. The codes used in the numerical experiments are provided as a Supplementary Material, as well as maintained on a GitHub repository (https://github.com/watanabe-j/avrevol).

### 3.3 Numerical experiment: results

The present series evaluations as implemented in the `R` package `qfratio` reached numerical convergence within reasonable computational times in all cases except 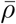 with large (*p* = 200) or highly integrated (**G**_0.5_) matrices (detailed below). The order of evaluation *M* required to accomplish a given level of accuracy varied across evolvability measures and eigenvalue conformations, but in most cases increased with *p*(Figs. 2–5; Tables 1–6). The rate of scaling with respect to *p* tended to be more rapid in the series evaluation than in Monte Carlo evaluation, when the required accuracy for the former and the number of iteration for the latter are pre-determined as described above.

**Table 1.** Results of numerical experiments for 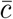. Tabulated are combinations of eigenvalue conformation and number of variables *p*, order of evaluation *M* required to attain the accuracy specified in text, partial sum from series evaluation 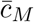 and its error bound, average computational time of series and Monte Carlo evaluations, *t*_S_ and *t*_MC_, empirical standard error from one Monte Carlo run with 10,000 iterations SE_MC_, Hansen & Houle’s delta method approximation *ĉ*_HH_, and its deviation from 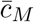 in percent Δ_HH_.

| Matrix | $p$ | $M$ | $\bar{c}_M$ | Error bound | $t_S$ (ms) | $t_{MC}$ (ms) | $SE_{MC}$ | $\hat{c}_{HH}$ | $\Delta_{HH}$ |
| --- | --- | --- | --- | --- | --- | --- | --- | --- | --- |
| $\mathbf{G}_{0.1}$ | 5 | 33 | 0.8289 | $6.637 \times 10^{-9}$ | 0.5 | 2.8 | $2.029 \times 10^{-3}$ | 0.8187 | −1.2% |
| | 20 | 19 | 0.7194 | $9.622 \times 10^{-9}$ | 0.6 | 10.9 | $5.355 \times 10^{-4}$ | 0.7188 | −0.1% |
| | 50 | 9 | 0.6977 | $9.309 \times 10^{-9}$ | 0.9 | 32.8 | $2.019 \times 10^{-4}$ | 0.6976 | 0.0% |
| | 200 | 5 | 0.6872 | $2.978 \times 10^{-9}$ | 14.8 | 239.2 | $4.753 \times 10^{-5}$ | 0.6872 | 0.0% |
| $\mathbf{G}_{0.5}$ | 5 | 130 | 0.4016 | $4.687 \times 10^{-9}$ | 0.5 | 2.6 | $2.138 \times 10^{-3}$ | 0.3803 | −5.3% |
| | 20 | 25 | 0.3097 | $8.858 \times 10^{-9}$ | 0.6 | 10.7 | $2.580 \times 10^{-4}$ | 0.3094 | −0.1% |
| | 50 | 10 | 0.2991 | $5.217 \times 10^{-9}$ | 1.0 | 32.8 | $9.143 \times 10^{-5}$ | 0.2991 | 0.0% |
| | 200 | 5 | 0.2944 | $3.512 \times 10^{-9}$ | 15.1 | 242.3 | $2.077 \times 10^{-5}$ | 0.2944 | 0.0% |
| $\mathbf{G}_Q$ | 5 | 290 | 0.4730 | $7.239 \times 10^{-9}$ | 0.6 | 2.6 | $3.212 \times 10^{-3}$ | 0.4456 | −5.8% |
| | 20 | 1800 | 0.1501 | $6.932 \times 10^{-9}$ | 1.1 | 10.9 | $1.132 \times 10^{-3}$ | 0.1468 | −2.2% |
| | 50 | 7800 | 0.0632 | $8.195 \times 10^{-9}$ | 3.4 | 32.6 | $4.837 \times 10^{-4}$ | 0.0627 | −0.7% |
| | 200 | 260 000 | 0.0162 | $4.445 \times 10^{-10}$ | 120.4 | 243.4 | $1.233 \times 10^{-4}$ | 0.0162 | 0.1% |

**Figure 2.**
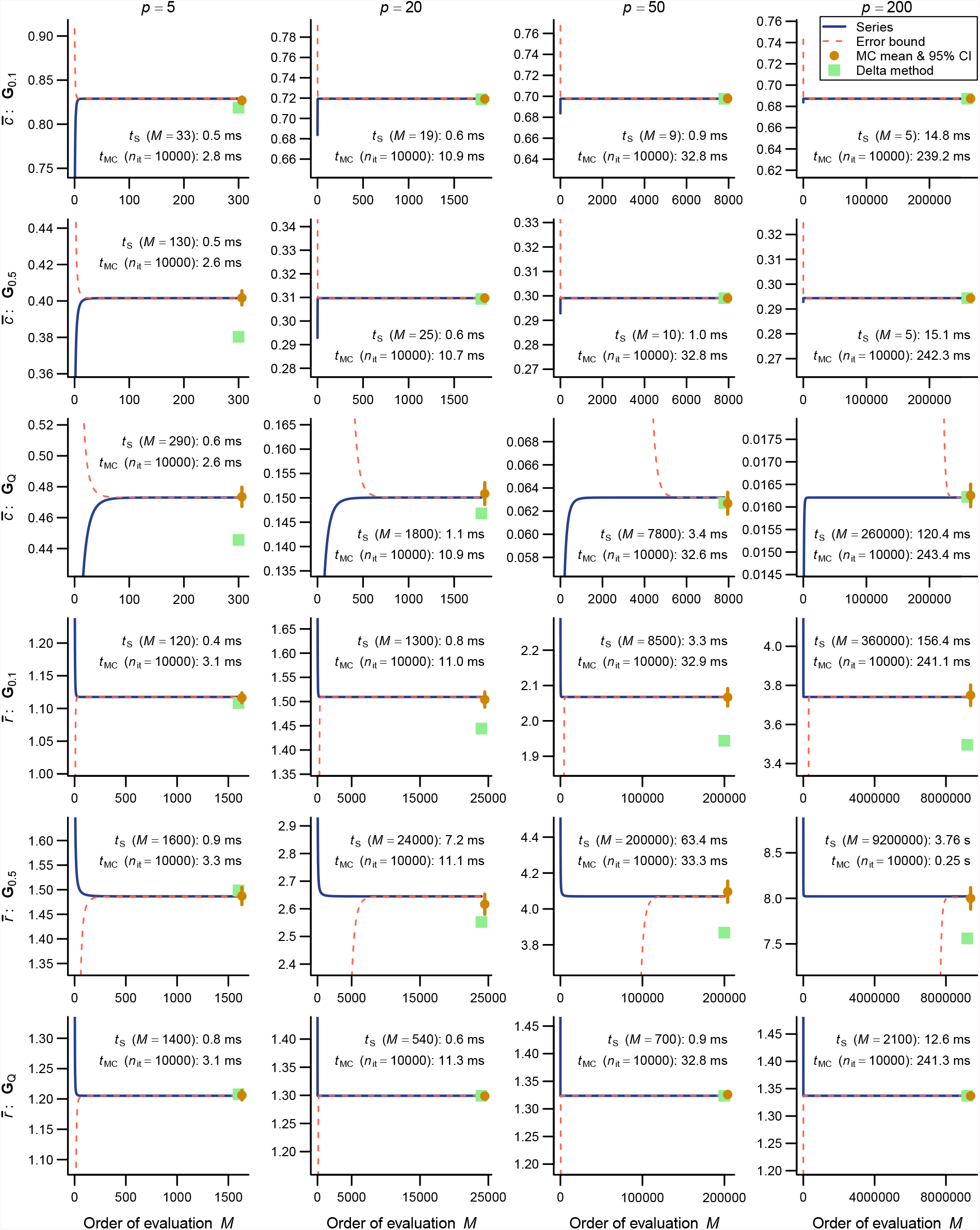
Results of numerical experiments for 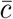 (top three rows) and 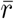 (bottom three rows). Partial sums from series expressions are shown in blue solid lines, and associated truncation error bounds as deviations from the partial sum are in red broken lines, across varying orders of evaluation *M*. Mean and its 95% confidence interval from a Monte Carlo run with 10,000 iterations are shown in brown circles and vertical bars. Delta method approximations by Hansen & Houle (2008, 2009) are shown in green squares. Rows correspond to eigenvalue conformations (see text) and columns to number of variables *p*. Each panel represents *±*10% region of the terminal value of series evaluation. Average computational times from 10 runs for the series *t*_S_ and Monte Carlo evaluations with 10,000 iteration *t*_MC_ are shown in each panel, along with *M* required for numerical convergence in the former.

For 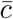, the computational time required for the error bound to numerically converge below 1.0 × 10^−8^ was at most in the order of ∼0.1 s and compared favorably with the Monte Carlo evaluation with 10,000 iterations across the entire range of *p* examined (Fig. 2; Table 1). For 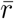 and 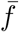, the computational time varied from ∼1.5 ms in *p* = 5 to several seconds in *p* = 200, and in some eigenvalue conformations the series evaluation was outcompeted by the Monte Carlo evaluation with a fixed number of iterations in terms of plain computational time (Tables 2, 3). However, it is to be remembered that results from Monte Carlo evaluations always involve stochastic error, whose standard error (SE) scales only with the inverse square root of the number of iteration (i.e., 100 times as many calculations are required to reduce the SE by the factor of 1/10). It is also notable that the partial sums themselves typically appeared to reach asymptotes much earlier than the error bounds did in these measures (Figs. 2, 3). It took substantially longer time to accurately evaluate *ā* than the other one-matrix measures, since it involves a double series so that the computational burden scales roughly with *M*^2^. Consequently, the computational time required for *ā* to numerically converge at the order of 10^−9^ varied from ∼7 ms to ∼170 s (Fig. 3; Table 4). Technically, this large computational time was partly because of the necessity to scale individual terms in the evaluation of *ā* to avoid numerical overflow and underflow.

**Table 2.** Results of numerical experiments for 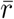. See Table 1 for legend.

| Matrix | $p$ | $M$ | $\bar{r}_M$ | Error bound | $t_S$ (ms) | $t_{MC}$ (ms) | $SE_{MC}$ | $\hat{r}_{HH}$ | $\Delta_{HH}$ |
| --- | --- | --- | --- | --- | --- | --- | --- | --- | --- |
| $\mathbf{G}_{0.1}$ | 5 | 120 | 1.1176 | $-6.518 \times 10^{-9}$ | 0.4 | 3.1 | $3.897 \times 10^{-3}$ | 1.1082 | -0.8% |
| | 20 | 1300 | 1.5095 | $-3.571 \times 10^{-9}$ | 0.8 | 11.0 | $7.891 \times 10^{-3}$ | 1.4443 | -4.3% |
| | 50 | 8500 | 2.0677 | $-9.517 \times 10^{-9}$ | 3.3 | 32.9 | $1.279 \times 10^{-2}$ | 1.9439 | -6.0% |
| | 200 | 360 000 | 3.7407 | $-9.967 \times 10^{-10}$ | 156.4 | 241.1 | $2.630 \times 10^{-2}$ | 3.4955 | -6.6% |
| $\mathbf{G}_{0.5}$ | 5 | 1600 | 1.4865 | $-8.769 \times 10^{-9}$ | 0.9 | 3.3 | $8.931 \times 10^{-3}$ | 1.4986 | 0.8% |
| | 20 | 24 000 | 2.6450 | $-8.893 \times 10^{-9}$ | 7.2 | 11.1 | $1.872 \times 10^{-2}$ | 2.5521 | -3.5% |
| | 50 | 200 000 | 4.0700 | $-6.271 \times 10^{-9}$ | 63.4 | 33.3 | $2.966 \times 10^{-2}$ | 3.8681 | -5.0% |
| | 200 | 9 200 000 | 8.0216 | $-9.269 \times 10^{-9}$ | 3759.2 | 245.1 | $6.091 \times 10^{-2}$ | 7.5602 | -5.8% |
| $\mathbf{G}_Q$ | 5 | 1400 | 1.2052 | $-7.504 \times 10^{-9}$ | 0.8 | 3.1 | $4.079 \times 10^{-3}$ | 1.2078 | 0.2% |
| | 20 | 540 | 1.2992 | $-9.765 \times 10^{-9}$ | 0.6 | 11.3 | $2.585 \times 10^{-3}$ | 1.2990 | 0.0% |
| | 50 | 700 | 1.3237 | $-7.742 \times 10^{-9}$ | 0.9 | 32.8 | $1.737 \times 10^{-3}$ | 1.3237 | 0.0% |
| | 200 | 2100 | 1.3370 | $-3.993 \times 10^{-9}$ | 12.6 | 241.3 | $8.872 \times 10^{-4}$ | 1.3370 | 0.0% |

**Table 3.** Results of numerical experiments for 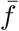. See Table 1 for legend.

| Matrix | $p$ | $M$ | $\bar{f}_M$ | Error bound | $t_S$ (ms) | $t_{MC}$ (ms) | $SE_{MC}$ |
| --- | --- | --- | --- | --- | --- | --- | --- |
| $\mathbf{G}_{0.1}$ | 5 | 160 | 0.9031 | $3.628 \times 10^{-9}$ | 0.7 | 3.9 | $5.258 \times 10^{-4}$ |
| | 20 | 1600 | 0.7172 | $5.471 \times 10^{-9}$ | 1.3 | 12.6 | $1.459 \times 10^{-3}$ |
| | 50 | 11 000 | 0.5718 | $8.857 \times 10^{-10}$ | 6.0 | 38.4 | $1.986 \times 10^{-3}$ |
| | 200 | 380 000 | 0.3743 | $1.267 \times 10^{-9}$ | 258.9 | 298.6 | $2.214 \times 10^{-3}$ |
| $\mathbf{G}_{0.5}$ | 5 | 2300 | 0.6569 | $6.730 \times 10^{-9}$ | 1.5 | 4.0 | $1.382 \times 10^{-3}$ |
| | 20 | 33 000 | 0.3987 | $8.228 \times 10^{-9}$ | 11.6 | 12.8 | $1.626 \times 10^{-3}$ |
| | 50 | 240 000 | 0.2772 | $9.889 \times 10^{-9}$ | 92.6 | 37.9 | $1.627 \times 10^{-3}$ |
| | 200 | 9 900 000 | 0.1559 | $3.968 \times 10^{-9}$ | 6052.1 | 295.2 | $1.434 \times 10^{-3}$ |
| $\mathbf{G}_Q$ | 5 | 2800 | 0.8082 | $9.271 \times 10^{-9}$ | 1.6 | 3.9 | $1.119 \times 10^{-3}$ |
| | 20 | 910 | 0.7614 | $9.615 \times 10^{-9}$ | 1.1 | 12.9 | $7.039 \times 10^{-4}$ |
| | 50 | 930 | 0.7518 | $8.400 \times 10^{-9}$ | 1.9 | 38.2 | $4.640 \times 10^{-4}$ |
| | 200 | 2400 | 0.7470 | $1.679 \times 10^{-10}$ | 31.7 | 298.6 | $2.364 \times 10^{-4}$ |

**Table 4.**
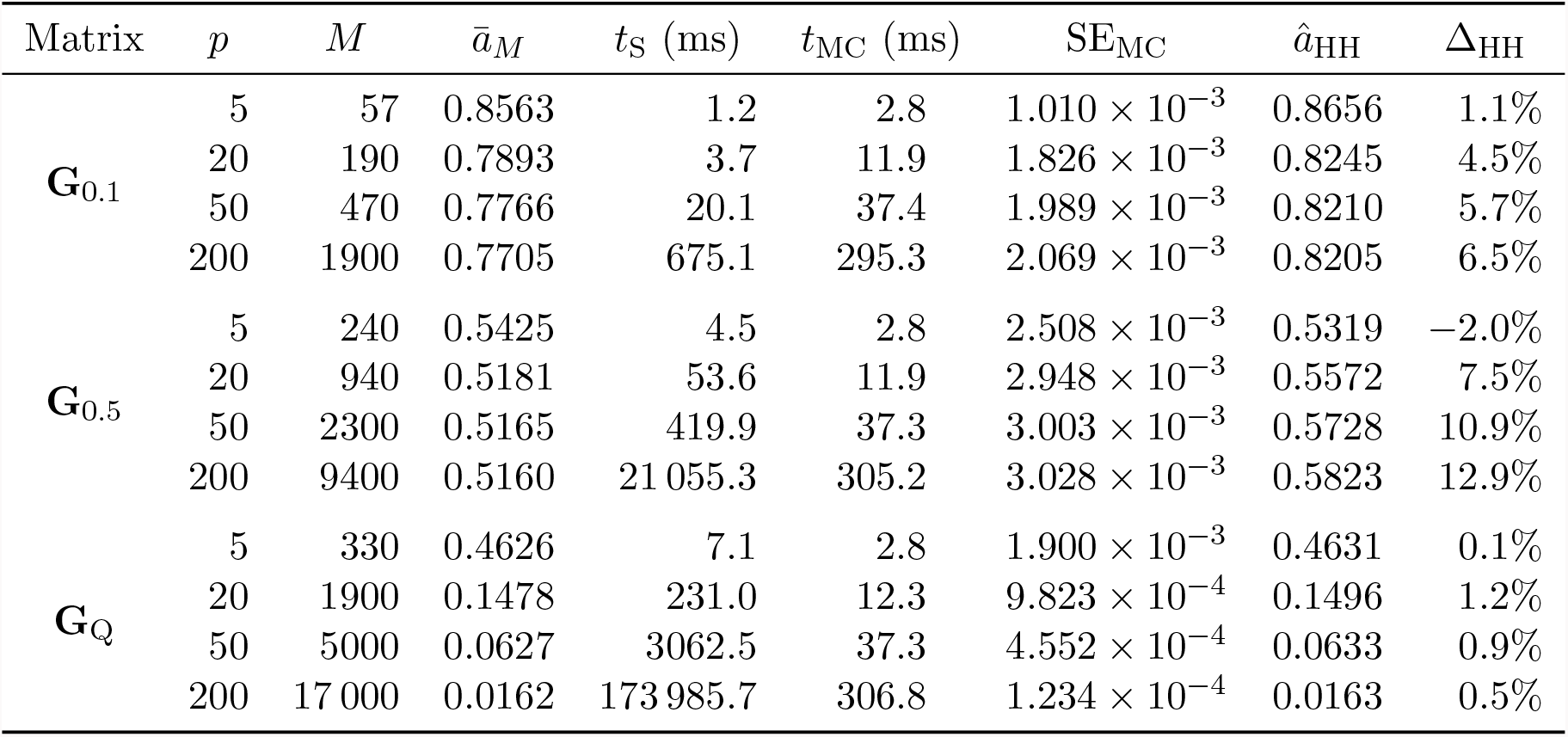
Results of numerical experiments for *ā*. See Table 1 for legend.

**Figure 3.**
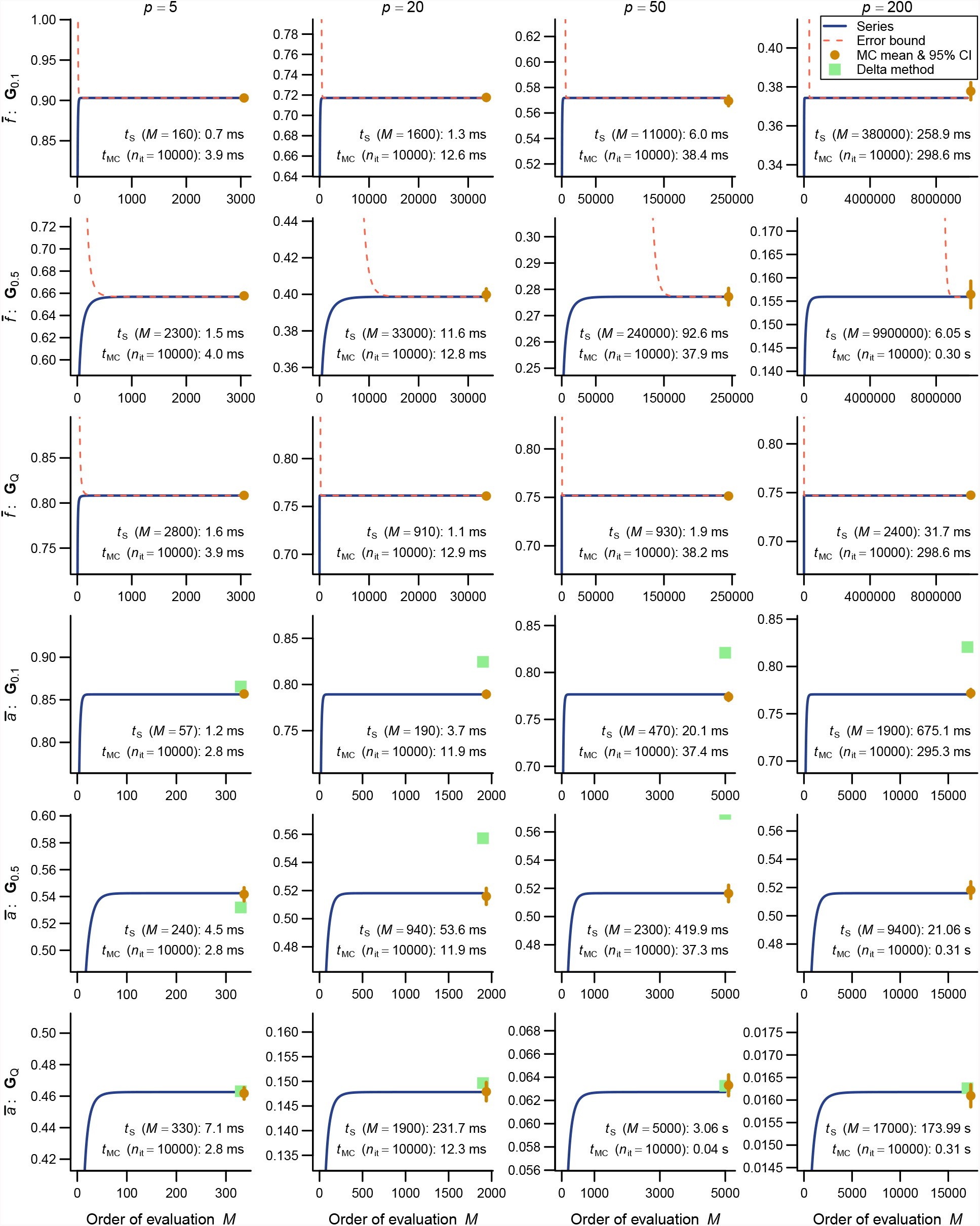
Results of numerical experiments for 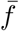 (top three rows) and *ā* (bottom three rows). See Figure 2 for legend. Delta method approximations are not available for 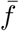, and truncation error bounds are not available for *ā*. In some results for *ā*, delta method approximation is too inaccurate to be plotted in the panels.

For two-matrix comparisons, 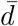 behaved similarly to 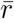 with which it shares the same functional form (Fig. 4; Table 5). In some comparisons between matrices with the same eigenvalues and different eigenvectors, the error bound of 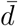 could not be calculated because the argument matrix ((**G**_1_ − **G**_2_)^2^) was singular due to the artificial conformations of eigenvalues in **G**_0.1_ and **G**_0.5_ where the trailing eigenvalues are identical in magnitude. Series evaluation of 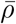 took substantially more computational time and memory than the other measures. For *p* ≥ 20, the series could not be evaluated until numerical convergence in any comparisons that involve **G**_0.5_ due to excessively large memory requirement for the 16 GB RAM machine used here for main calculations, at least when the accuracy in the order of 10^−9^ is aimed at (Fig. 5; Table 6 involves accurate values from a larger machine). This is apparently because 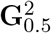 has rather small eigenvalues, which translate into extremely large magnitudes of higher-order terms, *d*_*i,j*,1_ in (18). For *p* = 200, accurate series evaluation of 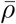 seemed infeasible in most combinations for too much computational memory required. The only exception in the present test cases is the comparison between two **G**_Q_’s with different eigenvectors where the series converged fairly rapidly (Table 6).

**Table 5.** Results of numerical experiments for 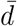. Comparisons were made in pairs of covariance matrices with same and different eigenvectors (evecs). Error bound was unavailable for comparisons between two **G**_0.1_’s and two **G**_0.5_’s with different eigenvectors (see text). See Table 1 for further explanations.

| Matrices | $p$ | $M$ | $\bar{d}_M$ | Error bound | $t_S$ (ms) | $t_{MC}$ (ms) | $SE_{MC}$ | $\hat{d}_{HH}$ | $\Delta_{HH}$ |
| --- | --- | --- | --- | --- | --- | --- | --- | --- | --- |
| $\mathbf{G}_{0.1}$ vs. $\mathbf{G}_{0.5}$,<br>same evecs | 5 | 170 | 0.7256 | $-5.603 \times 10^{-9}$ | 0.6 | 3.2 | $2.914 \times 10^{-3}$ | 0.7189 | -0.9% |
| | 20 | 3900 | 1.4375 | $-8.787 \times 10^{-9}$ | 1.8 | 11.7 | $9.087 \times 10^{-3}$ | 1.3736 | -4.4% |
| | 50 | 34 000 | 2.2428 | $-6.090 \times 10^{-9}$ | 11.2 | 33.0 | $1.572 \times 10^{-2}$ | 2.1176 | -5.6% |
| | 200 | 1 600 000 | 4.4371 | $-1.852 \times 10^{-9}$ | 651.5 | 241.0 | $3.287 \times 10^{-2}$ | 4.1696 | -6.0% |
| $\mathbf{G}_{0.1}$ vs. $\mathbf{G}_Q$,<br>same evecs | 5 | 2900 | 0.4409 | $-9.244 \times 10^{-9}$ | 1.4 | 3.2 | $1.372 \times 10^{-3}$ | 0.4429 | 0.5% |
| | 20 | 1200 | 1.1657 | $-5.016 \times 10^{-9}$ | 1.0 | 11.4 | $4.027 \times 10^{-3}$ | 1.1456 | -1.7% |
| | 50 | 11 000 | 1.8752 | $-4.953 \times 10^{-9}$ | 4.3 | 33.0 | $9.865 \times 10^{-3}$ | 1.7781 | -5.2% |
| | 200 | 580 000 | 3.6679 | $-2.875 \times 10^{-9}$ | 243.1 | 240.9 | $2.446 \times 10^{-2}$ | 3.4190 | -6.8% |
| $\mathbf{G}_{0.5}$ vs. $\mathbf{G}_Q$,<br>same evecs | 5 | 1200 | 0.8575 | $-6.347 \times 10^{-9}$ | 0.9 | 3.2 | $2.907 \times 10^{-3}$ | 0.8620 | 0.5% |
| | 20 | 9000 | 2.4270 | $-9.410 \times 10^{-9}$ | 3.1 | 11.2 | $1.346 \times 10^{-2}$ | 2.3229 | -4.3% |
| | 50 | 80 000 | 3.9804 | $-9.196 \times 10^{-9}$ | 25.7 | 33.5 | $2.607 \times 10^{-2}$ | 3.7449 | -5.9% |
| | 200 | 3 900 000 | 8.0124 | $-2.997 \times 10^{-9}$ | 1589.6 | 240.2 | $5.761 \times 10^{-2}$ | 7.5050 | -6.3% |
| Two $\mathbf{G}_{0.1}$,<br>different evecs | 5 | 7000 | 0.8330 | — | 2.6 | 3.4 | $3.243 \times 10^{-3}$ | 0.8465 | 1.6% |
| | 20 | 9900 | 1.7493 | — | 3.4 | 11.2 | $8.644 \times 10^{-3}$ | 1.7500 | 0.0% |
| | 50 | 93 000 | 2.7882 | — | 28.5 | 33.2 | $1.413 \times 10^{-2}$ | 2.7693 | -0.7% |
| | 200 | 4 900 000 | 5.5980 | — | 1943.8 | 242.0 | $2.892 \times 10^{-2}$ | 5.5357 | -1.1% |
| Two $\mathbf{G}_{0.5}$,<br>different evecs | 5 | 7000 | 1.8627 | — | 2.6 | 3.4 | $7.318 \times 10^{-3}$ | 1.8929 | 1.6% |
| | 20 | 9900 | 3.9116 | — | 3.4 | 11.4 | $1.937 \times 10^{-2}$ | 3.9131 | 0.0% |
| | 50 | 93 000 | 6.2347 | — | 28.4 | 33.5 | $3.188 \times 10^{-2}$ | 6.1923 | -0.7% |
| | 200 | 4 900 000 | 12.5174 | — | 1938.2 | 240.5 | $6.569 \times 10^{-2}$ | 12.3783 | -1.1% |
| Two $\mathbf{G}_Q$,<br>different evecs | 5 | 3600 | 1.0412 | $-9.597 \times 10^{-9}$ | 1.6 | 3.4 | $3.909 \times 10^{-3}$ | 1.0546 | 1.3% |
| | 20 | 1200 | 1.2087 | $-4.817 \times 10^{-9}$ | 1.0 | 11.5 | $2.174 \times 10^{-3}$ | 1.2095 | 0.1% |
| | 50 | 2300 | 1.2433 | $-3.719 \times 10^{-9}$ | 1.9 | 33.1 | $1.344 \times 10^{-3}$ | 1.2434 | 0.0% |
| | 200 | 14 000 | 1.2595 | $-8.004 \times 10^{-11}$ | 20.4 | 243.9 | $6.708 \times 10^{-4}$ | 1.2595 | 0.0% |
| $\mathbf{G}_{0.1}$ vs. $\mathbf{G}_{0.5}$,<br>different evecs | 5 | 450 | 1.4432 | $-8.000 \times 10^{-9}$ | 0.7 | 3.2 | $5.645 \times 10^{-3}$ | 1.4478 | 0.3% |
| | 20 | 9900 | 2.9662 | $-9.012 \times 10^{-9}$ | 3.4 | 11.2 | $1.587 \times 10^{-2}$ | 2.8938 | -2.4% |
| | 50 | 93 000 | 4.6991 | $-9.283 \times 10^{-9}$ | 29.8 | 33.2 | $2.698 \times 10^{-2}$ | 4.5298 | -3.6% |
| | 200 | 4 900 000 | 9.4026 | $-5.707 \times 10^{-9}$ | 2004.6 | 242.8 | $5.638 \times 10^{-2}$ | 8.9991 | -4.3% |
| $\mathbf{G}_{0.1}$ vs. $\mathbf{G}_Q$,<br>different evecs | 5 | 690 | 0.9545 | $-9.978 \times 10^{-9}$ | 0.7 | 3.2 | $3.271 \times 10^{-3}$ | 0.9630 | 0.9% |
| | 20 | 2200 | 1.4997 | $-9.404 \times 10^{-9}$ | 1.3 | 11.3 | $6.282 \times 10^{-3}$ | 1.4564 | -2.9% |
| | 50 | 13 000 | 2.0938 | $-5.509 \times 10^{-9}$ | 5.2 | 33.0 | $1.160 \times 10^{-2}$ | 1.9798 | -5.4% |
| | 200 | 590 000 | 3.7798 | $-9.522 \times 10^{-9}$ | 248.9 | 241.3 | $2.529 \times 10^{-2}$ | 3.5236 | -6.8% |
| $\mathbf{G}_{0.5}$ vs. $\mathbf{G}_Q$,<br>different evecs | 5 | 6800 | 1.5135 | $-9.579 \times 10^{-9}$ | 2.5 | 3.2 | $5.735 \times 10^{-3}$ | 1.5199 | 0.4% |
| | 20 | 8600 | 2.7842 | $-9.060 \times 10^{-9}$ | 3.2 | 11.6 | $1.592 \times 10^{-2}$ | 2.6625 | -4.4% |
| | 50 | 82 000 | 4.2077 | $-9.025 \times 10^{-9}$ | 26.4 | 33.4 | $2.843 \times 10^{-2}$ | 3.9590 | -5.9% |
| | 200 | 3 900 000 | 8.1259 | $-9.530 \times 10^{-9}$ | 1577.7 | 237.9 | $5.910 \times 10^{-2}$ | 7.6115 | -6.3% |

**Table 6.** Results of numerical experiments for 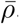. Parentheses around 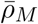 denote values that did not reach numerical convergence with specified *M* ; for these conditions, more accurate values with larger *M* were calculated on a separate machine and are shown for reference 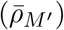—this calculation was not timed. Asterisks after *t*_S_ denote times from single runs rather than 10 runs. For *p* = 200, it was infeasible to apply the series evaluation in most combinations for excessive computational memory required. See Table 5 for further explanations.

| Matrices | $p$ | $M$ | $\bar{\rho}_M$ | $\bar{\rho}_{M'}$ | $t_S$ (s) | $t_{MC}$ (s) | $SE_{MC}$ |
| --- | --- | --- | --- | --- | --- | --- | --- |
| $\mathbf{G}_{0.1}$ vs. $\mathbf{G}_{0.5}$,<br>same evecs | 5 | 2600 | 0.9034 | — | 0.60 | 0.00 | $6.306 \times 10^{-4}$ |
| | 20 | 4000 | (0.8960) | 0.8990 | 2.27 | 0.01 | $7.670 \times 10^{-4}$ |
| | 50 | 10 000 | (0.9092) | 0.9192 | 37.37* | 0.03 | $7.896 \times 10^{-4}$ |
| $\mathbf{G}_{0.1}$ vs. $\mathbf{G}_Q$,<br>same evecs | 5 | 3600 | 0.9090 | — | 1.34 | 0.00 | $7.897 \times 10^{-4}$ |
| | 20 | 2100 | 0.7306 | — | 0.48 | 0.01 | $6.966 \times 10^{-4}$ |
| | 50 | 9800 | 0.5720 | — | 35.68 | 0.04 | $8.863 \times 10^{-4}$ |
| $\mathbf{G}_{0.5}$ vs. $\mathbf{G}_Q$,<br>same evecs | 5 | 3700 | 0.8369 | — | 1.47 | 0.00 | $1.138 \times 10^{-3}$ |
| | 20 | 4000 | (0.5286) | 0.5308 | 2.26 | 0.01 | $1.466 \times 10^{-3}$ |
| | 50 | 10 000 | (0.3589) | 0.3657 | 37.41* | 0.04 | $1.337 \times 10^{-3}$ |
| Two $\mathbf{G}_{0.1}$,<br>different evecs | 5 | 190 | 0.8143 | — | 0.02 | 0.00 | $1.091 \times 10^{-3}$ |
| | 20 | 1800 | 0.5141 | — | 2.96 | 0.01 | $1.831 \times 10^{-3}$ |
| | 50 | 9900 | 0.3269 | — | 830.93 | 0.03 | $1.856 \times 10^{-3}$ |
| Two $\mathbf{G}_{0.5}$,<br>different evecs | 5 | 2900 | 0.4254 | — | 1.02 | 0.00 | $3.076 \times 10^{-3}$ |
| | 20 | 4000 | (0.1559) | 0.1585 | 15.49 | 0.01 | $2.098 \times 10^{-3}$ |
| | 50 | 10 000 | (0.0715) | 0.0767 | 845.44* | 0.03 | $1.457 \times 10^{-3}$ |
| Two $\mathbf{G}_Q$,<br>different evecs | 5 | 3900 | 0.6644 | — | 1.88 | 0.00 | $2.239 \times 10^{-3}$ |
| | 20 | 430 | 0.5816 | — | 0.18 | 0.01 | $1.295 \times 10^{-3}$ |
| | 50 | 180 | 0.5657 | — | 0.37 | 0.03 | $7.993 \times 10^{-4}$ |
| $\mathbf{G}_{0.1}$ vs. $\mathbf{G}_{0.5}$,<br>different evecs | 5 | 2700 | 0.5904 | — | 0.87 | 0.00 | $2.056 \times 10^{-3}$ |
| | 20 | 4000 | (0.2835) | 0.2856 | 15.03 | 0.01 | $1.936 \times 10^{-3}$ |
| | 50 | 10 000 | (0.1532) | 0.1584 | 845.41* | 0.04 | $1.543 \times 10^{-3}$ |
| $\mathbf{G}_{0.1}$ vs. $\mathbf{G}_Q$,<br>different evecs | 5 | 3500 | 0.7293 | — | 1.50 | 0.00 | $1.729 \times 10^{-3}$ |
| | 20 | 1800 | 0.5435 | — | 2.91 | 0.01 | $1.647 \times 10^{-3}$ |
| | 50 | 9200 | 0.4293 | — | 717.31 | 0.04 | $1.684 \times 10^{-3}$ |
| $\mathbf{G}_{0.5}$ vs. $\mathbf{G}_Q$,<br>different evecs | 5 | 3600 | 0.5276 | — | 1.57 | 0.00 | $2.636 \times 10^{-3}$ |
| | 20 | 4000 | (0.2979) | 0.3001 | 15.13 | 0.01 | $1.939 \times 10^{-3}$ |
| | 50 | 10 000 | (0.2009) | 0.2077 | 845.67* | 0.03 | $1.524 \times 10^{-3}$ |

**Figure 4.**
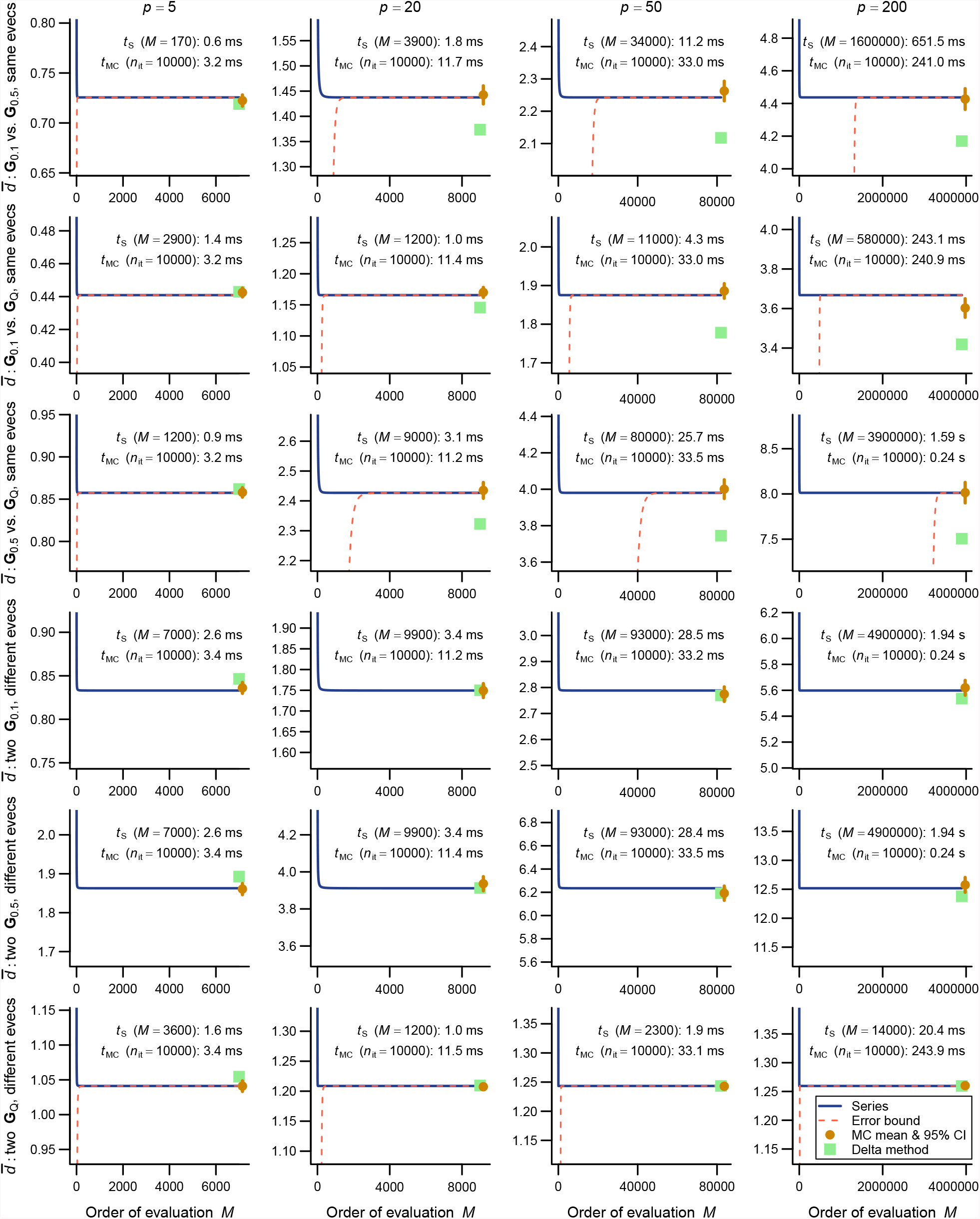
Results of numerical experiments for 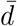. Top three rows are comparisons between covariance matrices with different eigenvalues and same eigenvectors, whereas the bottom three rows are between matrices with same eigenvalues and different eigenvectors. Comparisons between different eigenvalues and different eigenvectors are not shown for space limitations. See Figure 2 for legend. Error bounds are not available in some same-eigenvalue comparisons because of the singularity of argument matrices (see text).

**Figure 5.**
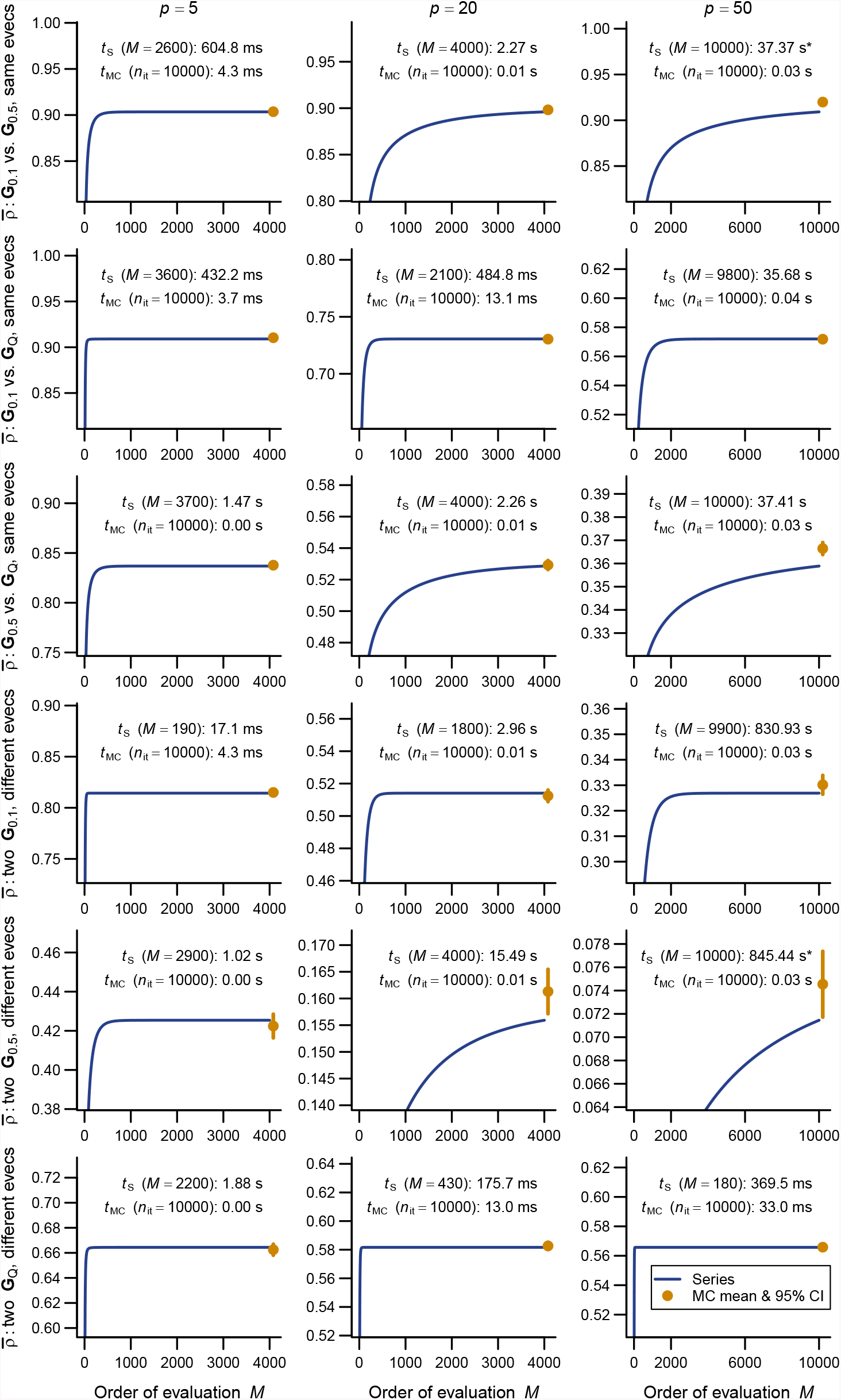
Results of numerical experiments for 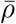. Top three rows are comparisons between covariance matrices with different eigenvalues and same eigenvectors, whereas the bottom three rows are between matrices with same eigenvalues and different eigenvectors. Comparisons between different eigenvalues and different eigenvectors are not shown for space limitations. See Figure 2 for legend. Delta method approximation or error bound is not available for 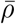. Asterisks after computational times denote ones from single runs rather than from 10 runs.

As expected, the Monte Carlo evaluation always yielded an unbiased estimate with the confidence intervals having the intended nominal coverage of the converged series where available. As implemented in the `qfratio` package, iterations in the order of 10^4^ can be executed within a second, so might be favored over the new series expression when the computational burden is a constraint and stochastic error is tolerated. Interestingly, relative accuracies of the series and Monte Carlo evaluations showed a qualitatively similar pattern of variation across eigenvalue conformations for a given average measure and *p*; the Monte Carlo evaluation tended to yield a large SE when the series evaluation or its error bound was slow to converge (Figs. 2–4). In other words, eigenvalue conformations seem to largely determine the accuracy at which average measures can be empirically evaluated.

The delta method approximations by Hansen & Houle (2008, 2009) can be evaluated almost instantaneously (from ∼0.1 ms to ∼7 ms in *p* = 5 and 200), as they are straightforward arithmetic operations once eigenvalues of the argument matrix are obtained. For 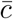 in large *p*, they showed notable empirical accuracy (Fig. 2; Table 1) (see also Hansen & Houle, 2008). In many other cases, however, they yielded inaccurate values, often deviating from the converged series by >5%, and it is even difficult to discern a consistent trend in the directions of errors (Figs. 2–4; Tables 2, 4, 5). Except in the former cases, it is rather questionable when those approximations can be reliably used.

## 4 Discussion

The present paper derived new expressions for average evolvability measures using results on the moments of ratios of quadratic forms in normal variables. These series expressions are in theory exact, although for practical evaluation summation must be truncated, yielding truncation error. A great advantage in the present approach is that a hard upper bound for the truncation error is known for some of the measures: namely, average conditional evolvability 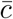, average respondability 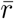, average flexibility 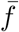, and average response difference 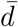 (above). This is unlike the confidence intervals from Monte Carlo evaluations, which always involve some uncertainty. In addition, empirically speaking, evaluation of most of the average measures, excepting average autonomy *ā* and average response correlation 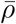, can be faster than Monte Carlo evaluation for modest-sized problems (e.g., *p* ≤ 50), depending on the accuracy aimed at. The accuracy and speed of the series evaluation will facilitate practical studies that involve quantification and/or comparison of evolvability or integration using average measures, for which the Monte Carlo evaluation or delta method approximation has traditionally been used.

As a practical guide for calculating average measures, the series evaluation is recommended whenever numerical convergence can be reached within reasonable computational time and memory. Its feasibility largely depends on the eigenvalues of **G**, but it is always easy to check whether convergence has been reached by inspecting a profile of the partial sum and by using error bounds when available. When the computational time and/or memory is a constraint (for *ā* and 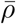 in large *p* with many small eigenvalues), then an alternative will be Monte Carlo evaluation, which is adequate because the evolvability measures considered herein have finite second moments (Appendix B). The delta method approximations as proposed by Hansen & Houle (2008, 2009) can yield rather inaccurate values with hardly predictable errors (above), thus will not remain a viable option in estimating average measures. This is perhaps except when even the (typically sub-second) computational burden of the Monte Carlo evaluation is a bottleneck, e.g., when one has large Bayesian samples of **G**. However, a primary problem in the delta method remains that, although the method empirically works well in certain situations (e.g., 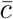 in large *p*), there is no theoretical guarantee for it to work well in general situations; the accuracy always need to be assessed by another method of evaluation.

It should be remembered that, in empirical analyses, **G** is almost always estimated with uncertainty, so that estimates of (average) evolvability measures involve error and potentially bias (Houle & Meyer, 2015; Grabowski & Porto, 2017). Although uncertainty may partially be incorporated in a Bayesian sampling of **G** (Aguirre *et al*., 2014) or a parametric-bootstrap-like sampling with multivariate normal approximation (Houle & Meyer, 2015), these methods will not by themselves deal with potential estimation bias. The present series expressions are not free from this problem, which need to be addressed in future investigations. It might be possible to derive sampling moments of the (average) measures under simple sampling conditions (e.g., Wishartness), as was recently done for eigenvalue dispersion indices of covariance and correlation matrices (Watanabe, 2022). A potentially promising fact is that the expectations of zonal and invariant polynomials of Wishart matrices are known (Constantine, 1963; Davis, 1980). It remains to be done to derive equivalent results to polynomials in more complicated functions of random matrices like the ones appearing in the present expressions. Apart from that, simulation-based approaches would also be useful in obtaining rough estimates of sampling error and bias (Haber, 2011; Grabowski & Porto, 2017). In any case, the present expressions will greatly facilitate future investigations into sampling properties of average measures as they provide quasi-exact values given (estimates of) **G**, unlike the approximate methods used in previous studies.

Although the present paper dealt with six evolvability measures, this is not to endorse all of them. Some of these measures may be disfavored from a biological or other theoretical ground. For example, Hansen & Houle (2008) criticized the use of (average) response correlation 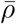 in the random skewers analysis for its insensitivity to potentially different magnitudes of response vectors and recommended (average) response difference 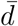 as an alternative. (To be fair, Cheverud (1996) argued for the use of 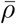 by claiming that there tends to be a large sampling bias in the magnitude of response vectors from empirical covariance matrices, but that remains to be justified.) Rohlf (2017) also pointed out difficulties in interpreting 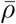 without knowledge on the eigenvalues and eigenvectors of the matrices compared, and recommended simply using those quantities for comparing matrices. The present author refrains from going into details on biological validity or utility of individual evolvability measures, because they will ultimately depend on the scope and context of individual studies as well as the nature of the data analyzed.

It seems worth repeating one of Rohlf’s (2017) insightful caveats here. Rohlf (2017) considered the angle between response vectors (arc cosine of response correlation *ρ*) from two **G** matrices in bivariate traits and empirically showed that the angle can have a bimodal distribution when **β** is distributed uniformly on the unit circle. The interpretation of the mean of such a multimodal distribution will not be straightforward, although, at least as empirically observed, the multimodality may not be a universal phenomenon (see Machado *et al*., 2018). At present, the possibility remains that other evolvability measures can have multimodal distributions as well. Generally speaking, it is known that the distribution of the ratio of quadratic forms **x**^*T*^**Ax**/**x**^*T*^**Bx**, where **B** is positive definite, has different functional forms across intervals bordered by eigenvalues of **B**^−1^**A** (Hillier, 2001; Forchini, 2002, 2005), around which distinct density peaks can potentially be observed. Formal evaluation of the conditions for multimodality seems rather tedious (if tractable at all) and is not pursued herein. It should at least be borne in mind that the average measures are insensitive to the potential multimodality and that, generally speaking, a scalar summary index like them can mask those nuanced aspects of the underlying distribution and the multivariate covariance structure.

The present paper suggested slight modifications for some evolvability measures to accommodate potentially singular **G** matrices. As originally defined by Hansen & Houle (2008), conditional evolvability *c* and autonomy *a* (and integration *i*) could not be evaluated for singular **G** as they required matrix inversion. This restriction can be released by using the generalized inverse as done above. Conditional evolvability *c* and autonomy *a* are zero unless **β** ∈ *R*(**G**). The corresponding average measures (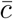 and *ā*) are also zero unless the same condition is satisfied with nonzero probability (i.e., when **β** is continuously distributed, its distribution is to be restricted to *R*(**G**)), as partially surmised by Hansen & Houle (2008). The new expressions enable to calculate *c* (4), *a* (6), *i* and their averages (52), (53) even when **G** is singular, as long as **β** ∈ *R*(**G**). Biologically speaking, the singularity of **G** represents the presence of absolute genetic constraint, where no genetic variance is available in certain directions. Although detection of absolute constraint in empirical systems requires different frameworks than evolvability measures (Mezey & Houle, 2005; Hine & Blows, 2006; Meyer & Kirkpatrick, 2008; Pavlicev *et al*., 2009), the present development will facilitate investigations into systems with absolute constraints. These will also involve, e.g., shape variables that lack variation in certain directions in the full (nominal) trait space (see below). However, it is necessary to distinguish the singularity in a true **G** matrix and that in its empirical estimates (**Ĝ**, say) due to small sample size. It should be viewed with much caution whether evolvability measures can be meaningfully applied to the latter, because *R*(**Ĝ**) as a space fluctuates due to random sampling when the sample size is not large enough to span the entire *R*(**G**).

Previous treatments of evolvability measures (Hansen & Houle, 2008; Kirkpatrick, 2009; Marroig *et al*., 2009; Bolstad *et al*., 2014) and random skewers (Revell, 2007) almost exclusively considered the simple condition where selection gradients are spherically distributed, **β** ∼ *N*_*p*_(**0**_*p*_, *σ*^2^**I**_*p*_), or uniformly distributed on a unit hypersphere. As detailed in Appendix C, the present results can be extended to a more general condition where the selection gradient has an arbitrary normal distribution, **β** ∼ *N*_*p*_(**η, Σ**). These extended results can potentially be useful when interest is in characterizing and comparing evolvability of populations under a certain directional and/or correlated selection regime. Of course, the original expressions of evolvability measures for a fixed **β** can be used when the selection is considered deterministic, so the utility of the new extended expressions will be in when the selection is considered random but potentially directional and/or correlated.

Chevin (2013) theoretically investigated the rate of adaptation (change in log average fitness across generations) in the same condition where **β** has a general normal distribution. He notably showed that, when stabilizing selection is assumed to be weak, the rate of adaptation is approximately equal to **β**^*T*^**Gβ**. He called this quantity “evolvability”, ascribing it to Hansen & Houle (2008), and discussed at length on its interpretation. However, that quantity should be distinguished from the evolvability *e* as defined by Hansen *et al*. (2003b) and Hansen & Houle (2008), which is standardized by the norm of the selection gradient **β**.^4^ This standardization can also be seen as projection of the selection gradients onto a unit hypersphere (Mardia & Jupp, 1999); directional and correlational selections can be expressed as concentration of density on the hypersphere. Accordingly, Chevin’s quantity and the present expressions for evolvability measures under general normal selection regimes have different interpretations and utility. The former would be more appropriate as a measure of absolute rate of adaptation, whereas the latter seems to be more appropriate for characterizing **G** matrices independently of the magnitude of selection.

The present expressions for a general normal case would be particularly useful when the selection gradients are assumed to be restricted to a certain subspace rather than distributed across the entire trait space, i.e., when **Σ** is singular. Such restriction can potentially arise from biological reasons, but probably are more often encountered from structural constraints in certain types of traits, e.g., shape variables. When the support of the (nominal) traits is restricted to a certain subspace, it would be sensible to assume that selection is also restricted to the same subspace (e.g., when geometric shapes are concerned, it is senseless to consider the components of selection corresponding to geometric rotation or translocation). Along with the accommodation of potentially singular **G** matrices, this extension allows for application of average measures to various types of traits encountered in the current evolutionary biology. Overall, the present development will facilitate theoretical and empirical investigations into the evolution of multivariate characters, by providing accurate means to quantify and compare various aspects of evolvability, integration, and/or constraint.

## 5 Acknowledgements

The author would like to thank Thomas F. Hansen and Diogo Melo for constructive comments. This work was supported by the Newton International Fellowships by the Royal Society (NIF\R1\180520) and the Overseas Research Fellowships by the Japan Society for the Promotion of Science (202160529).

## Appendix A Zonal and invariant polynomials

This paper evaluates the expectations of ratios of quadratic forms using the zonal and invariant polynomials, mainly following Smith (1989) and Hillier *et al*. (2009, 2014). These polynomials are widely used in the literature of multivariate distribution theory, primarily to integrate out components affected by orthogonal rotations from functions of random matrices. To be precise, they are not strictly required to derive results relevant to the present applications (see Bao & Kan, 2013), but allow for a conceptually elegant derivation and provide alternative interpretations of some of the results as hypergeometric functions of matrix argument. A brief introduction to these mathematical toolkits is provided in this section, as they appear to have attained little use in the biological literature. For more details, readers are directed to, e.g., James (1964), Muirhead (1982, 2006), Takemura (1984), Gross & Richards (1987), Mathai *et al*. (1995), and Chikuse (2003, appendix A).

### A.1 Notations

In this section, **Y** denotes a *p* × *p* symmetric matrix with eigenvalues *y*_1_, *y*_2_, …, *y*_*p*_. **H** denotes a *p* × *p* orthogonal matrix, and the space of *p* × *p* orthogonal matrices is denoted by *O*(*p*).

For indexing polynomials, (ordered) partitions *κ* of positive integers *k* are used; *κ* = (*k*_1_, *k*_2_, …), where *k*_*i*_ are nonnegative integers that satisfy *k*_1_ ≥ *k*_2_ ≥ … and _*i*_ *k*_*i*_ = *k*. For instance, the possible partitions of *k* = 4 are (4), (3, 1), (2, 2), (2, 1, 1), and (1, 1, 1, 1). The partition of *k* with a single nonzero integer part is called the top-order partition of *k*, and is denoted by [*k*].

The Pochhammer’s symbol is

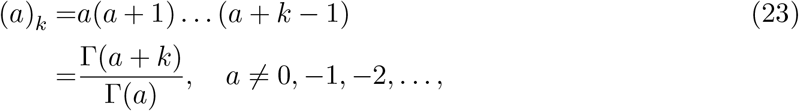

where Γ(·) is the gamma function, with the understanding (*a*)_0_ = 1. A multivariate extension (*a*)_*κ*_ (indexed by a partition) is defined as

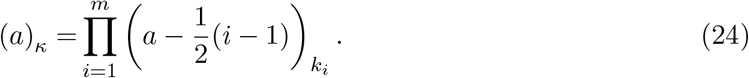

### A.2 Definitions of zonal polynomials

The zonal polynomials are a certain class of polynomials in matrix entries which has several useful properties. The primary definition of the zonal polynomials comes from the group representation theory (James, 1960, 1961, 1964), which tends to be too abstract to comprehend for anyone new to the concept. There are alternative, more operational definitions of the zonal polynomials (Muirhead, 1982; Takemura, 1984), which would be more approachable to most biologists. However, those alternative definitions do not generalize into the invariant polynomials of multiple matrix arguments, which are also relevant to the present application. For this reason, the representation-theoretic definition is briefly outlined herein, omitting technical details. See Gross & Richards (1987) for a readable and sufficiently comprehensive introduction to the topic.

Consider the vector space *V*_*k*_ of homogeneous polynomials *φ*(**Y**) of degree *k* in the elements of the symmetric matrix **Y**. The transformation with a *p* × *p* invertible matrix **L**,

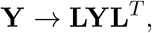

transforms the vector space by

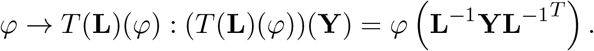

(Technically, *T*(**L**) here is a representation, which abstracts the transformation of the right-hand side by a linear transformation.) It is known that the vector space *V*_*k*_ decomposes into a direct sum of irreducible invariant subspaces *V*_*κ*_ as

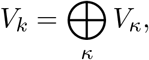

where the direct summation runs over all partitions *κ* of *k* into not more than *p* parts. By *V*_*κ*_ being invariant, it is meant that *T*(**L**)*V*_*κ*_ ⊂ *V*_*κ*_ for any invertible **L**, and by them being irreducible invariant, it is meant that they do not contain any proper invariant subspace by themselves. It is further known that each *V*_*κ*_ contains a unique one-dimensional subspace that is invariant under **Y** → **HYH**^*T*^ (restricting the above transformation to **L** ∈ *O*(*p*)). Then, [tr (**Y**)]^*k*^ ∈ *V*_*k*_ has a unique decomposition

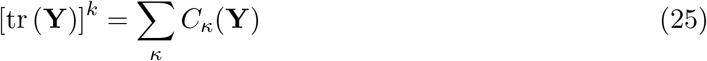

into some polynomials *C*_*κ*_(**Y**) ∈ *V*_*κ*_ which satisfy *C*_*κ*_(**Y**) = *C*_*κ*_(**HYH**^*T*^). These polynomials *C*_*κ*_(**Y**) are called the zonal polynomials corresponding to the partitions *κ*. In short, the zonal polynomials of degree *k* constitute a basis of the subspace of *V*_*k*_ that is invariant under the transformation **Y** → **HYH**^*T*^, each corresponding to the irreducible subspace represented by a partition *κ* of *k*. In terms of (25), they can be regarded as a generalization of powers of scalars into matrices.

These polynomials can be scaled by arbitrary constants, and there exist several different normalizations in the literature. The normalization used in (25), with all coefficients for *C*_*κ*_(**Y**) being unity, is primarily for convenience of notation.

### A.3 Basic properties of zonal polynomials

A consequence of the above definition with (25) is that *C*_*κ*_(**Y**) are polynomials in the eigenvalues *y*_1_, …, *y*_*p*_ of **Y** (note that tr (**Y**) = tr (**HYH**^*T*^) = ∑_*i*_ *y*_*i*_). It also entails that they are symmetric against permutation of the eigenvalues. Alternatively, or equivalently, they can be expressed as polynomials in the traces of matrix powers tr 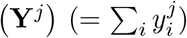. Unfortunately, no closed-form expressions are known for the zonal polynomials, except for a few special cases, so the coefficients in these polynomials in general need to be determined recursively (see Muirhead, 1982, 2006).

The zonal polynomials are originally defined for symmetric matrices, but the definition can be extended. If **X** is positive definite and **Y** is symmetric, the eigenvalues of **XY** and **X**^1/2^**YX**^1/2^ are the same. Therefore, a zonal polynomial of **XY** is defined as *C*_*κ*_(**XY**) = *C*_*κ*_(**X**^1/2^**YX**^1/2^).

Properties of the zonal polynomials have been extensively investigated (Constantine, 1963; James, 1964; Muirhead, 1982). Elementary properties involve the following:

- *C*_*κ*_(*a***Y**) = *a*^*k*^*C*_*κ*_(**Y**) holds for any constant *a*.
- If the number of parts in *κ* exceeds the rank of **Y**, *C*_*κ*_(**Y**) = 0 (Muirhead, 1982, corollary 7.2.4(i)).
- If **Y** is positive definite, *C*_*κ*_(**Y**) > 0 (Muirhead, 1982, corollary 7.2.4(ii)).
- *C*_*κ*_(**Y**) = *C*_*κ*_(diag (**Y, 0**)), where the right-hand side is with respect to a block diagonal matrix with a square matrix of 0’s of arbitrary dimension.

One of the most important properties of the zonal polynomials is the following integral identity.

#### Proposition 2

(James (1960)). *Let* **A** *and* **Y** *be p* × *p symmetric matrices. Then*,

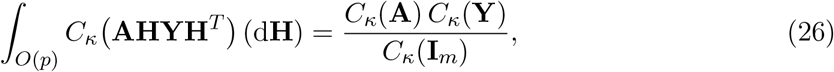

*where* (d**H**) *is the normalized invariant measure on O*(*p*) *that satisfies* ∫_*O*(*p*)_(d**H**) = 1.

Accessible proofs are given in Muirhead (1982, theorem 7.2.5) and Gross & Richards (1987, proposition 5.5).

### A.4 Top-order zonal polynomials

The zonal polynomials corresponding to the top-order partition, [*k*] = (*k*), are called the top-order zonal polynomials and appear frequently in the multivariate distribution theory. An explicit expression of the top-order zonal polynomials is (James, 1964, (123)):

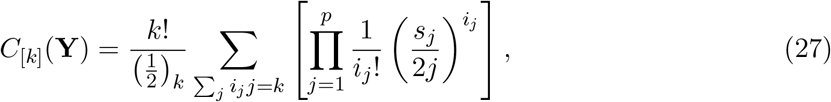

where 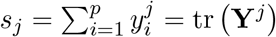 and the summation is over all such combinations of the nonnegative integers *i*_*j*_ that satisfy 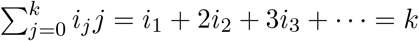. An alternative expression is given in Gross & Richards (1987, (6.8.1)).

For the identity matrix **I**_*p*_, the expression simplifies as:

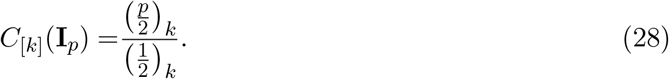

### A.5 Hypergeometric functions of matrix argument

With the zonal polynomials as a generalization of powers into matrices, the hypergeometric functions of matrix argument are defined as

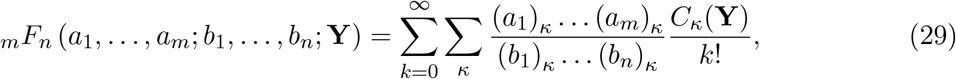

where the summation ∑_*κ*_ is across all ordered partitions *κ* of *k*, and (*x*)_*κ*_ is defined as in (24). Conditions for convergence are as follows: (i) when *m* ≤ *n*, the series converges for all **Y**; (ii) when *m* = *n* + 1, the series converges for all ∥**Y**∥ < 1, where ∥**Y**∥ is the maximum of the absolute values of the eigenvalues; (iii) when *m* > *n* + 1, the series diverges for all **Y** ≠ 0, unless the series terminates (when any *a*_*i*_ is a negative integer). The following paragraphs list a few results relevant to the present applications. More extensive treatments on this topic can be found in, e.g., Muirhead (1982), Gross & Richards (1987), and Chikuse (2003).

If any of *a*_*i*_ is 1/2, the series can be expressed as a weighted sum of top-order zonal polynomials, as (1/2)_*κ*_ = 0 unless *κ* = [*k*] (see (24)):

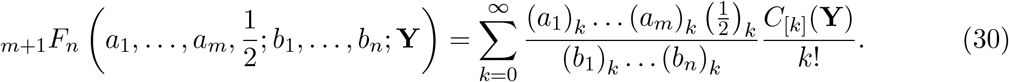

Analogous to the scalar equivalent, _0_*F*_0_ is an exponential series but about the trace:

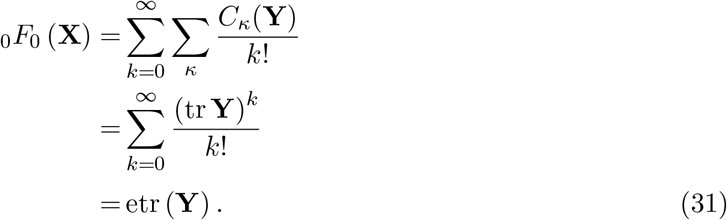

Provided that all eigenvalues of **Y** are less than 1 in absolute values, _1_*F*_0_ is a generalization of the binomial series (Muirhead, 1982, corollary 7.3.5):

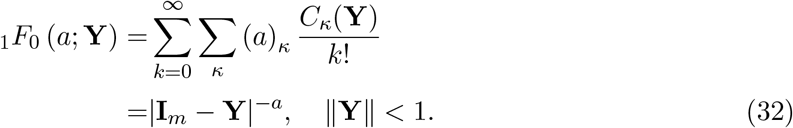

### A.6 Invariant polynomials of multiple matrix arguments

The invariant polynomials of multiple matrix arguments provide some extension of the zonal polynomials into multiple matrices (Davis, 1979, 1980; Chikuse, 1980; Chikuse & Davis, 1986a). Accessible introductions to the topic are found in, e.g., Chikuse & Davis (1986b) and Mathai *et al*. (1995, appendix).

Let 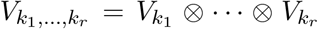 be the vector space of polynomials of degree *k*_1_, …, *k*_*r*_ in the elements of *p* × *p* symmetric matrices **Y**_1_, …, **Y**_*r*_, respectively. Consider the following simultaneous transformation with a *p* × *p* invertible matrix **L**:

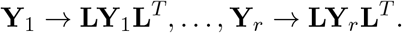

First noting the subspaces in 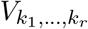 that are invariant against this transformation and then by decomposing them into irreducible ones, one has the following decomposition:

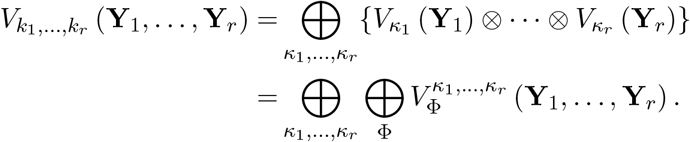

Here, 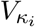 are analogous to *V*_*κ*_ discussed above but for **Y**_*i*_, and 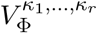 are irreducible invariant subspaces indexed by Φ and *κ*_1_, …, *κ*_*r*_. Φ are certain ordered partitions of 2*f*, with *f* = *k*_1_ + · · · + *k*_*r*_, into not more than *p* parts, whose range is determined by *κ*_1_, …, *κ*_*r*_. *κ*_*i*_ run across partitions of *k*_*i*_, *i* = 1, …, *r*, into not more than *p* parts. It is known that only those 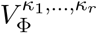 with partitions taking the form Φ = 2*ϕ*, where *ϕ* is a partition of *f*, contain a subspace invariant under the simultaneous transformation **Y**_1_ → **HY**_1_**H**^*T*^, …, **Y**_*r*_ → **HY**_*r*_**H**^*T*^, which is one-dimensional. The corresponding polynomials are called the invariant polynomials, 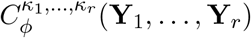, indexed by *κ*_1_, …, *κ*_*r*_ and *ϕ* ∈ *κ*_1_ · · · *κ*_*r*_. (There is a potential issue of non-uniqueness, but this is not concerned in most applications.)

Explicit forms of the invariant polynomials are even less obvious than the zonal polynomials, but they are polynomials in matrix traces of the form tr 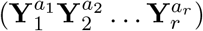, so that the total degrees in elements of **Y**_1_, …, **Y**_*r*_ are *k*_1_, …, *k*_*r*_, respectively.

An important generalization of Proposition 2 enabled by the invariant polynomials is the following (see also Mathai *et al*., 1995, theorem A.3).

#### Proposition 3

(Davis (1980), Chikuse (1980)). *Let* **A**_1_, …, **A**_*r*_ *and* **Y**_1_, …, **Y**_*r*_ *be p*×*p symmetric matrices, and κ*_1_, …, *κ*_*r*_ *be ordered partitions of k*_1_, …, *k*_*r*_ *into not more than p parts*.

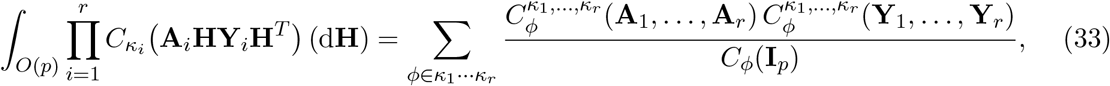

*where the summation runs across all permissible partitions ϕ of f* = *k*_1_ + · · · + *k*_*r*_ *corresponding to κ*_1_, …, *κ*_*r*_.

One notable corollary from this result is (Davis, 1980; Chikuse, 1980)

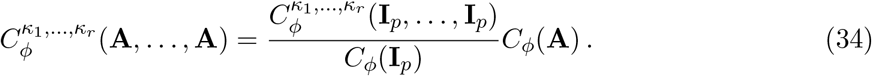

The top-order invariant polynomials, in which *ϕ* = [*f*], the top-order partition of *f*, are of particular importance here. This partition occurs only when *κ*_*i*_ = [*k*_*i*_] for all *i*. An explicit form of the top-order invariant polynomials is (Chikuse, 1987; Hillier *et al*., 2009):

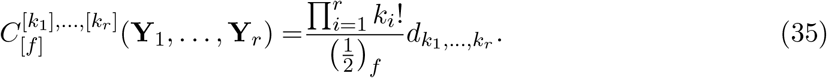

Here, 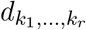 is defined as:

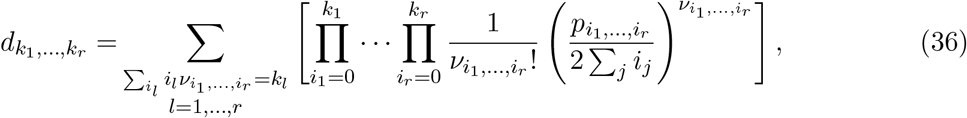

where *i*_*j*_ run from 0 to *k*_*j*_ (but at least one of *i*_*j*_ is nonzero), and 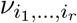 are nonnegative integers; these collectively satisfy the *r* linear constraints 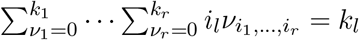 for *l* = 1, …, *r*.^5^

And 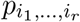 are the coefficients for 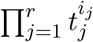 in the expansion of the following:

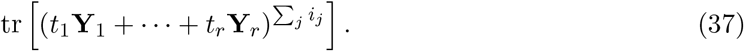

Note that (35) simplifies into (27) when *r* = 1. In addition, it holds that (Chikuse, 1987)

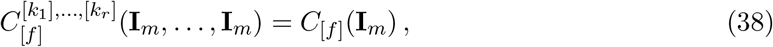

regardless of individual values of *k*_*i*_ (see also (28)).

### A.7 Example application to evolvability measures

As a demonstration for how above mathematical toolkits act in the present problem of evaluating moments of (multiple) ratios of quadratic forms, an expression of the average autonomy *ā* is derived here, under the simplest case that **β** ∼ *N*_*p*_(**0**_*p*_, *σ*^2^**I**_*p*_), where *σ*^2^ is an arbitrary positive constant, and **G** is nonsingular. The derivation here is a simplified version of that in Smith (1989). When **G** is singular, the approach of Bao & Kan (2013) can be used instead.

As discussed in the text, **u** is uniformly distributed on the unit hypersphere *S*^*p*−1^ under this assumption. Thus, the average autonomy *ā*, from the definition in (5), is to be evaluated as the following integral:

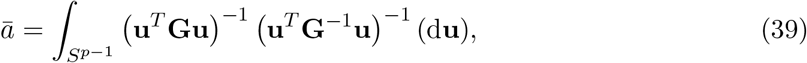

where (d**u**) denotes the normalized uniform measure on *S*^*p*−1^ (see, e.g. Muirhead, 1982). To evaluate this, consider the following transformation:

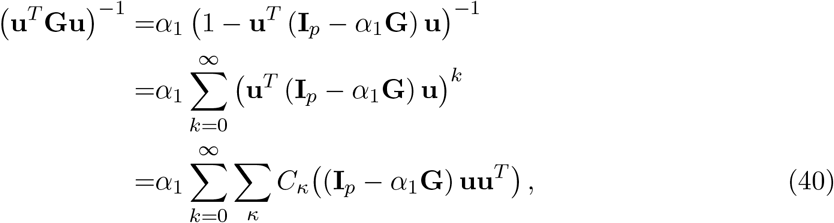

where *α*_1_ is a positive constant that satisfy 0 < *α*_1_ < 2/*λ*_max_(**G**) (so that |**u**^*T*^ (**I**_*p*_ − *α*_1_**G**) **u**| < 1), the second equation is from the fact 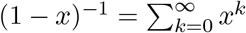 when |*x*| < 1, and the last equation is from (25) and the fact **u**^*T*^ (**I**_*p*_ − *α*_1_ **G**) **u** = tr (**I**_*p*_ − *α*_1_ **G**) **uu**^*T*^). Note that (40) can be written as _1_*F*_0_ (1; (**I**_*p*_ − *α*_1_ **G**) **uu**^*T*^ from (32) and is uniformly convergent under the assumptions. (This series does not converge when **G** is singular.) By similar transformations, 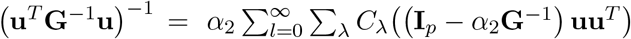, where 0 < *α*_2_ < 2/*λ*_max_(**G**^−1^). Then, the above integral becomes:

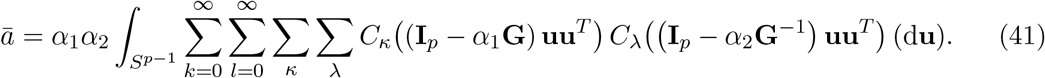

Here, term-by-term integration is permissible because the series is uniformly convergent (see also Bao & Kan, 2013).

Note that integration of **u** over *S*^*p*−1^ is equivalent to that of **H** over *O*(*p*) (e.g., Mardia & Jupp, 1999), whence the change of variables **u** = **He** is possible, where **e** = (1, 0, … 0)^*T*^ (or **e** could be any unit vector). By denoting **ee**^*T*^ = **E**, the integral is now

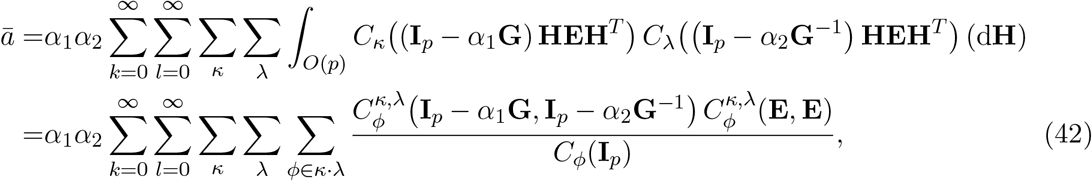

by applying Proposition 3. Furthermore, using (34), 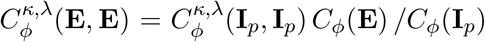. However, *C*_*ϕ*_(**E**) is nonzero only if *ϕ* = [*k* + *l*], the top-order partition, because **E** has only one nonzero eigenvalue, which is unity. Hence, the summations for *κ, λ, ϕ* disappear, leaving only the terms corresponding to the top-order partitions, [*k*], [*l*], [*k* + *l*]. Also note that, from (38), 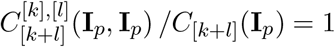. The integral finally becomes

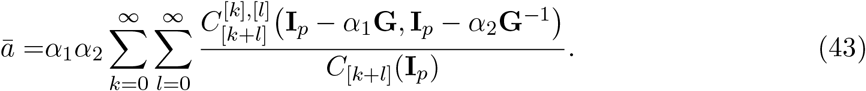

Inserting (28) yields (14) in the text. The expressions for all average measures presented in the main text under the same assumption can be derived in a similar manner.

### A.8 Generating functions for top-order zonal and invariant polynomials

For practical evaluation of top-order zonal and invariant polynomials, it is more efficient to handle 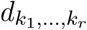 in (35) and (36) than the polynomials themselves. Chikuse (1987) showed that the same 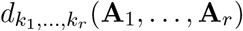 appear as coefficients in the following, which is the joint moment generating function of **x**^*T*^**A**_1_**x**/2, …, **x**^*T*^**A**_*r*_**x**/2 for **x** ∼ *N*_*p*_(**0**_*p*_, **I**_*p*_):

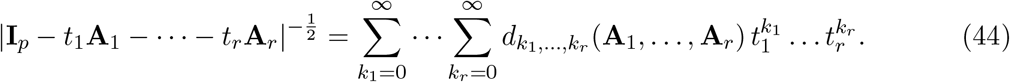

Chikuse (1987) and Hillier *et al*. (2009, 2014) presented a few recursive algorithms to evaluate 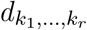.

Bao & Kan (2013) and Hillier *et al*. (2014) presented further improvements for evaluating moments of ratios of quadratic forms in potentially noncentral normal variables, **x** ∼ *N*_*p*_(**μ, I**_*p*_). Following those papers, the coefficients 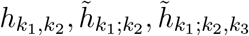 that appear in the following expansions are used in the present paper:

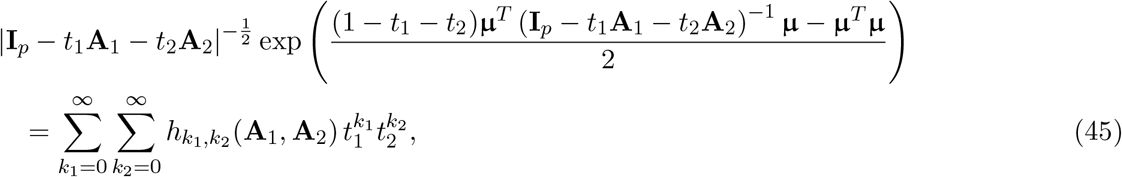

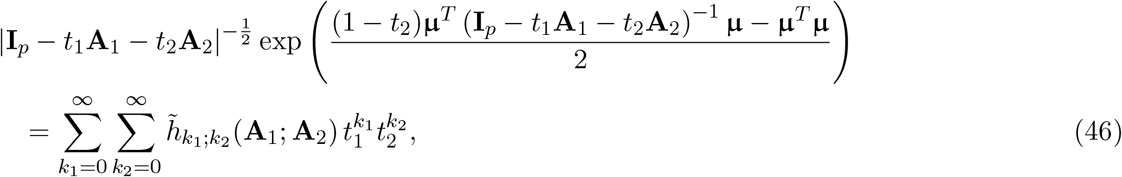

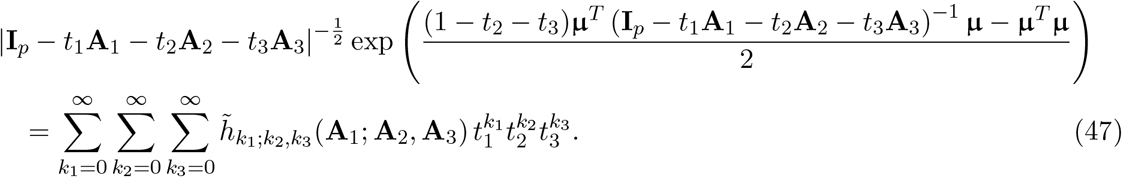

(45) and (46) are from Bao & Kan (2013), and (47) is a slight extension from (46) used in appendices B and C. All these simplify to (44) when **μ** = **0**_*p*_. Note that the first arguments in 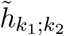 and 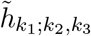 are not interchangeable with the other arguments, unlike those in 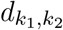 or 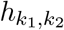.

## Appendix B Moment of multiple ratio of quadratic forms

Among the average evolvability measures mentioned in the text, the expressions for 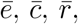, and 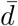 can be directly derived from Proposition 1. For *ā*, 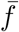, and 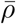, an expression is required for the expectation of the form 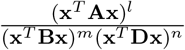; it suffices here to concern integer-valued *l*. Smith (1989) derived such an expression for **B, D** being positive definite. However, it is possible to derive the following, equivalent expression with a slightly general condition that **B, D** being nonnegative definite, following a similar line of arguments as in Bao & Kan (2013).

### Proposition 4

*Let* **x** ∼ *N*_*p*_ (**μ, I**_*p*_), **A** *be a symmetric p* × *p matrix*, **B, D** *be symmetric, nonnegative definite p* × *p matrices, l be a positive integer, and m, n be positive real numbers*.

*When the expectation of* 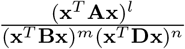 *exists, it is*

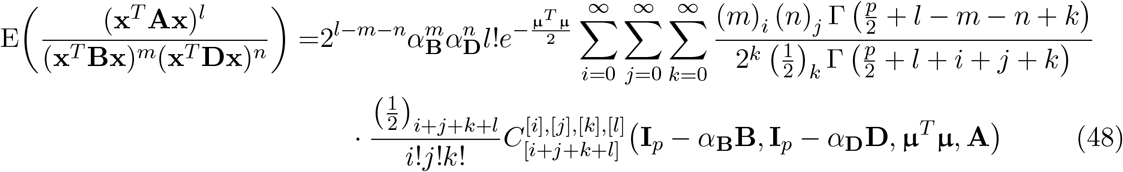

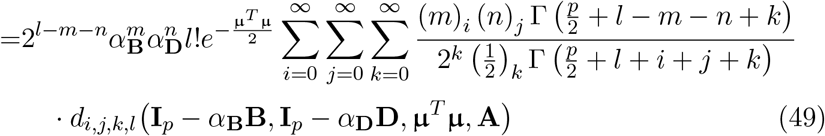

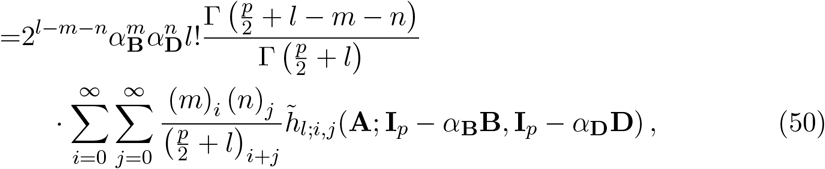

*where α*_**B**_, *α*_**D**_ *are to satisfy* 0 < *α*_**B**_ < 2/*λ*_max_(**B**) *and* 0 < *α*_**D**_ < 2/*λ*_max_(**D**).

Proof is omitted because it is almost identical to that of the proposition 3 in Bao & Kan (2013, pp. 237–243), but by using (47) in place of (45).

*Remark*. Conditions for the existence of the above moment were stated by Smith (1989) for positive definite **B, D**; in the notation here, the moment exists if and only if 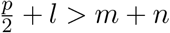. For positive semidefinite **B, D**, dealing with their null spaces is critical (see Roberts, 1995). Deriving general existence conditions in that case does not seem straightforward, so only the following simple cases are noted here. *(i) When N*(**D**) ⊆ *N*(**B**) *(or N*(**B**) ⊆ *N*(**D**)): In the former case, it is possible to find a positive constant *ϵ* that always satisfies (**x**^*T*^**Bx**)^*m*^(**x**^*T*^**Dx**)^*n*^ ≥ (*ϵ***x**^*T*^**Bx**)^*m*+*n*^, so that (**x**^*T*^**Ax**)^*l*^/[(**x**^*T*^**Bx**)^*m*^(**x**^*T*^**Dx**)^*n*^] ≤ *ϵ*^−(*m*+*n*)^(**x**^*T*^**Ax**)^*l*^/(**x**^*T*^**Bx**)^*m*+*n*^. The desired moment of the left-hand side exists if and only if that of the right-hand side does, and the necessary and sufficient condition for the latter is given in the proposition 1 of Bao & Kan (2013), which encompasses the positive definite case above. The same argument applies when *N*(**B**) ⊆ *N*(**D**) with obvious modifications. *(ii) When N*(**D**) ⊈ *N*(**B**) *and N*(**B**) ⊈ *N*(**D**): Consider the space *F* = {**y** : **y** ⊥ *N*(**B**), **y** ⊥ *N*(**D**), **y** ∈ ℝ^*p*^}. Assuming *F* ≠ ∅, it is possible to find a positive semidefinite matrix **F** that always satisfies (**x**^*T*^**Ax**)^*l*^/[(**x**^*T*^**Bx**)^*m*^(**x**^*T*^**Dx**)^*n*^] ≤ (**x**^*T*^**Ax**)^*l*^/(**x**^*T*^**Fx**)^*m*+*n*^, by choosing arbitrary small positive numbers and a basis of *F* as its nonzero eigenvalues and eigenvectors, respectively. The proposition 1 of Bao & Kan (2013) applied to the right-hand side defines a sufficient condition for the existence of the moment. This is admittedly a restrictive condition, and the existence condition in the case *F* = ∅ is not always obvious.

In the present applications, the moment existence conditions can be easily confirmed for *ē*, 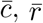, and 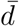 by directly applying the proposition 1 of Bao & Kan (2013). *ā* and 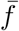 always fall within the case (i) above, and the moments exist as long as they are defined (in case of 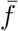, **β** should not be restricted onto *N*(**G**)). On the other hand, 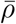 may fall within either (i) or (ii) depending on the structures of the two **G**’s. Even in the latter case and if *F* = ∅, which is a pathologic condition in this application, this specific moment exists as long as *ρ* is defined with probability 1, since −1 ≤ *ρ* ≤ 1 whenever defined (above). In addition, the same criteria can be used to confirm that these evolvability measures have finite second moments, which formally validate the use of Monte Carlo evaluations.

## Appendix C Evolvability measures in general normal cases

### C.1 Conditions considered

This section considers a general *p*-variate normality condition for the selection gradient, **β** ∼ *N*_*p*_ (**η, Σ**). Regarding ratios of quadratic forms, however, tractable results are available only for normal variables with a spherical covariance structure, i.e., *N*_*q*_ (**μ, I**_*q*_) for some *q* and arbitrary **μ**. Let **A** be an arbitrary *p* × *p* symmetric matrix and **K** be a *p* × *q* matrix that satisfies **KK**^*T*^ = **Σ**, where *q* (≤ *p*) is the rank of **Σ**. Writing **β** = **η** + **Kβ**_0_ with **β**_0_ ∼ *N*_*q*_(**0**_*q*_, **I**_*q*_), the quadratic form in **β** with respect to **A** can be written as **β**^*T*^**Aβ** = (**η** + **Kβ**_0_)^*T*^**A**(**η** + **Kβ**_0_) (see also Mathai & Provost, 1992, chapter 3). Certain assumptions are required to express this quadratic form as one in normal variables with a spherical covariance structure.

1. **When Σ is nonsingular** In this case, *p* = *q* and **K** is invertible. It suffices to consider the transformation **β**_*_ = **K**^−1^**β** ∼ *N*_*p*_ (**μ, I**_*p*_), where **μ** = **K**^−1^**η**. The quadratic form can be rewritten as 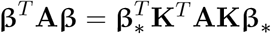, a new quadratic form in **β**_*_ with respect to **K**^*T*^**AK**, to which results below apply.
2. **When Σ is singular** In this case, **K** is not invertible and at least one of the following additional conditions need to be satisfied for the results below to be applicable. *(i)* **η** ∈ *R*(**Σ**) *(including* **η** = **0**_*p*_): Under this condition, it holds that **η** = **KK**^−^**η** (e.g., Schott, 2016, theorem 5.25) since *R*(**Σ**) = *R*(**K**). It is then possible to consider 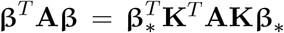, with **β**_*_ = **K**^−^**β** ∼ *N*_*p*_ (**μ, I**_*p*_), where **μ** = **K**^−^**η**, as a generalization of the nonsingular case. *(ii) R*(**A**) ⊆ *R*(**Σ**): From the symmetry of **A** and the fact that *R*(**K**) = *R* (K^−*T*^) it holds that **A** = **K**^−*T*^**K**^*T*^**AKK**^−^ under this condition (Schott, 2016, theorem 5.25). Then the same transformation as for *(i)* applies. (Utility of this condition in the present applications is relatively limited, since all evolvability measures except *ρ* involve at least one **I**_*p*_ in place of **A**.) *(iii)* **Aη** = **0**_*p*_: Under this condition, the above quadratic form simplifies into 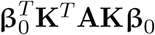, to which results below apply with **μ** = **0**_*q*_.

In practice, the conditions (1) or (2)(i) above seem to cover most cases of biological interest. That is, if the variation of selection gradient **β** is restricted to a certain subspace (condition (2)), then it seems plausible that the mean selection gradient **η** is also in the same subspace (condition (2)(i)). Otherwise, **β** ought to contain a nonzero deterministic component (**η**) that is strictly unaffected by (linearly independent of) its potential variability (represented by **Σ**).

### C.2 Average evolvability measures

Expressions for average evolvability measures for **β** ∼ *N*_*p*_ (**η, Σ**) or equivalently **β**_*_ ∼ *N*_*q*_ (**μ, I**_*q*_) can be derived from Propositions 1 and 4.

The average evolvability *ē* is

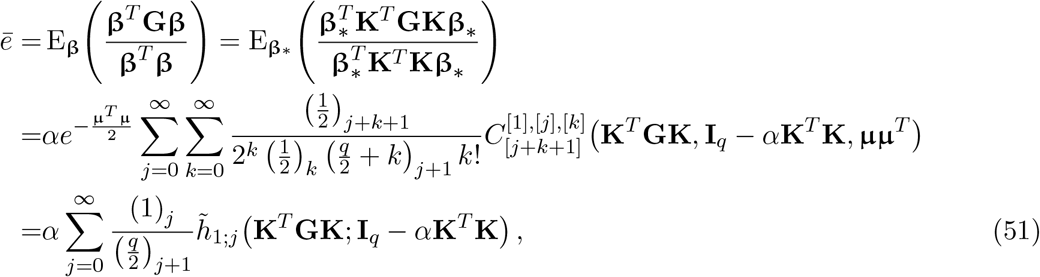

where *α* is a positive constant that satisfies 0 < *α* < 2/*λ*_max_(**K**^*T*^**K**).

As discussed in the text (2.4), the average conditional evolvability 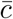 and average autonomy *ā* can be nonzero only if the distribution of **β** is strictly within *R*(**G**). This happens (i) when **G** is nonsingular; or (ii) when **G** is singular, **η** ∈ *R*(**G**) and *R*(**Σ**) ⊆ *R*(**G**).

Under these conditions, the following expression can be obtained for the average conditional evolvability 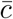:

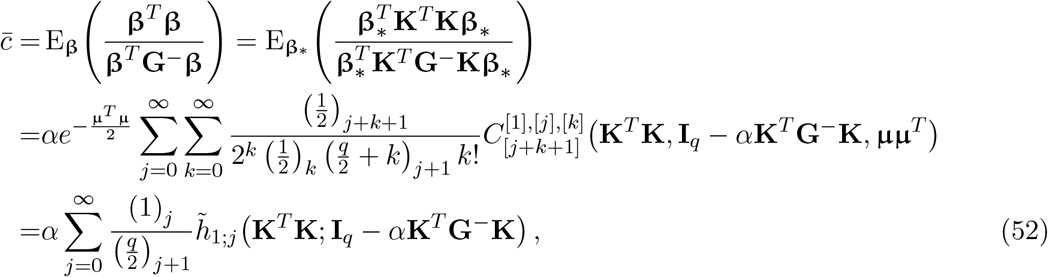

where *α* is to satisfy 0 < *α* < 2/*λ*_max_(**K**^*T*^**G**^−^**K**).

Under the same conditions, the average autonomy *ā* is

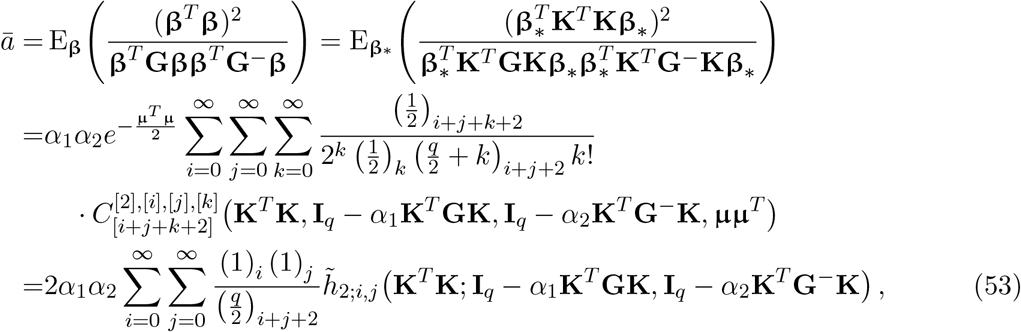

where *α*_1_ and *α*_2_ are to satisfy 0 < *α*_1_ < 2/*λ*_max_(**K**^*T*^**GK**)and 0 < *α*_2_ < 2/*λ*_max_(**K**^*T*^**G**^−^**K**). As before, the average integration *ī* is 1 − *ā*.

The average respondability 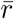 is

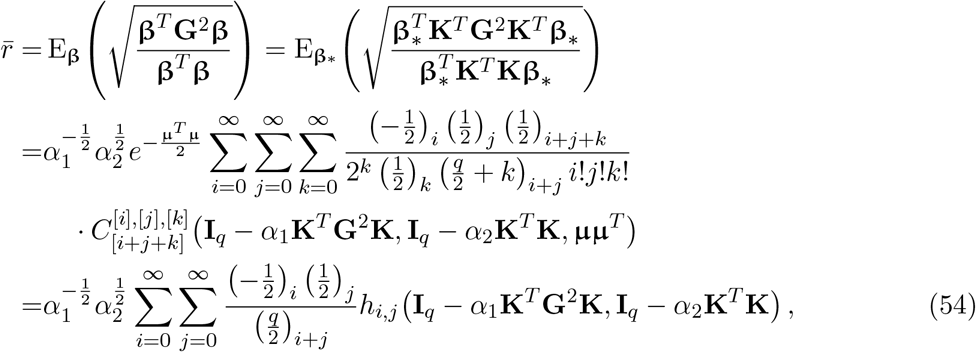

where *α*_1_ and *α*_2_ are to satisfy 0 < *α*_1_ < 2/*λ*_max_(**K**^*T*^**G**^2^**K**) and 0 < *α*_2_ < 2/*λ*_max_(**K**^*T*^**K**).

The average flexibility 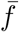 is

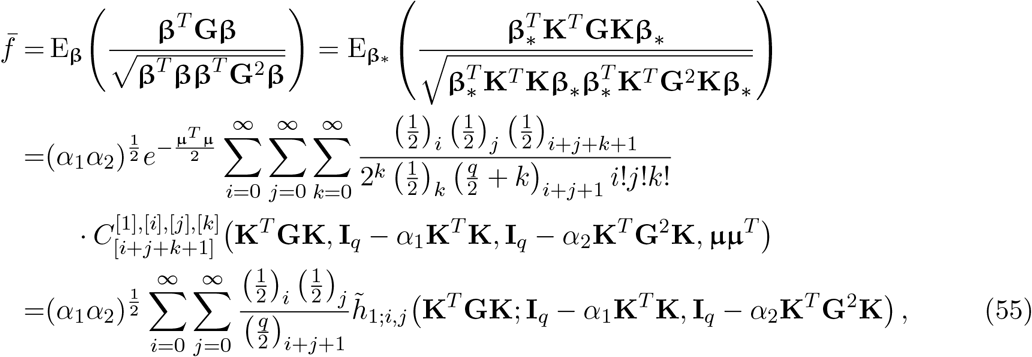

where *α*_1_ and *α*_2_ are to satisfy 0 < *α*_1_ < 2/*λ*_max_(**K**^*T*^**K**) and 0 < *α*_2_ < 2/*λ*_max_(**K**^*T*^**G**^2^**K**).

The average response difference 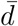 is

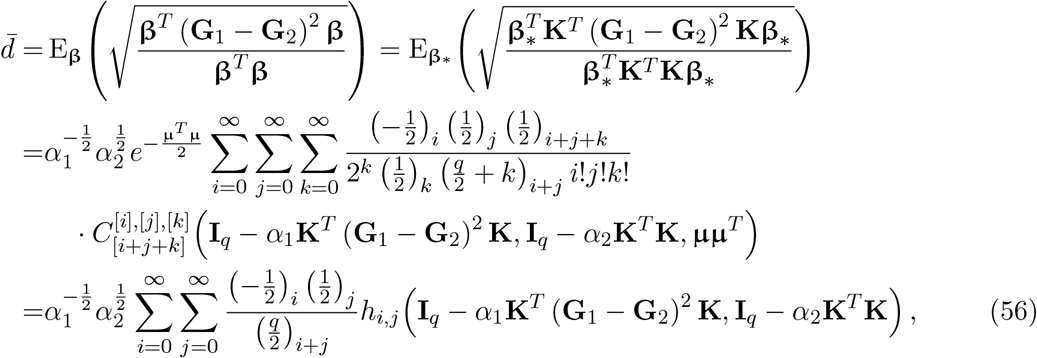

where *α*_1_ and *α*_2_ are to satisfy 0 < *α*_1_ < 2/*λ*_max_ (**K**^*T*^ (**G**_1_ − **G**_2_)^2^ **K**) and 0 < *α*_2_ < 2/*λ*_max_ (**K**^*T*^**K**).

The average response correlation 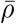 is

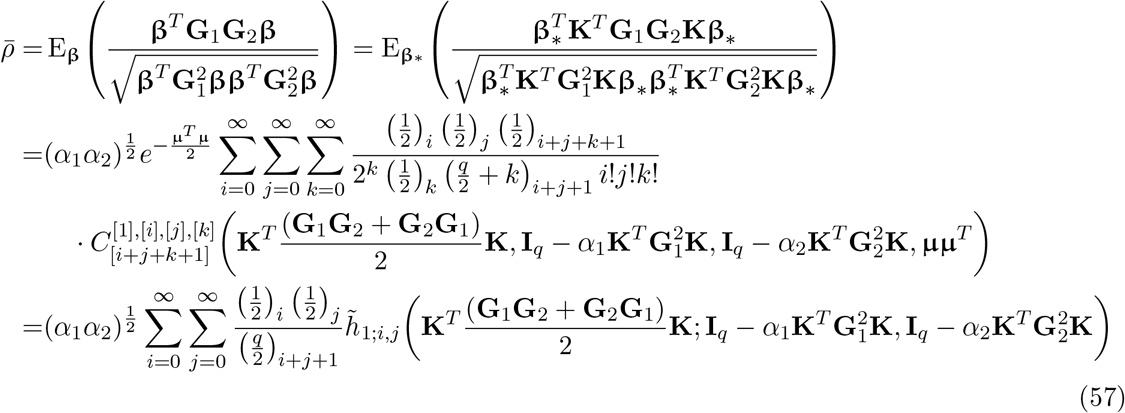

where *α*_1_ and *α*_2_ are to satisfy 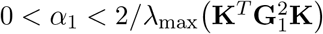 and 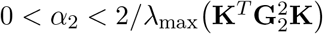.

### C.3 Practical evaluation and truncation error

The above series expressions for average measures in general cases can be evaluated by one of the recursive algorithms described by Hillier *et al*. (2009, 2014) and Bao & Kan (2013). When the mean **μ** is nonzero, the series can be most efficiently evaluated by recursions for *h*_*i,j*_ and 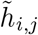 described in Hillier *et al*. (2014, section 5.4) and Bao & Kan (2013, section 5). When possible, it is recommended to inspect a profile of the partial sum across varying orders as the signs of the terms can in general fluctuate in those series.

At present, truncation error bounds for general cases are available only for *ē* and 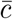. From the theorem 7 of Hillier *et al*. (2014), a truncation error bound for these average measures evaluated up to *k* = *M* is obtained from the following, by inserting (**A, B**) = (**K**^*T*^**GK, K**^*T*^**K**) and (**K**^*T*^**K, K**^*T*^**G**^−^**K**) for *ē* and 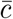, respectively:

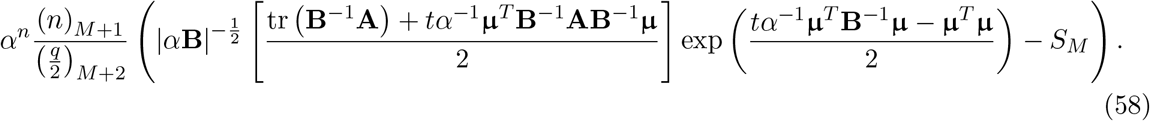

Here, *n* = 1, *t* = 2, and 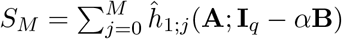, with 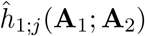 being the coefficient of 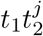 in the expansion

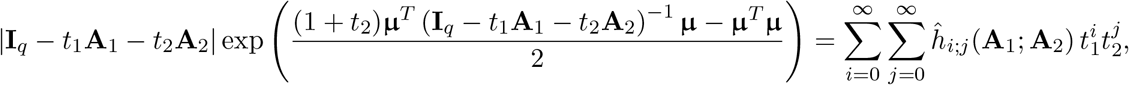

which can be evaluated by a recursive algorithm similar to the one presented in Hillier *et al*. (2014, theorem 6).^6^ This expression bounds the truncation error in absolute values unlike those in (19)–(22) which are one-sided. This is because 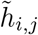 can take both positive and negative values when **μ** is nonzero, so that the partial sums are not always increasing.

Under the restriction **K**^*T*^**K** = **I**_*q*_, a similar error bound can be derived for 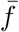, by letting **A** = **K**^*T*^**GK, B** = **K**^*T*^**G**^2^**K**, *n* = 1/2, *t* = 3, and 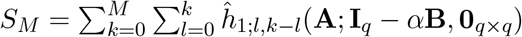 in (58). Here, 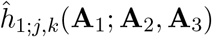 is the coefficient of 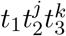 in the expansion of

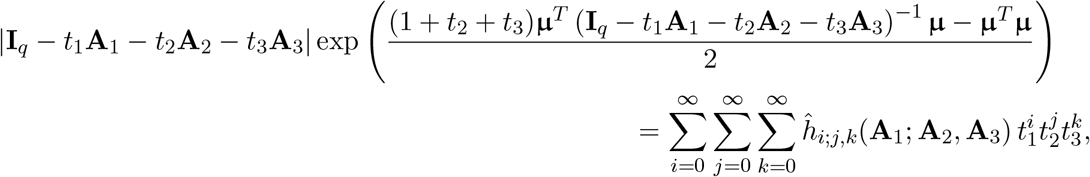

which can be evaluated by the same recursive algorithm for 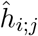 with obvious modifications.

## Appendix D Connections to previous results

### D.1 Average conditional evolvability in two traits

Hansen & Houle (2008) stated that the average conditional evolvability, under the uniform distribution of **u** on *S*^*p*−1^, equals to the geometric mean of the eigenvalues of **G** when *p* = 2. It is possible to confirm that the present result is consistent with their argument as follows. From (13),

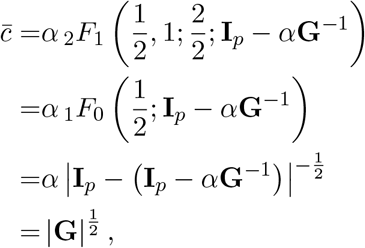

where in the second equation the denominator parameter cancels with one of the numerator parameters, and the third equation is from (32). Since the determinant is equal to the product of the eigenvalues, this proves the argument.

### D.2 Average respondability under complete integration

Kirkpatrick (2009) considered the average selection response 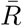, which equals 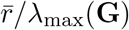(**G**) here when **u** is uniformly distributed on *S*^*p*−1^. This section shows that the present expression for 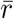 (15) entails his result for 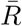 under the case of complete integration, **G** = diag (*λ*_max_, 0, …, 0), the only situation for which an analytic expression was given. The expression is (Kirkpatrick, 2009, (6) and (A7) therein)

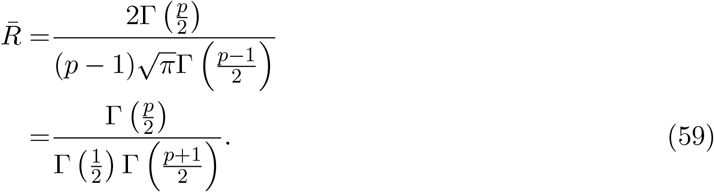

Taking 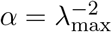, (15) becomes

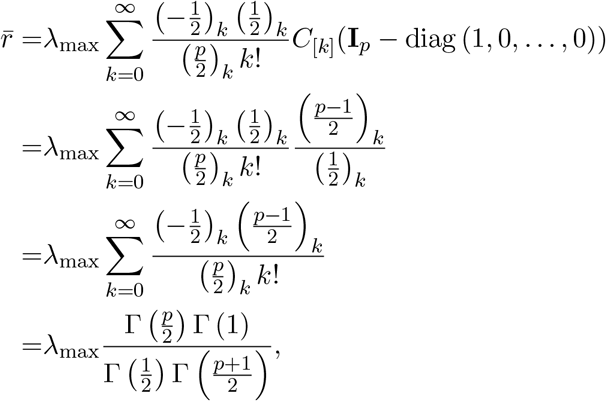

where the second equation comes from the fact *C*_[*k*]_(diag (0, 1, …, 1)) = *C*_[*k*]_(**I**_*p*−1_) and (28), and the last equation is from the Gauss hypergeometric theorem: 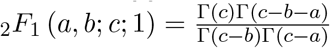. The equivalence to (59) is obvious.

## Footnotes

1 The term evolvability bears various meanings in the current evolutionary biology (e.g., Brown, 2014; Crother & Murray, 2019; Feiner *et al*., 2021; Jablonski, 2022; Love *et al*., 2022; Riederer *et al*., 2022). In the present paper, the use of this term is restricted to certain mathematical aspects of the **G** matrix and response vectors—the Hansen–Houle evolvability—unless stated otherwise.

2 A potential alternative to the average measures in the above sense is the average of the evolvability measures calculated from the **β**’s aligned with all individual traits (Armbruster *et al*., 2009; Haber, 2011; Pavlicev & Hansen, 2011; Machado *et al*., 2018). Although that would be valid for descriptive purposes, its interpretability in the context of multivariate selection might be limited. In any case, the calculation of such averages is straightforward and not concerned herein.

3 Rohlf (2017, p. 544) asserted that *ρ* cannot be negative because he assumed, inappropriately in this context, the response vectors to have no polarity. Negative *ρ* can occur when, e.g., **β** bisects the wider of the two angles formed by the major axes of highly integrated **G** matrices. Except for this subtle fallacy, his criticisms on previous interpretations of *ρ* or 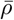 (e.g., Melo *et al*., 2016) seem largely pertinent (see also Discussion).

4 This confusion apparently arose from the custom of using the same symbol **β** for both standardized and unstandardized selection gradients (above). It seems advisable to use a different symbol for the former, e.g., **u** herein.

5 A few examples will clarify the indexing in (36) (semicolons separate terms of the summation). For (*k*_1_, *k*_2_) = (1, 1): (*i*_1_, *j*_2_) = (1, 1), *ν*_1,1_ = 1; (*ν*_1_, *ν*_2_) = (1, 0), (0, 1), *ν*_1,0_ = 1, *ν*_0,1_ = 1. For (*k*_1_, *k*_2_) = (2, 1): (*i*_1_, *i*_2_) = (2, 1), *ν*_2,1_ = 1; (*i*_1_, *i*_2_) = (2, 0), (0, 1), *ν*_2,0_ = 1, *ν*_0,1_ = 1; (*i*_1_, *i*_2_) = (1, 1), (1, 0), *ν*_1,1_ = 1, *ν*_1,0_ = 1; (*i*_1_, *i*_2_) = (1, 0), (0, 1), *ν*_1,0_ = 2, *ν*_0,1_ = 1.

6 The recursive algorithm for evaluating 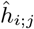 was said to be “very similar” to that for *h*_*i*;*j*_ but was not explicitly stated in Hillier *et al*. (2014). This is obtained by, in the definition and updating equation for *g*_*i,j*_ in the theorem 6 therein, replacing all the subtractions of terms (but not indices) by additions.

