## Supplementary material for "Exact expressions and numerical evaluation of average evolvability measures for characterizing and comparing G matrices": an_ae_fun.R

```
## These functions are maintained on the following repository:
## https://github.com/watanabe-j/avrevol
##
#' Series evaluation of average evolvability measures
#'
#' These functions evaluate series expressions of average evolvability measures
#' as their partial sums.
#'
#' All these functions are wrappers of either \code{qfratio::qfrm()} or
#' \code{qfratio::qfmrm()} for moments of simple and multiple ratios of
#' quadratic forms in normal variables.  See documentations of those functions
#' for technical details.
#'
#' When \eqn{\bm{G}} is singular, average conditional evolvability
#' and average autonomy can be nonzero only when the distribution of
#' \eqn{\bm{\beta}} is within the range of \eqn{\bm{G}}.
#' In \code{avr_cevo()} and \code{avr_auto()},
#' this condition is assessed by the QR decomposition of \eqn{\bm{G}}
#' (with pivoting).  When \eqn{\bm{G}} is not full-rank, \eqn{\bm{\mu}} and
#' \eqn{\bm{\Sigma}} are projected onto the range of \eqn{\bm{G}} using
#' the Q part of the QR decomposition.  If these are equal to the original ones,
#' then the arguments are passed to evaluation of the moment using quadratic
#' forms; otherwise, these functions return 0 (formatted as an \code{qfrm}
#' object).
#'
#' @param G,G1,G2
#'   Target covariance matrix/matrices.  Assumed validly structured
#'   (symmetric, nonnegative definite), although symmetry is checked.
#' @param m
#'   Order of evaluation \eqn{m} at which partial sum is truncated
#'   (\eqn{M} in, e.g., Hillier et al., 2014)
#'   Adjust this value depending on desired accuracy.
#' @param mu,Sigma
#'   Mean vector and covariance matrix, respectively, of selection gradients;
#'   default zero vector and identity matrix.
#'   Passed to \code{qfratio::qfrm()} or \code{qfratio::qfmrm()}.
#' @param cpp_method
#'   Option to specify \code{C++} algorithm to avoid numerical
#'   overflow/underflow in \code{qfratio::qfmrm()}.
#'   Default \code{"coef_wise"} is typically the most robust and
#'   modestly fast option. See documentation of
#'   \code{qfratio::\link[qfratio]{qfrm}()} for details.
#' @param tol_conv
#'   Tolerance against which numerical convergence is (roughly) checked.
#'   Default for \code{avr_auto()} and \code{avr_rcor()} is stricter than
#'   the default in \code{qfratio} because no error bound is available for these
#'   measures.
#' @param tol_qr
#'   Tolerance passed to \code{qr()} to determine singularity of \code{G}
#' @param ...
#'   Additional arguments are passed to \code{qfratio::qfrm()} or
#'   \code{qfratio::qfmrm()}.  Notable arguments involve \code{mu} and
#'   \code{Sigma} for the mean vector and covariance matrix, respectively,
#'   of selection gradients.
#'
#' @return
#' \code{qfrm} object defined by the package \code{qfratio}, which is a list
#' including \code{$statistic} (the partial sum) and \code{$error_bound}.
#' That package also defines \code{print} and \code{plot} methods.
#'

#### avr_evol ####
#' Series evaluation of average evolvability
avr_evol <- function(G, m = 100, ...) {
    stopifnot("G should be symmetric" = isSymmetric(G))
    qfratio::qfrm(G, p = 1, q = 1, m = m, ...)
}

#### avr_cevo ####
#' Series evaluation of average conditional evolvability
avr_cevo <- function(G, m = 100, mu = rep.int(0, dim(G)[1]),
                     Sigma = diag(dim(G)[1]), tol_qr = 1e-7, ...) {
    stopifnot("G should be symmetric" = isSymmetric(G))
    qrG <- qr(G, tol = tol_qr)
    if (qrG$rank < dim(G)[1]) {
        proj_RG <- tcrossprod(qr.Q(qrG)[, seq_len(qrG$rank)])
        mu_ <- proj_RG %*% c(mu)
        Sigma_ <- tcrossprod(proj_RG, tcrossprod(proj_RG, Sigma))
        beta_in_RG <- qfratio:::iseq(mu, mu_) && qfratio:::iseq(Sigma, Sigma_)
        if (beta_in_RG) {
            Gi <- MASS::ginv(G)
            return(qfratio::qfrm(B = Gi, p = 1, q = 1, m = m,
                                 mu = mu, Sigma = Sigma, ...))
        } else {
            message("Singular covariance matrix; ",
                    "average conditional evolvability is 0")
            return(qfratio:::new_qfrm(0, 0, 0, 0, exact = TRUE))
        }
    } else {
        Gi <- chol2inv(chol(G))
        return(qfratio::qfrm(B = Gi, p = 1, q = 1, m = m,
                             mu = mu, Sigma = Sigma, ...))
    }
}

#### avr_resp ####
#' Series evaluation of average respondability
avr_resp <- function(G, m = 100, ...) {
    stopifnot("G should be symmetric" = isSymmetric(G))
    Gsq <- crossprod(G)
    qfratio::qfrm(Gsq, p = 1/2, q = 1/2, m = m, ...)
}

#### avr_flex ####
#' Series evaluation of average flexibility
avr_flex <- function(G, m = 100, ...) {
    stopifnot("G should be symmetric" = isSymmetric(G))
    Gsq <- crossprod(G)
    qfratio::qfmrm(G, Gsq, p = 1, q = 1/2, r = 1/2, m = m, ...)
}

#### avr_auto ####
#' Series evaluation of average autonomy
avr_auto <- function(G, m = 100, mu = rep.int(0, dim(G)[1]),
                     Sigma = diag(dim(G)[1]),
                     cpp_method = c("coef_wise", "double", "long_double"),
                     tol_conv = sqrt(.Machine$double.eps),
                     tol_qr = 1e-7, ...) {
    cpp_method <- match.arg(cpp_method)
    stopifnot("G should be symmetric" = isSymmetric(G))
    qrG <- qr(G, tol = tol_qr)
    if (qrG$rank < dim(G)[1]) {
        proj_RG <- tcrossprod(qr.Q(qrG)[, seq_len(qrG$rank)])
        mu_ <- proj_RG %*% c(mu)
        Sigma_ <- tcrossprod(proj_RG, tcrossprod(proj_RG, Sigma))
        beta_in_RG <- qfratio:::iseq(mu, mu_) && qfratio:::iseq(Sigma, Sigma_)
        if (beta_in_RG) {
            Gi <- MASS::ginv(G)
            return(qfratio::qfmrm(B = G, D = Gi, p = 2, q = 1, r = 1, m = m,
                                  mu = mu, Sigma = Sigma,
                                  cpp_method = cpp_method, ...))
        } else {
            message("Singular covariance matrix; average autonomy is 0")
            return(qfratio:::new_qfrm(0, 0, 0, 0, exact = TRUE))
        }
    } else {
        Gi <- chol2inv(chol(G))
        return(qfratio::qfmrm(B = G, D = Gi, p = 2, q = 1, r = 1, m = m,
                              mu = mu, Sigma = Sigma,
                              cpp_method = cpp_method,
                              tol_conv = tol_conv, ...))
    }
}

#### avr_inte ####
#' Series evaluation of average integration
avr_inte <- function(G, ...) {
    auto <- avr_auto(G = G, ...)
    qfratio:::new_qfrm(statistic = 1 - auto$statistic,
                       terms = 1 - auto$terms,
                       error_bound = auto$error_bound,
                       seq_error = auto$seq_error)
}

#### avr_rdif ####
#' Series evaluation of average response difference
avr_rdif <- function(G1, G2, m = 100, ...) {
    stopifnot(
        "G1 and G2 should be symmetric" = isSymmetric(G1) && isSymmetric(G2),
        "G1 and G2 should have the same dimension" = all(dim(G1) == dim(G2))
    )
    Gdsq <- crossprod(G1 - G2)
    qfratio::qfrm(Gdsq, p = 1/2, q = 1/2, m = m, ...)
}

#### avr_rcor ####
#' Series evaluation of average response correlation
avr_rcor <- function(G1, G2, m = 100,
                     cpp_method = c("coef_wise", "double", "long_double"),
                     tol_conv = sqrt(.Machine$double.eps),
                     ...) {
    cpp_method <- match.arg(cpp_method)
    stopifnot(
        "G1 and G2 should be symmetric" = isSymmetric(G1) && isSymmetric(G2),
        "G1 and G2 should have the same dimension" = all(dim(G1) == dim(G2))
    )
    G12 <- crossprod(G1, G2)
    G1sq <- crossprod(G1)
    G2sq <- crossprod(G2)
    qfratio::qfmrm(G12, G1sq, G2sq, p = 1, q = 1/2, r = 1/2, m = m,
                   cpp_method = cpp_method,
                   tol_conv = tol_conv, ...)
}
```
