## Supplementary material for "Exact expressions and numerical evaluation of average evolvability measures for characterizing and comparing G matrices": hh_ae_fun.R

```
## These functions are maintained on the following repository:
## https://github.com/watanabe-j/avrevol
##
#' Delta method approximation of average evolvability measures
#'
#' These functions calculate Hansen & Houle's (2008, 2009)
#' delta method approximations for average evolvability measures.
#'
#' The notation \eqn{\mathrm{Var}} in Hansen & Houle's (2008, 2009) seems to
#' represent average sum of squares from the average (divided by k, the number
#' of variables) and not the ordinary unbiased estimator of variance for
#' infinite populations (divided by k - 1).
#' The package \code{evolvability} v 2.0.0 incorrectly uses the latter divisor.
#'
#' \code{svd(..., nu = 0L, nv = 0L)$d} is used to extract eigenvalues because
#' it is slightly faster than \code{eigen(..., symmetric = TRUE)$values}.
#' Note that the squared singular values of \code{A} is equal to the eigenvalues
#' of \code{crossprod(A)}.
#'
#' Evolvability measures not treated in Hansen & Houle (2008) (flexibility and
#' response correlation) have not been implemented (although this should be
#' doable). All the expressions are for the spherical normal (or uniform on
#' hypersphyere) distribution of selection gradients.
#'
#' @param G,G1,G2
#'   Target covariance matrix/matrices.  Assumed validly structured
#'   (symmetric, nonnegative definite), although symmetry is checked.
#'

#### hh_cevo ####
#' Average conditional evolvability using Hansen--Houle delta method
hh_cevo <- function(G) {
    stopifnot("G should be symmetric" = isSymmetric(G))
    I <- function(x) sum((x - mean(x)) ^ 2) / mean(x)^2 / length(x)
    L <- svd(G, nu = 0L, nv = 0L)$d
    p <- length(L)
    (1 + 2 * I(1 / L) / (p + 2)) / mean(1 / L)
}

#### hh_resp ####
#' Average respondability using Hansen--Houle delta method
hh_resp <- function(G) {
    stopifnot("G should be symmetric" = isSymmetric(G))
    I <- function(x) sum((x - mean(x)) ^ 2) / mean(x)^2 / length(x)
    L <- svd(G, nu = 0L, nv = 0L)$d
    p <- length(L)
    sqrt(mean(L ^ 2)) * (1 - I(L ^ 2) / 4 / (p + 2))
}

#### hh_rdif ####
#' Average response difference using Hansen--Houle delta method
hh_rdif <- function(G1, G2) {
    stopifnot(
        "G1 and G2 should be symmetric" = isSymmetric(G1) && isSymmetric(G2),
        "G1 and G2 should have the same dimension" = all(dim(G1) == dim(G2))
    )
    I <- function(x) sum((x - mean(x)) ^ 2) / mean(x)^2 / length(x)
    L12sq <- svd(G1 - G2, nu = 0L, nv = 0L)$d ^ 2
    p <- length(L12sq)
    sqrt(mean(L12sq)) * (1 - I(L12sq) / 4 / (p + 2))
}

#### hh_auto ####
#' Average autonomy using Hansen--Houle delta method
hh_auto <- function(G) {
    stopifnot("G should be symmetric" = isSymmetric(G))
    I <- function(x) sum((x - mean(x)) ^ 2) / mean(x)^2 / length(x)
    H <- function(x) 1 / mean(1 / x)
    L <- svd(G, nu = 0L, nv = 0L)$d
    p <- length(L)
    H(L) / mean(L) *
        ( 1 + 2 * (I(L) + I(1 / L) - 1 + H(L) / mean(L) +
         2 * I(L) * I(1 / L) / (p + 2)) / (p + 2) )
}
```
