## Supplementary material for "Exact expressions and numerical evaluation of average evolvability measures for characterizing and comparing G matrices": mc_ae_fun.R

```
## These functions are maintained on the following repository:
## https://github.com/watanabe-j/avrevol
##
#' Monte Carlo evaluation of average evolvability measures
#'
#' These functions execute Monte Carlo evaluation of
#' average evolvability measures
#'
#' All these functions are wrappers of either \code{qfratio::rqfr()} or
#' \code{qfratio::rqfmr()} for random number generation of simple and multiple
#' ratios of quadratic forms in normal variables.  See documentations of
#' those functions for technical details.
#'
#' @param nit
#'   Number of iteration
#' @param G,G1,G2
#'   Target covariance matrix/matrices.  Assumed validly structured
#'   (symmetric, nonnegative definite), although symmetry is checked.
#' @param mu,Sigma
#'   Mean vector and covariance matrix, respectively, of selection gradients;
#'   default zero vector and identity matrix.
#'   Passed to \code{qfratio::rqfr()} or \code{qfratio::rqfmr()}.
#' @param ...
#'   Additional arguments are passed to \code{qfratio::rqfr()} or
#'   \code{qfratio::rqfmr()}.  The only useful arguments will be \code{mu} and
#'   \code{Sigma} for the mean vector and covariance matrix, respectively,
#'   of selection gradients.
#'
#' @return
#' Numeric vector of length \code{b}
#'

#### mc_cevo ####
#' Monte Carlo sampling for (average) conditional evolvability
mc_cevo <- function(nit, G, mu = rep.int(0, dim(G)[1]),
                    Sigma = diag(dim(G)[1]), tol_qr = 1e-7, ...) {
    stopifnot("G should be symmetric" = isSymmetric(G))
    qrG <- qr(G, tol = tol_qr)
    if (qrG$rank < dim(G)[1]) {
        proj_RG <- tcrossprod(qr.Q(qrG)[, seq_len(qrG$rank)])
        mu_ <- proj_RG %*% c(mu)
        Sigma_ <- tcrossprod(proj_RG, tcrossprod(proj_RG, Sigma))
        beta_in_RG <- qfratio:::iseq(mu, mu_) && qfratio:::iseq(Sigma, Sigma_)
        if (beta_in_RG) {
            Gi <- MASS::ginv(G)
            return(qfratio::rqfr(nit, B = Gi, p = 1, q = 1,
                                 mu = mu, Sigma = Sigma, ...))
        } else {
            message("Singular covariance matrix; ",
                    "conditional evolvability is 0 with probability 1")
            return(rep.int(0, nit))
        }
    } else {
        Gi <- chol2inv(chol(G))
        return(qfratio::rqfr(nit, B = Gi, p = 1, q = 1, mu = mu, Sigma = Sigma,
                             ...))
    }
}

#### mc_resp ####
#' Monte Carlo sampling for (average) respondability
mc_resp <- function(nit, G, ...) {
    stopifnot("G should be symmetric" = isSymmetric(G))
    return(qfratio::rqfr(nit, crossprod(G), p = 1/2, q = 1/2, ...))
}

#### mc_flex ####
#' Monte Carlo sampling for (average) flexibility
mc_flex <- function(nit, G, ...) {
    stopifnot("G should be symmetric" = isSymmetric(G))
    return(qfratio::rqfmr(nit, G, crossprod(G), p = 1, q = 1/2, r = 1/2, ...))
}

#### mc_auto ####
#' Monte Carlo sampling for (average) autonomy
mc_auto <- function(nit, G, mu = rep.int(0, dim(G)[1]),
                    Sigma = diag(dim(G)[1]), tol_qr = 1e-7, ...) {
    stopifnot("G should be symmetric" = isSymmetric(G))
    qrG <- qr(G, tol = tol_qr)
    if (qrG$rank < dim(G)[1]) {
        proj_RG <- tcrossprod(qr.Q(qrG)[, seq_len(qrG$rank)])
        mu_ <- proj_RG %*% c(mu)
        Sigma_ <- tcrossprod(proj_RG, tcrossprod(proj_RG, Sigma))
        beta_in_RG <- qfratio:::iseq(mu, mu_) && qfratio:::iseq(Sigma, Sigma_)
        if (beta_in_RG) {
            Gi <- MASS::ginv(G)
            return(qfratio::rqfmr(nit, B = G, D = Gi, p = 2, q = 1, r = 1,
                                  mu = mu, Sigma = Sigma, ...))
        } else {
            message("Singular covariance matrix; ",
                    "autonomy is 0 with probability 1")
            return(rep.int(0, nit))
        }
    } else {
        Gi <- chol2inv(chol(G))
        return(qfratio::rqfmr(nit, B = G, D = Gi, p = 2, q = 1, r = 1,
                              mu = mu, Sigma = Sigma, ...))
    }
}

#### mc_rdif ####
#' Monte Carlo sampling for (average) response difference
mc_rdif <- function(nit, G1, G2, ...) {
    stopifnot(
        "G1 and G2 should be symmetric" = isSymmetric(G1) && isSymmetric(G2),
        "G1 and G2 should have the same dimension" = all(dim(G1) == dim(G2))
    )
    return(qfratio::rqfr(nit, crossprod(G1 - G2), p = 1/2, q = 1/2, ...))
}

#### mc_rcor ####
#' Monte Carlo sampling for (average) response correlation
mc_rcor <- function(nit, G1, G2, ...) {
    stopifnot(
        "G1 and G2 should be symmetric" = isSymmetric(G1) && isSymmetric(G2),
        "G1 and G2 should have the same dimension" = all(dim(G1) == dim(G2))
    )
    return(qfratio::rqfmr(nit, crossprod(G1, G2), crossprod(G1), crossprod(G2),
                          p = 1, q = 1/2, r = 1/2, ...))
}
```
