## Supplementary material for "Exact expressions and numerical evaluation of average evolvability measures for characterizing and comparing G matrices": plot_fun.R

```
CImean <- function(X, alpha = 0.95) {
    N <- length(X)
    Mean <- mean(X)
    SD <- sd(X)
    SE <- SD / sqrt(N)
    cp <- qt(c((1 - alpha) / 2, (1 + alpha) / 2), N - 1L)
    Mean + SE * cp
}

ae_plot <- function(what = c("cevo", "resp", "flex", "auto", "rdif", "rcor"),
                    p, suffix = c("1", "5", "q", "r", "15", "1q", "5q",
                                  "1r", "5r", "qr", "15r", "1qr", "5qr"),
                    main, xlab = bquote("Order of evaluation"* ~~italic(M)),
                    ylab, xlim, ylim, ylim_f,
                    col_am = "royalblue4", lwd_am = 1.5, lty_am = 1,
                    col_ae = "tomato", lwd_ae = 1, lty_ae = 2,
                    col_hh = "lightgreen", cex_hh = 1.5, pch_hh = 15,
                    col_M = "magenta", cex_M = 1.5, pch_M = 17, fc_M = 0.995,
                    col_mc = "orange3", lwd_mc = 2, lty_mc = 1,
                    cex_mc = 1.5, pch_mc = 20, add_legend = TRUE,
                    cex_time = 1, line_time1 = -3.5, line_time2 = -2,
                    cex_axis = 1, line_axis_x = NA, line_axis_y = NA,
                    cex_legend = 1, pos_leg = "topright",
                    use_short_mb = TRUE, xlim_short_m = FALSE,
                    show_M_nit = TRUE,
                    mark_M = if (what %in% c("cevo", "resp", "flex", "rdif") && !is.na(obja$error_bound)) obja$seq_error[m1] < 1e-8 else obja$terms[m1] < 1e-10,
                    ...) {
    require(qfratio)
    what <- match.arg(what)
    suffix <- match.arg(suffix)
    wh <- strtrim(what, 2L)
    obja <- get(paste0(wh, "a_p", sprintf("%03d", p), "_", suffix))
    objm <- get(paste0(wh, "m_p", sprintf("%03d", p), "_", suffix))
    mbres <- get(paste0("mb_", wh, "_p", sprintf("%03d", p), "_", suffix))
    m1 <- length(obja$terms)
    if (use_short_mb &&
       exists(paste0("m_", wh, "_p", sprintf("%03d", p), "_", suffix))) {
        m1 <- get(paste0("m_", wh, "_p", sprintf("%03d", p), "_", suffix)) + 1
    }
    b <- length(objm)
    if (missing(main)) {
        if (suffix == "1") main <- bquote(italic(p) == .(p)* ", " ~~ bold(G)[0.1])
        if (suffix == "5") main <- bquote(italic(p) == .(p)* ", " ~~ bold(G)[0.5])
        if (suffix == "q") main <- bquote(italic(p) == .(p)* ", " ~~ bold(G)[Q])
        if (suffix == "r") main <- bquote(italic(p) == .(p)* ", " ~~ bold(G)[Q])
        if (suffix == "15") main <- bquote(italic(p) == .(p)* ", " ~~ bold(G)[0.1]* " vs."* ~~ bold(G)[0.5]* ", same eigenvectors")
        if (suffix == "1q") main <- bquote(italic(p) == .(p)* ", " ~~ bold(G)[0.1]* " vs."* ~~ bold(G)[Q]* ", same eigenvectors")
        if (suffix == "5q") main <- bquote(italic(p) == .(p)* ", " ~~ bold(G)[0.5]* " vs."* ~~ bold(G)[Q]* ", same eigenvectors")
        if (suffix == "1r") main <- bquote(italic(p) == .(p)* ", two" ~~ bold(G)[0.1]* "'s, different eigenvectors")
        if (suffix == "5r") main <- bquote(italic(p) == .(p)* ", two" ~~ bold(G)[0.5]* "'s, different eigenvectors")
        if (suffix == "qr") main <- bquote(italic(p) == .(p)* ", two" ~~ bold(G)[Q]* "'s, different eigenvectors")
        if (suffix == "15r") main <- bquote(italic(p) == .(p)* ", " ~~ bold(G)[0.1]* " vs."* ~~ bold(G)[0.5]* ", different eigenvectors")
        if (suffix == "1qr") main <- bquote(italic(p) == .(p)* ", " ~~ bold(G)[0.1]* " vs."* ~~ bold(G)[Q]* ", different eigenvectors")
        if (suffix == "5qr") main <- bquote(italic(p) == .(p)* ", " ~~ bold(G)[0.5]* " vs."* ~~ bold(G)[Q]* ", different eigenvectors")
    }
    if (missing(ylab)) {
        ylab <- switch(what,
                       cevo = "Average conditional evolvability",
                       resp = "Average respondability",
                       flex = "Average flexibility",
                       auto = "Average autonomy",
                       rdif = "Average response difference",
                       rcor = "Average response correlation")
    }
    if (missing(xlim)) {
        xlimu <- if (xlim_short_m) m1 else length(obja$terms)
        xlim <- c(0, (xlimu - 1L) * 1.02)
    }
    if (missing(ylim)) {
        if (missing(ylim_f)) {
            ylim_f <- if (what %in% c("cevo", "flex", "auto", "rcor")) c(0.9, 1.05)
                      else c(0.95, 1.1)
        }
        ylim <- obja$statistic * ylim_f
    }
    plot(obja, add_legend = FALSE,
         xlim = xlim, ylim = ylim, xlab = xlab, ylab = ylab,
         col_m = col_am, lwd_m = lwd_am, lty_m = lty_am,
         col_e = col_ae, lwd_e = lwd_ae, lty_e = lty_ae,
         main = main, axes = FALSE, ...)
    box(bty = "l")
    axis(1, labels = FALSE)
    axis(1, tick = FALSE, cex.axis = cex_axis, line = line_axis_x)
    axis(2, labels = FALSE)
    axis(2, tick = FALSE, cex.axis = cex_axis, line = line_axis_y, las = 1)
    if (mark_M) {
        points(m1 - 1, sum(obja$terms[1:m1]) * fc_M, col = col_M, cex = cex_M, pch = pch_M)
    }
    if (what %in% c("cevo", "resp", "auto", "rdif")) {
        objh <- eval(parse(text = paste0(wh, "h_p",
                                         sprintf("%03d", p), "_", suffix)))
        points(xlim[2] * 0.98, objh, col = col_hh, cex = cex_hh, pch = pch_hh)
    }
    lines(cbind(xlim[2], CImean(objm)), col = col_mc, lwd = lwd_mc)
    points(xlim[2], mean(objm), col = col_mc, cex = cex_mc, pch = pch_mc)
    Times <- with(mbres, tapply(time, expr, mean))
    if (is.na(Times["an"])) {
        stres <- eval(parse(text = paste0("st_", wh, "a_p", sprintf("%03d", p), "_", suffix)))
        Times <- Times / 10^9
        if (show_M_nit) {
            mtext(bquote(italic(t)[S]~~ "("* italic(M) == .(sprintf("%i", m1 - 1L))* "): "* .(sprintf("%.2f", stres["elapsed"]))* " s*"), 1, line = line_time1, at = xlim[2], adj = 1, cex = cex_time)
            mtext(bquote(italic(t)[MC]~~ "("* italic(n)[it] == .(sprintf("%i", b))* "): "* .(sprintf("%.2f", Times["mc"]))* " s"), 1, line = line_time2, at = xlim[2], adj = 1, cex = cex_time)
        } else {
            mtext(bquote(italic(t)[S]* ": "* .(sprintf("%.2f", stres["elapsed"]))* " s*"), 1, line = line_time1, at = xlim[2], adj = 1, cex = cex_time)
            mtext(bquote(italic(t)[MC]* ": "* .(sprintf("%.2f", Times["mc"]))* " s"), 1, line = line_time2, at = xlim[2], adj = 1, cex = cex_time)
        }
    } else {
        if (max(Times) >= 10^9) {
            Times <- Times / 10^9
            if (show_M_nit) {
                mtext(bquote(italic(t)[S]~~ "("* italic(M) == .(sprintf("%i", m1 - 1L))* "): "* .(sprintf("%.2f", Times["an"]))* " s"), 1, line = line_time1, at = xlim[2], adj = 1, cex = cex_time)
                mtext(bquote(italic(t)[MC]~~ "("* italic(n)[it] == .(sprintf("%i", b))* "): "* .(sprintf("%.2f", Times["mc"]))* " s"), 1, line = line_time2, at = xlim[2], adj = 1, cex = cex_time)
            } else {
                mtext(bquote(italic(t)[S]* ": "* .(sprintf("%.2f", Times["an"]))* " s"), 1, line = line_time1, at = xlim[2], adj = 1, cex = cex_time)
                mtext(bquote(italic(t)[MC]* ": "* .(sprintf("%.2f", Times["mc"]))* " s"), 1, line = line_time2, at = xlim[2], adj = 1, cex = cex_time)
            }
        } else {
            Times <- Times / 10^6
            if (show_M_nit) {
                mtext(bquote(italic(t)[S]~~ "("* italic(M) == .(sprintf("%i", m1 - 1L))* "): "* .(sprintf("%.1f", Times["an"]))* " ms"), 1, line = line_time1, at = xlim[2], adj = 1, cex = cex_time)
                mtext(bquote(italic(t)[MC]~~ "("* italic(n)[it] == .(sprintf("%i", b))* "): "* .(sprintf("%.1f", Times["mc"]))* " ms"), 1, line = line_time2, at = xlim[2], adj = 1, cex = cex_time)
            } else {
                mtext(bquote(italic(t)[S]* ": "* .(sprintf("%.1f", Times["an"]))* " ms"), 1, line = line_time1, at = xlim[2], adj = 1, cex = cex_time)
                mtext(bquote(italic(t)[MC]* ": "* .(sprintf("%.1f", Times["mc"]))* " ms"), 1, line = line_time2, at = xlim[2], adj = 1, cex = cex_time)
            }
        }
    }
    if (add_legend) {
        legend(pos_leg, cex = cex_legend,
            legend = c("Series", if (what %in% c("cevo", "resp", "flex", "rdif")) "Error bound" else NULL, "Convergence", "MC mean & 95% CI", if (what %in% c("cevo", "resp", "auto", "rdif")) "Delta method" else NULL),
            col    = c(col_am,   if (what %in% c("cevo", "resp", "flex", "rdif")) col_ae        else NULL, col_M,          col_mc,             if (what %in% c("cevo", "resp", "auto", "rdif")) col_hh         else NULL),
            lwd    = c(lwd_am,   if (what %in% c("cevo", "resp", "flex", "rdif")) lwd_ae        else NULL, NA,             NA,                 if (what %in% c("cevo", "resp", "auto", "rdif")) NA             else NULL),
            lty    = c(lty_am,   if (what %in% c("cevo", "resp", "flex", "rdif")) lty_ae        else NULL, NA,             NA,                 if (what %in% c("cevo", "resp", "auto", "rdif")) NA             else NULL),
            pch    = c(NA,       if (what %in% c("cevo", "resp", "flex", "rdif")) NA            else NULL, pch_M,          pch_mc,             if (what %in% c("cevo", "resp", "auto", "rdif")) pch_hh         else NULL),
            pt.cex = c(NA,       if (what %in% c("cevo", "resp", "flex", "rdif")) NA            else NULL, cex_M,          cex_mc,             if (what %in% c("cevo", "resp", "auto", "rdif")) cex_hh         else NULL),
        )
    }
}

ae_plot_mat <- function(what = c("cevo", "resp", "flex", "auto", "rdif", "rcor"),
    ps = c(5, 20, 50, 200),
    suffixes = if (what %in% c("cevo", "resp" , "flex", "auto")) c("1", "5") else c("1q", "qr"),
    device = c("x11", "pdf", "png"), file = paste0("mat_", what, ".", device),
    width = 80 * length(ps) / 25.4 * if (device == "png") dpi else 1,
    height = 80 * length(suffixes) / 25.4 * if (device == "png") dpi else 1,
    dpi = 300, bg_png = "transparent",
    new_dev = dev.cur() == 1, do_dev.off = new_dev, colormodel = "rgb",
    cex_time = 0.8, ylabs = NULL, cex_xlab = 0.8, cex_ylab = cex_xlab,
    line_ylab = 3.5, line_plab = 0, line_xlab = 3,
    show_plab = TRUE, show_xlab = TRUE, show_ylab = TRUE, add_legend = TRUE,
    parlist = list(mfrow = c(length(suffixes), length(ps)), mar = c(3, 3, 0, 0), oma = c(1.5, 2, 2, 1)),
    ...) {
    what <- match.arg(what)
    device <- match.arg(device)
    symb_bq <- switch(what, cevo = bquote(bar(italic(c))),
                            resp = bquote(bar(italic(r))),
                            flex = bquote(bar(italic(f))),
                            auto = bquote(bar(italic(a))),
                            rdif = bquote(bar(italic(d))),
                            rcor = bquote(bar(italic(rho))))
    if (missing(ylabs)) {
        names(suffixes) <- suffixes
        ylabs <- lapply(suffixes, function(s) {
            switch(s, "1" = bquote(.(symb_bq): ~~ bold(G)[0.1]),
                      "5" = bquote(.(symb_bq): ~~ bold(G)[0.5]),
                      "q" = bquote(.(symb_bq): ~~ bold(G)[Q]),
                      "15" = bquote(.(symb_bq): bold(G)[0.1]* " vs. "* bold(G)[0.5]* ", same evecs"),
                      "1q" = bquote(.(symb_bq): bold(G)[0.1]* " vs. "* bold(G)[Q]* ", same evecs"),
                      "5q" = bquote(.(symb_bq): bold(G)[0.5]* " vs. "* bold(G)[Q]* ", same evecs"),
                      "1r" = bquote(.(symb_bq): "two" ~~ bold(G)[0.1]* ", different evecs"),
                      "5r" = bquote(.(symb_bq): "two" ~~ bold(G)[0.5]* ", different evecs"),
                      "qr" = bquote(.(symb_bq): "two" ~~ bold(G)[Q]* ", different evecs"),
                      "15r" = bquote(.(symb_bq): ~~ bold(G)[0.1]* " vs. "* bold(G)[0.5]* ", different evecs"),
                      "1qr" = bquote(.(symb_bq): ~~ bold(G)[0.1]* " vs. "* bold(G)[Q]* ", different evecs"),
                      "5qr" = bquote(.(symb_bq): ~~ bold(G)[0.5]* " vs. "* bold(G)[Q]* ", different evecs"))
        })
    }
    if (new_dev) {
        if (device == "pdf") {
            pdf(file, width = width, height = height, colormodel = colormodel)
            if (do_dev.off) on.exit(dev.off())
        } else if (device == "png") {
            png(file, width = width, height = height, res = dpi, bg = bg_png)
            if (do_dev.off) on.exit(dev.off())
        } else {
            x11(width = width, height = height)
        }
    }
    par(parlist)
    for (s in suffixes) {
        for (p in ps) {
            ae_plot(what, p, s, main = "", xlab = "", ylab = "",
                cex_time = cex_time,
                add_legend = (p == ps[length(ps)]) && (s == suffixes[1]) && add_legend, ...)
            if (p == ps[1] && show_ylab) {
                mtext(ylabs[[s]], 2, line_ylab, cex = cex_ylab)
            }
            if (s == suffixes[1] && show_plab) {
                mtext(bquote(italic(p)==.(p)), 3, line_plab, cex = cex_xlab)
            }
            if (s == suffixes[length(suffixes)] && show_xlab) {
                mtext(bquote("Order of evaluation"* ~~italic(M)), 1, line_xlab,
                      cex = cex_xlab)
            }
        }
    }
}
```
