## Supplementary material for "Exact expressions and numerical evaluation of average evolvability measures for characterizing and comparing G matrices": README.txt

###############
## Supplementary Material for
##   Exact expressions and numerical evaluation of average evolvability measures
##   for characterizing and comparing G matrices
## Junya Watanabe - University of Cambridge, UK.
##
## This package contains supplementary R script for reproducing
## numerical results of the above paper.
##
## Contents:
## - suppl_script.R: Main script file
## - R/
##     an_ae_fun.R:  Functions for analytic/series evaluation
##     hh_ae_fun.R:  Functions for Hansen-Houle delta method
##     mc_ae_fun.R:  Functions for Monte Carlo evaluation
##     plot_fun.R:   Functions for graphical outputs
##
## The main functions in this supplement (an_ae_fun.R, hh_ae_fun.R, mc_ae_fun.R)
## are maintained in the following repository:
## https://github.com/watanabe-j/avrevol
##
## Usage:
## From command line, move to this root directory and run
## $  Rscript suppl_script.R
## Alternatively, use R GUI and
## R> setwd("path/to/this/directory")
## R> source("suppl_script.R")  # Or copy-paste its contents if preferred
## Note that warnings for numerical non-convergence may happen
##
## WARNING:
## Running the script typically takes a long time (e.g., ~8 h)
## and writes a large image file (~1 GB) as an output.
## It then draws several graphic outputs in two PDFs.
## A large RAM (>8 GB) will be required.
##
## The scripts were tested with the following dependencies:
## R              v. 4.2.3 (incl. stats, graphics, grDevices)
## microbenchmark v. 1.4.9
## eigvaldisp     v. 0.0.0.9404
## qfratio        v. 1.0.0
## Rcpp           v. 1.0.10
## RcppEigen      v. 0.3.3.9.3
###############
