## Supplementary material for "Exact expressions and numerical evaluation of average evolvability measures for characterizing and comparing G matrices": suppl_script.R

```
###############
## Supplementary Material for
##   Exact expressions and numerical evaluation of average evolvability measures
##   for characterizing and comparing G matrices
## Junya Watanabe - University of Cambridge, UK.
##
## This R script is for reproducing numerical experiments in the paper.
## Running this script takes a long time and writes large files
## (see README.txt).
##
## The resultant environment will contain a number of objects like
## cea_p005_1 or rdm_p200_15r.  The names of these represent:
## - Evolvability measure (first two characters):
##    conditional evolvability (ce), respondability (re),
##    flexibility (fl), autonomy (au), response difference (rd), and
##    response correlation (rc)
## - Evaluation method (third character):
##    Analytic/series (a), Monte Carlo (m), and Hansen-Houle delta method (h)
## - Number of traits (_p***)
## - Shape of covariance matrix (suffix after _):
##    For one-matrix measures, Vrel = 0.1, 0.5 (1/5) or
##    quadratically decreasing eigenvalues (q).
##    For two-matrix mreasures, this is a combination of two from 1, 5, and q,
##    with "r" (rotated) at the end represents combination with different
##    eigenvectors.
##
## Objects like mb_ce_p005_1 are microbenchmark objects that contain
## execution times for analytic (an) and Monte Carlo (mc) evaluations.
## For response correlation with p = 50, most of these are replaced by
## objects like st_rca_p050_15 which are proc_time objects (from system.time()).
###############

# install.packages(c("microbenchmark", "devtools", "qfratio"))
# devtools::install_github("watanabe-j/eigvaldisp")
library(microbenchmark)
library(eigvaldisp)
library(qfratio)
source("R/an_ae_fun.R")
source("R/mc_ae_fun.R")
source("R/hh_ae_fun.R")

sink("output.Rout", split = TRUE)


n_it_mc <- 10000L
n_benchmark <- 10L

## Tolerance to be used to determine singularity
## (larger than default for large problems)
tol <- 1e-10

ps <- c(5L, 20L, 50L, 200L)
ms_cevo <- c( 300,  1800,   7800, 260000)
ms_resp <- c(1600, 24000, 200000,  9.2e6)
ms_flex <- c(3000, 33000, 240000,  9.9e6)
ms_auto <- c( 330,  1900,   5000,  17000)
ms_rdif <- c(7000,  9900,  93000,  4.9e6)
ms_rcor <- c(4000,  4000,  10000,    500)

s3 <- function(p) sprintf("%03d", p)

ceiling2d <- function(x) {
    digits <- 10 ^ (floor(log10(x)) - 1)
    as.integer(ceiling(x / digits) * digits)
}

get_avrevol_obj <- function(wh, p, suffix, prefix = NULL) {
    get( paste0(prefix, wh, "a_p", s3(p), "_", suffix) )
}

get_m_from_errorbound <- function(wh, p, suffix, thr_e = 1e-8) {
    obj <- get_avrevol_obj(wh, p, suffix)
    m <- with(obj, ceiling2d(which(abs(seq_error) < thr_e)[1] - 1))
    if (is.na(m)) m <- length(obj$terms) - 1
    return(m)
}

get_m_from_asymptote <- function(wh, p, suffix, sdig = 9) {
    obj <- get_avrevol_obj(wh, p, suffix)
    m <- with(obj, ceiling2d(which(round(cumsum(terms), sdig) ==
                                   round(statistic, sdig))[1] - 1))
    if (is.na(m)) m <- length(obj$terms) - 1
    return(m)
}

###########
## Generate covariance matrices and
## execute series, Monte Carlo, and delta method evaluations
for (i in seq_along(ps)) {
    p      <- ps     [i]
    m_cevo <- ms_cevo[i]
    m_resp <- ms_resp[i]
    m_flex <- ms_flex[i]
    m_auto <- ms_auto[i]
    m_rdif <- ms_rdif[i]
    m_rcor <- ms_rcor[i]

    G1 <- GenCov(p = p, VR = 0.1, shape = "elongate")
    G5 <- GenCov(p = p, VR = 0.5, shape = "elongate")
    Gq <- GenCov(p = p, shape = "quadratically_decreasing")
    L1 <- diag(G1)
    L5 <- diag(G5)
    Lq <- diag(Gq)
    G1r <- GenCov(evalues = L1, evectors = "Givens", random_rotation = FALSE,
                  which_rotate = "head", scale. = sum(L1))
    G5r <- GenCov(evalues = L5, evectors = "Givens", random_rotation = FALSE,
                  which_rotate = "head", scale. = sum(L5))
    Gqr <- GenCov(evalues = Lq, evectors = "Givens", random_rotation = FALSE,
                  which_rotate = "head", scale. = sum(Lq))

    assign(paste0("G1_p", s3(p)), G1)
    assign(paste0("G5_p", s3(p)), G5)
    assign(paste0("Gq_p", s3(p)), Gq)
    assign(paste0("G1r_p", s3(p)), G1r)
    assign(paste0("G5r_p", s3(p)), G5r)
    assign(paste0("Gqr_p", s3(p)), Gqr)

    assign(paste0("cea_p", s3(p), "_1"), avr_cevo(G1, m_cevo, tol_sing = tol))
    assign(paste0("rea_p", s3(p), "_1"), avr_resp(G1, m_resp, tol_sing = tol))
    assign(paste0("fla_p", s3(p), "_1"), avr_flex(G1, m_flex, tol_sing = tol))
    assign(paste0("aua_p", s3(p), "_1"), avr_auto(G1, m_auto, tol_sing = tol,
                                             nthreads = if (p == 200) 0 else 1))
    assign(paste0("cea_p", s3(p), "_5"), avr_cevo(G5, m_cevo, tol_sing = tol))
    assign(paste0("rea_p", s3(p), "_5"), avr_resp(G5, m_resp, tol_sing = tol))
    assign(paste0("fla_p", s3(p), "_5"), avr_flex(G5, m_flex, tol_sing = tol))
    assign(paste0("aua_p", s3(p), "_5"), avr_auto(G5, m_auto, tol_sing = tol,
                                             nthreads = if (p == 200) 0 else 1))
    assign(paste0("cea_p", s3(p), "_q"), avr_cevo(Gq, m_cevo, tol_sing = tol))
    assign(paste0("rea_p", s3(p), "_q"), avr_resp(Gq, m_resp, tol_sing = tol))
    assign(paste0("fla_p", s3(p), "_q"), avr_flex(Gq, m_flex, tol_sing = tol))
    assign(paste0("aua_p", s3(p), "_q"), avr_auto(Gq, m_auto, tol_sing = tol,
                                             nthreads = if (p == 200) 0 else 1))

    assign(paste0("rda_p", s3(p), "_15"),  avr_rdif(G1, G5,  m_rdif,
                                                    tol_sing = tol))
    assign(paste0("rda_p", s3(p), "_1q"),  avr_rdif(G1, Gq,  m_rdif,
                                                    tol_sing = tol))
    assign(paste0("rda_p", s3(p), "_5q"),  avr_rdif(G5, Gq,  m_rdif,
                                                    tol_sing = tol))
    assign(paste0("rda_p", s3(p), "_1r"),  avr_rdif(G1, G1r, m_rdif,
                                                    tol_sing = tol))
    assign(paste0("rda_p", s3(p), "_5r"),  avr_rdif(G5, G5r, m_rdif,
                                                    tol_sing = tol))
    assign(paste0("rda_p", s3(p), "_qr"),  avr_rdif(Gq, Gqr, m_rdif,
                                                    tol_sing = tol))
    assign(paste0("rda_p", s3(p), "_15r"), avr_rdif(G1, G5r, m_rdif,
                                                    tol_sing = tol))
    assign(paste0("rda_p", s3(p), "_1qr"), avr_rdif(G1, Gqr, m_rdif,
                                                    tol_sing = tol))
    assign(paste0("rda_p", s3(p), "_5qr"), avr_rdif(G5, Gqr, m_rdif,
                                                    tol_sing = tol))

    if (p <= 50) {
        assign(paste0("rca_p", s3(p), "_15"),
               avr_rcor(G1, G5,  m_rcor, tol_sing = tol))
        assign(paste0("rca_p", s3(p), "_1q"),
               avr_rcor(G1, Gq,  m_rcor, tol_sing = tol))
        assign(paste0("rca_p", s3(p), "_5q"),
               avr_rcor(G5, Gq,  m_rcor, tol_sing = tol))
        assign(paste0("rca_p", s3(p), "_1r"),
               avr_rcor(G1, G1r, m_rcor, tol_sing = tol))
        assign(paste0("rca_p", s3(p), "_5r"),
               avr_rcor(G5, G5r, m_rcor, tol_sing = tol))
        assign(paste0("rca_p", s3(p), "_qr"),
               avr_rcor(Gq, Gqr, m_rcor, tol_sing = tol))
        assign(paste0("rca_p", s3(p), "_15r"),
               avr_rcor(G1, G5r, m_rcor, tol_sing = tol))
        assign(paste0("rca_p", s3(p), "_1qr"),
               avr_rcor(G1, Gqr, m_rcor, tol_sing = tol))
        assign(paste0("rca_p", s3(p), "_5qr"),
               avr_rcor(G5, Gqr, m_rcor, tol_sing = tol))
    }

    assign(paste0("cem_p", s3(p), "_1"), mc_cevo(n_it_mc, G1))
    assign(paste0("rem_p", s3(p), "_1"), mc_resp(n_it_mc, G1))
    assign(paste0("flm_p", s3(p), "_1"), mc_flex(n_it_mc, G1))
    assign(paste0("aum_p", s3(p), "_1"), mc_auto(n_it_mc, G1))
    assign(paste0("cem_p", s3(p), "_5"), mc_cevo(n_it_mc, G5))
    assign(paste0("rem_p", s3(p), "_5"), mc_resp(n_it_mc, G5))
    assign(paste0("flm_p", s3(p), "_5"), mc_flex(n_it_mc, G5))
    assign(paste0("aum_p", s3(p), "_5"), mc_auto(n_it_mc, G5))
    assign(paste0("cem_p", s3(p), "_q"), mc_cevo(n_it_mc, Gq))
    assign(paste0("rem_p", s3(p), "_q"), mc_resp(n_it_mc, Gq))
    assign(paste0("flm_p", s3(p), "_q"), mc_flex(n_it_mc, Gq))
    assign(paste0("aum_p", s3(p), "_q"), mc_auto(n_it_mc, Gq))

    assign(paste0("rdm_p", s3(p), "_15"), mc_rdif(n_it_mc, G1, G5))
    assign(paste0("rdm_p", s3(p), "_1q"), mc_rdif(n_it_mc, G1, Gq))
    assign(paste0("rdm_p", s3(p), "_5q"), mc_rdif(n_it_mc, G5, Gq))
    assign(paste0("rdm_p", s3(p), "_1r"), mc_rdif(n_it_mc, G1, G1r))
    assign(paste0("rdm_p", s3(p), "_5r"), mc_rdif(n_it_mc, G5, G5r))
    assign(paste0("rdm_p", s3(p), "_qr"), mc_rdif(n_it_mc, Gq, Gqr))
    assign(paste0("rdm_p", s3(p), "_15r"), mc_rdif(n_it_mc, G1, G5r))
    assign(paste0("rdm_p", s3(p), "_1qr"), mc_rdif(n_it_mc, G1, Gqr))
    assign(paste0("rdm_p", s3(p), "_5qr"), mc_rdif(n_it_mc, G5, Gqr))
    assign(paste0("rcm_p", s3(p), "_15"), mc_rcor(n_it_mc, G1, G5))
    assign(paste0("rcm_p", s3(p), "_1q"), mc_rcor(n_it_mc, G1, Gq))
    assign(paste0("rcm_p", s3(p), "_5q"), mc_rcor(n_it_mc, G5, Gq))
    assign(paste0("rcm_p", s3(p), "_1r"), mc_rcor(n_it_mc, G1, G1r))
    assign(paste0("rcm_p", s3(p), "_5r"), mc_rcor(n_it_mc, G5, G5r))
    assign(paste0("rcm_p", s3(p), "_qr"), mc_rcor(n_it_mc, Gq, Gqr))
    assign(paste0("rcm_p", s3(p), "_15r"), mc_rcor(n_it_mc, G1, G5r))
    assign(paste0("rcm_p", s3(p), "_1qr"), mc_rcor(n_it_mc, G1, Gqr))
    assign(paste0("rcm_p", s3(p), "_5qr"), mc_rcor(n_it_mc, G5, Gqr))

    assign(paste0("ceh_p", s3(p), "_1"), hh_cevo(G1))
    assign(paste0("reh_p", s3(p), "_1"), hh_resp(G1))
    assign(paste0("auh_p", s3(p), "_1"), hh_auto(G1))
    assign(paste0("ceh_p", s3(p), "_5"), hh_cevo(G5))
    assign(paste0("reh_p", s3(p), "_5"), hh_resp(G5))
    assign(paste0("auh_p", s3(p), "_5"), hh_auto(G5))
    assign(paste0("ceh_p", s3(p), "_q"), hh_cevo(Gq))
    assign(paste0("reh_p", s3(p), "_q"), hh_resp(Gq))
    assign(paste0("auh_p", s3(p), "_q"), hh_auto(Gq))

    assign(paste0("rdh_p", s3(p), "_15"), hh_rdif(G1, G5))
    assign(paste0("rdh_p", s3(p), "_1q"), hh_rdif(G1, Gq))
    assign(paste0("rdh_p", s3(p), "_5q"), hh_rdif(G5, Gq))
    assign(paste0("rdh_p", s3(p), "_1r"), hh_rdif(G1, G1r))
    assign(paste0("rdh_p", s3(p), "_5r"), hh_rdif(G5, G5r))
    assign(paste0("rdh_p", s3(p), "_qr"), hh_rdif(Gq, Gqr))
    assign(paste0("rdh_p", s3(p), "_15r"), hh_rdif(G1, G5r))
    assign(paste0("rdh_p", s3(p), "_1qr"), hh_rdif(G1, Gqr))
    assign(paste0("rdh_p", s3(p), "_5qr"), hh_rdif(G5, Gqr))

    save.image("result.Rdata")
    cat("  p =", p, "done\n")
}

save.image("result.Rdata")


###########
## Choose sufficient order of evaluation from previous results
## and measure execution time with that (all except rho)
for (p in ps) {
    for (suffix in c("1", "5", "q")) {
        G <- get(paste0("G", suffix, "_p", s3(p)))
        for (wh in c("ce", "re", "fl")) {
            anfun <- switch(wh, ce = avr_cevo, re = avr_resp, fl = avr_flex)
            mcfun <- switch(wh, ce = mc_cevo, re = mc_resp, fl = mc_flex)
            m <- get_m_from_errorbound(wh, p, suffix)
            assign(paste0("m_", wh, "_p", s3(p), "_", suffix), m)
            assign(paste0("mb_", wh, "_p", s3(p), "_", suffix),
                   microbenchmark(an = anfun(G, m, tol_sing = tol),
                                  mc = mcfun(n_it_mc, G),
                                  times = n_benchmark))
        }
        m <- get_m_from_asymptote("au", p, suffix)
        assign(paste0("m_au_p", s3(p), "_", suffix), m)
        assign(paste0("mb_au_p", s3(p), "_", suffix),
               microbenchmark(an = avr_auto(G, m, tol_sing = tol),
                              mc = mc_auto(n_it_mc, G),
                              times = n_benchmark))
    }
    for (suffix in c("15", "1q", "5q", "15r", "1qr", "5qr")) {
        G1 <- get(paste0("G", substr(suffix, 1, 1), "_p", s3(p)))
        G2 <- get(paste0("G", substr(suffix, 2, 3), "_p", s3(p)))
        m <- get_m_from_errorbound("rd", p, suffix)
        assign(paste0("m_rd_p", s3(p), "_", suffix), m)
        assign(paste0("mb_rd_p", s3(p), "_", suffix),
               microbenchmark(an = avr_rdif(G1, G2, m, tol_sing = tol),
                              mc = mc_rdif(n_it_mc, G1, G2),
                              times = n_benchmark))
    }
    for (suffix in c("1r", "5r", "qr")) {
        G1 <- get(paste0("G", substr(suffix, 1, 1), "_p", s3(p)))
        G2 <- get(paste0("G", substr(suffix, 1, 2), "_p", s3(p)))
        m <- get_m_from_errorbound("rd", p, suffix)
        assign(paste0("m_rd_p", s3(p), "_", suffix), m)
        assign(paste0("mb_rd_p", s3(p), "_", suffix),
               microbenchmark(an = avr_rdif(G1, G2, m, tol_sing = tol),
                              mc = mc_rdif(n_it_mc, G1, G2),
                              times = n_benchmark))
    }
    cat("  Measures except rho for p =", p, "done..\n")
}

save.image("result.Rdata")

###########
## rho; don't do benchmarking of series evaluation for p = 200 or
## p = 50 and G5 is involved
for (p in ps) {
    for (suffix in c("15", "1q", "5q", "15r", "1qr", "5qr")) {
        G1 <- get(paste0("G", substr(suffix, 1, 1), "_p", s3(p)))
        G2 <- get(paste0("G", substr(suffix, 2, 3), "_p", s3(p)))
        if (p > 50 || (p == 50 && grepl("5", suffix))) {
            assign(paste0("m_rc_p", s3(p), "_", suffix), ms_rcor[ps == p])#dummy
            assign(paste0("mb_rc_p", s3(p), "_", suffix),
                microbenchmark(mc = mc_rcor(n_it_mc, G1, G2),
                               times = n_benchmark))
        } else {
            m <- get_m_from_asymptote("rc", p, suffix)
            assign(paste0("m_rc_p", s3(p), "_", suffix), m)
            assign(paste0("mb_rc_p", s3(p), "_", suffix),
                microbenchmark(an = avr_rcor(G1, G2, m, tol_sing = tol),
                               mc = mc_rcor(n_it_mc, G1, G2),
                               times = n_benchmark))
        }
    }
    for (suffix in c("1r", "5r", "qr")) {
        G1 <- get(paste0("G", substr(suffix, 1, 1), "_p", s3(p)))
        G2 <- get(paste0("G", substr(suffix, 1, 2), "_p", s3(p)))
        if (p > 50 || (p == 50 && grepl("5", suffix))) {
            assign(paste0("m_rc_p", s3(p), "_", suffix), ms_rcor[ps == p])#dummy
            assign(paste0("mb_rc_p", s3(p), "_", suffix),
                microbenchmark(mc = mc_rcor(n_it_mc, G1, G2),
                               times = n_benchmark))
        } else {
            m <- get_m_from_asymptote("rc", p, suffix)
            assign(paste0("m_rc_p", s3(p), "_", suffix), m)
            assign(paste0("mb_rc_p", s3(p), "_", suffix),
                microbenchmark(an = avr_rcor(G1, G2, m, tol_sing = tol),
                                mc = mc_rcor(n_it_mc, G1, G2),
                                times = n_benchmark))
        }
    }
    cat("  rho for p =", p, "done..\n")
    save.image("result.Rdata")
}

###########
## rho, p = 50
## For any combinations with G5, convergence is not attained,
## so benchmarking is moot; just do system.time
st_rca_p050_15  <- system.time(rca_p050_15 <-
                                   avr_rcor(G1_p050, G5_p050, ms_rcor[3]))
st_rca_p050_5q  <- system.time(rca_p050_5q <-
                                   avr_rcor(G5_p050, Gq_p050, ms_rcor[3]))
save.image("result.Rdata"); cat("  rho for p =", 50, "; 15, 5q done..\n")
st_rca_p050_5r  <- system.time(rca_p050_5r <-
                                   avr_rcor(G5_p050, G5r_p050, ms_rcor[3]))
save.image("result.Rdata"); cat("  rho for p =", 50, "; 5r done..\n")
st_rca_p050_15r <- system.time(rca_p050_15r <-
                                   avr_rcor(G1_p050, G5r_p050, ms_rcor[3]))
save.image("result.Rdata"); cat("  rho for p =", 50, "; 15r done..\n")
st_rca_p050_5qr <- system.time(rca_p050_5qr <-
                                   avr_rcor(G5_p050, Gqr_p050, ms_rcor[3]))
save.image("result.Rdata"); cat("  rho for p =", 50, "; 5qr done..\n")

###########
## rho, p = 200
## The only condition that reaches convergence is Gq vs Gqr
rca_p200_qr  <- avr_rcor(Gq_p200, Gqr_p200, ms_rcor[4])
m_rc_p200_qr <- get_m_from_asymptote("rc", 200, "qr")
mb_rc_p200_qr <- microbenchmark(
    an = avr_rcor(Gq_p200, Gqr_p200, m_rc_p200_qr, tol_sing = tol),
    mc = mc_rcor(n_it_mc, Gq_p200, Gqr_p200), times = n_benchmark)

save.image("result.Rdata")

###########
## These are used to supplement main calculations for rho with G5 and p >= 20
## Require ~250 GB RAM, so commented out here
# st_rca_p020_15_c  <- system.time(rca_p020_15_c  <- avr_rcor(G1_p020, G5_p020,  60000, nthreads = 0))
# save.image("result.Rdata")
# st_rca_p020_5q_c  <- system.time(rca_p020_5q_c  <- avr_rcor(G5_p020, Gq_p020,  60000, nthreads = 0))
# save.image("result.Rdata")
# st_rca_p020_5r_c  <- system.time(rca_p020_5r_c  <- avr_rcor(G5_p020, G5r_p020, 60000, nthreads = 0))
# save.image("result.Rdata")
# st_rca_p020_15r_c <- system.time(rca_p020_15r_c <- avr_rcor(G1_p020, G5r_p020, 60000, nthreads = 0))
# save.image("result.Rdata")
# st_rca_p020_5qr_c <- system.time(rca_p020_5qr_c <- avr_rcor(G5_p020, Gqr_p020, 60000, nthreads = 0))
# save.image("result.Rdata")
# cat("  rho for p = 20, long m done")

# st_rca_p050_15_c  <- system.time(rca_p050_15_c  <- avr_rcor(G1_p050, G5_p050,  125000, nthreads = 0))
# save.image("result.Rdata")
# st_rca_p050_5q_c  <- system.time(rca_p050_5q_c  <- avr_rcor(G5_p050, Gq_p050,  125000, nthreads = 0))
# save.image("result.Rdata")
# st_rca_p050_5r_c  <- system.time(rca_p050_5r_c  <- avr_rcor(G5_p050, G5r_p050, 125000, nthreads = 0))
# save.image("result.Rdata")
# st_rca_p050_15r_c <- system.time(rca_p050_15r_c <- avr_rcor(G1_p050, G5r_p050, 125000, nthreads = 0))
# save.image("result.Rdata")
# st_rca_p050_5qr_c <- system.time(rca_p050_5qr_c <- avr_rcor(G5_p050, Gqr_p050, 125000, nthreads = 0))
# save.image("result.Rdata")
# cat("  rho for p = 50, long m done")

###########
## Producing summary plots
source("R/plot_fun.R")

pdf("summary_excl_rho.pdf", height = 240 / 25.4, width = 320 / 25.4,
    colormodel = "rgb")
ae_plot_mat("cevo", suffixes = c("1", "5", "q"))
ae_plot_mat("resp", suffixes = c("1", "5", "q"))
ae_plot_mat("flex", suffixes = c("1", "5", "q"))
ae_plot_mat("auto", suffixes = c("1", "5", "q"))
ae_plot_mat("rdif", suffixes = c("15", "1q", "5q"))
ae_plot_mat("rdif", suffixes = c("1r", "5r", "qr"))
ae_plot_mat("rdif", suffixes = c("15r", "1qr", "5qr"))
dev.off()

pdf("summary_rho.pdf", height = 240 / 25.4, width = 240 / 25.4,
    colormodel = "rgb")
ae_plot_mat("rcor", ps = c(5, 20, 50), suffixes = c("15", "1q", "5q"))
ae_plot_mat("rcor", ps = c(5, 20, 50), suffixes = c("1r", "5r", "qr"))
ae_plot_mat("rcor", ps = c(5, 20, 50), suffixes = c("15r", "1qr", "5qr"))
dev.off()

sink()
```
